# DRP1 induces neuroinflammation *via* transcriptional regulation of NF-ĸB

**DOI:** 10.1101/2025.09.02.673863

**Authors:** Yanhao Lai, Rebecca Z. Fan, Harry J. Brown, Said S. Salehe, Ethan K. Tieu, Kim Tieu

## Abstract

Neuroinflammation is a major pathogenic mechanism in neurodegenerative diseases. Understanding the regulation of neuroinflammation is critical to therapeutic development. We report here that dynamin related protein 1 (DRP1), well-recognized for its role in mitochondrial fission, is a transcription factor that regulates neuroinflammation. Using multiple inflammatory models, we provide evidence demonstrating that DRP1, when challenged with pro-inflammatory lipopolysaccharides, translocates from the cytosol to the nucleus, then binds to the promoter region of *Rela* (encoding NF-κB) to activate its gene products and other downstream inflammatory cytokines. Our data also demonstrate the significant role of the proinflammatory lipocalin 2 in the brain. In combination, this study highlights a previously unidentified function of DRP1 in mediating neuroinflammation *via* the NF-κB-lipocalin 2 axis. Through such mechanisms of DRP1, this study also provides potential therapeutic targets for neurodegenerative diseases and other conditions linked to inflammation.

## Introduction

Neuroinflammation is a fundamental innate immune response in the central nervous system involving the activation of glia cells, release of inflammatory mediators (such as cytokines and chemokines), and generation of reactive oxygen and nitrogen species^1^. Emerging evidence indicates the central role of neuroinflammation in neurodegenerative diseases such as Alzheimer’s disease (AD), Parkinson’s disease (PD)^2^. Thus, targeting neuroinflammation represents one of the most promising therapeutic strategies for neurological disorders.

Activation of the nuclear factor kappa B (NF-κB) is a central signaling pathway in inflammation and this family of transcription factors have been intensively investigated as therapeutic targets for a wide range of diseases^3-5^. In mammals, the NF-κB family is composed of five different members: p65 (RelA), RelB, c-Rel, p50 (NF-κB1), p52 (NF-κB2)^5, 6^. Heterodimer of the p65 and p50 subunits is the most abundant form of NF-κB and the most potent activator of gene transcription^6, 7^. NF-κB can be activated by a wide variety of stimuli including bacterial toxins such as lipopolysaccharide (LPS), oxidative stresses, toxic proteins and inflammatory cytokines^6^. Upon activation, the NF-κB subunits assembles into the homo- or hetero-dimerized transcription factor complexes which then translocate into the nucleus and bind to the promoter regions of the target genes, leading to the transcription of pro-inflammatory cytokines^8, 9^. To date, NF-κB is known to regulate the expression of hundreds of different genes^8, 9, 3, 10^, including *LCN2*^11^ Therefore, identifying factors that regulate its expression is highly critical.

*LCN2* encodes the lipocalin 2 protein, also known as neutrophil gelatinase-associated lipocalin (NGAL) and 24p3. Lipocalin 2 is a member of the lipocalin superfamily, a group of secretory proteins that play a variety of physiological roles including bacteriostasis, cell differentiation, iron binding, and immune response^12, 13^. Additionally, lipocalin 2 was reported to act as a pro-inflammatory regulator in activated macrophage induced by NF-κB activation^14^. Although *LCN2* transcript is widely expressed in a wide range of tissues and organs, lipocalin 2 levels are either low or undetectable under basal conditions in the brain^15, 16^. However, under pathological conditions or exposure to inflammatory stimuli, its expression and secretion are drastically increased. Indeed, increased levels of *LCN2* mRNA and lipocalin 2 have been detected in the post-mortem brain tissues of PD and AD patients and their respective animal models^17-21^. Consistent with the role of lipocalin 2 in neuroinflammation, accumulating evidence shows that lipocalin 2 is highly expressed in reactive astrocytes and microglia^22-24^. Collectively, these studies indicate that lipocalin 2 is neurotoxic when upregulated under pathological conditions. However, the mechanism by which dynamin related protein 1 (DRP1) induces neuroinflammation *via* the NF-κB-lipocalin 2 axis has not been reported.

DRP1 is a member of the dynamin GTPase superfamily. Overall, DRP1 is best known for its role in mitochondrial fission^25, 26^. Independent laboratories have demonstrated that partial inhibition of DRP1 function is protective in experimental models of neurodegenerative diseases^26^. We recently reported that transcript and protein levels of DRP1 is highly elevated in the substantia nigra of PD patients^27^, further strengthening the potential involvement of this protein in PD. Consistent with the documented role of DRP1 in mitochondrial fission, the protective mechanism of blocking DRP1 function has been largely attributed to restoring the balance in mitochondrial fission and fusion. However, additional mechanisms are involved. For example, we recently reported that a partial DRP1 deletion improved autophagy flux independent of its mitochondrial fission role^28^. Other investigators have also reported that DRP1 predominantly resides in the cytosol with only approximately 3% cofractionating with the mitochondria^29^. Using live-cell imaging of fluorescently labeled DRP1, a subsequent study corroborated that much of DRP1 appears diffuse in the cytoplasm and only ∼2.5% of the total DRP1 puncta engage in mitochondrial fission^30^. Other laboratories have also reported that DRP1 also colocalizes with other organelles such as lysosomes^31^, endoplasmic reticulum^32^ and peroxisomes^33^. In combination, these studies suggest that DRP1 is a multifunctional protein that exerts its effects beyond mitochondrial division. In this study we provide evidence showing that DRP1 induced neuroinflammation through a transcriptional mechanism. Using LPS to induce microglial activation, we detected the translocation of DRP1 from the cytosol to the nucleus where it binds to the promoter of the *Rela* gene, resulting in the expression of its gene products and other downstream inflammatory cytokines, including *Lcn2*. Compared to other pro-inflammatory genes, *Lcn2* is the most inducible and cell-type dependent. Elevated *Lcn2* levels were also detected in transgenic mice overexpressing human α-synuclein and in old mice (18-24 months). These observations were significantly blunted in mutant mice with heterozygous knockout of DRP1. Collectively, our study identifies DRP1-NF-κB-lipocalin 2 axis in mediating neuroinflammation.

## Results

### Neuroinflammation in multiple mouse models is reduced by a partial DRP1-knockdown

Reduced DRP1 function has been shown to attenuate neuroinflammation in LPS-treated cultured microglia^34, 35^. However, the *in vivo* significance of such observations has not been established and the mechanism by which DRP1 induces neuroinflammation requires further investigation. To address these gaps of knowledge, we used our recently generated mutant mice with a partial DRP1 deficiency (*Dnm1l^+/-^*)^28^. These *Dnm1l^+/-^* mice and their wild-type (WT, *Dnm1l^+/+^*) littermates were injected intraperitoneally (ip.) with a single dose of LPS (5mg/kg)^36, 37^, and their ventral midbrains (VMB) were collected 6h later for RNA analysis. Taking an unbiased approach, we performed NanoString nCounter analysis using the Mouse Neuroinflammation Panel which included 757 target genes covering the core neuroinflammatory pathways and processes. Consistent with the proinflammatory effects of LPS, increased levels of numerous proinflammatory genes in the *Dnm1l^+/+^* mice were detected; however, significant protection was observed in the mutant *Dnm1l^+/-^* littermates (Fig. 1a and Supplementary Table 1). For brevity, only the top 10 differentially expressed genes were illustrated (Fig. 1a), among which, *Lcn2*, was most dramatically increased in LPS-treated WT mice. The normalized mRNA counts of *Lcn2* in LPS-treated *Dnm1l^+/+^* mice was 1510±209.5, whereas that of *Lcn2* in LPS-treated *Dnm1l^+/-^* mice was 543.6±199.9. These results indicate that a partial DRP1 reduction in the *Dnm1l^+/-^* littermates drastically attenuated this upregulation (p=0.0005)(Fig. 1b). No sex difference was detectable; therefore, the results were combined.

**Fig. 1:**
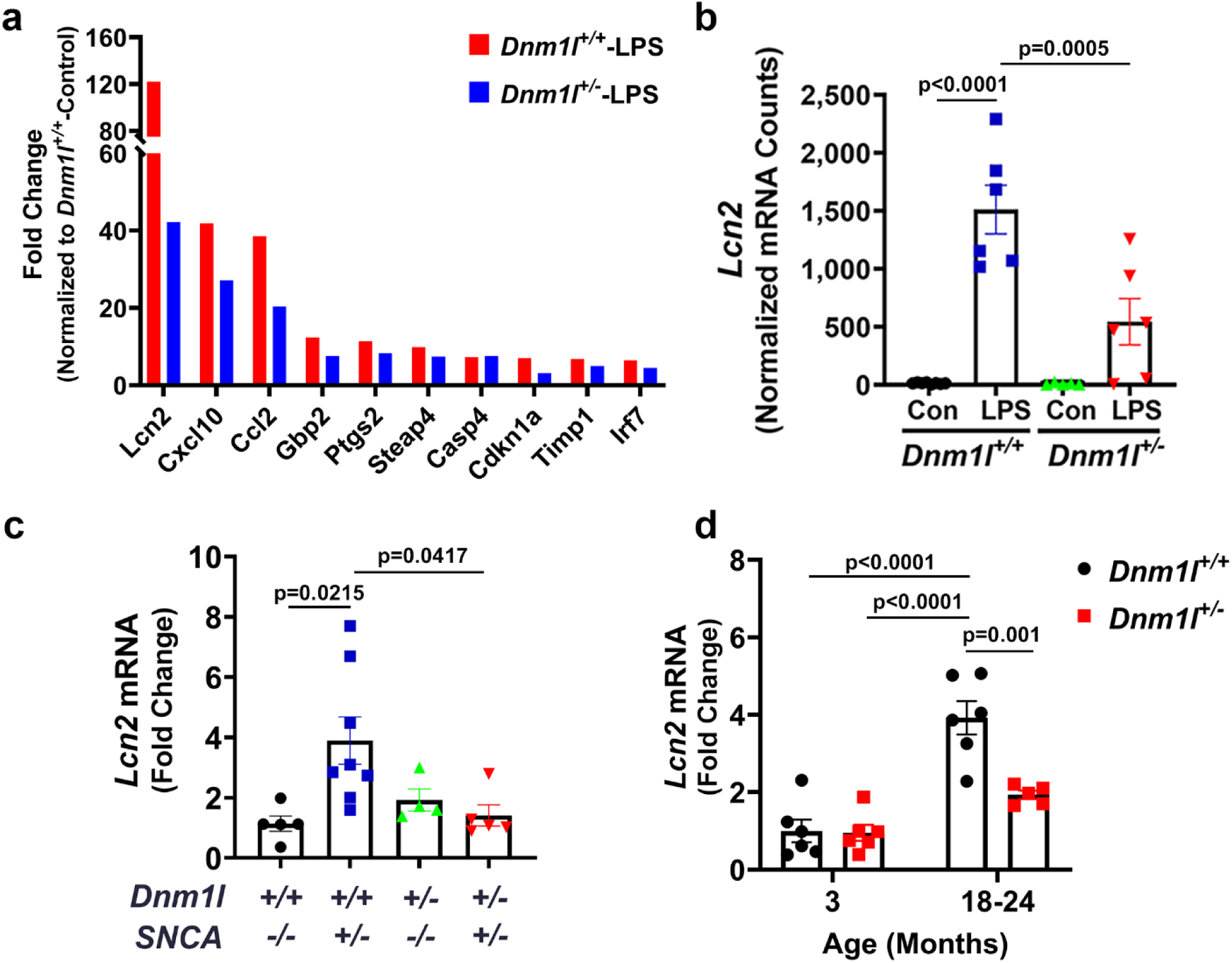
Identification of *Lcn2* as a lead proinflammatory gene target. **a**, 3-month-old *Dnm1l^+/-^* mice and their *Dnm1l^+/+^* littermates were injected (i.p.) with one dose of LPS (5 mg/kg) or saline (control). Ventral midbrains (VMB) were collected 6h later for RNA and subjected to NanoString nCounter gene expression analysis. n=5-8 (2-4F & 3-4M) mice per group. **b**, *Lcn2* levels were quantified for mRNA counts and normalized using *Aars*, *Ccdc127*, *Cnot10*, *Tada2b* and *Xpnpep1* as reference genes. N=5-8 (2-4F & 3-4M) mice per group, two-way ANOVA. **c**, *Lcn2* levels (normalized to *Gapdh*) in the VMB of 6-month-old of mice crossed between *Dnm1l^+/-^* and *SNCA^+/-^,* n=4-8 (2-4F & 1-5M) mice per group, one-way ANOVA. **d**, qPCR analysis of *Lcn2* levels (normalized to *Gapdh*) in the VMB of 3 months and 18-24 months old mice, n=6-7 mice per group (3F & 3-4M), two-way ANOVA. Tukey’s post hoc test was applied following ANOVA analysis for all data.

To investigate whether the effects of DRP1 inhibition against *Lcn2* elevation were specific to LPS exposure, we crossed *Dnm1l^+/-^* mice with transgenic mice overexpressing human α-synuclein. As shown in Fig. 1c, these *SNCA^+/-^* mice expressed 3.9-fold higher *Lcn2* levels than their WT counterparts by the age of 6 months. However, in the double mutant *Dnm1l^+/-^*:*SNCA^+/-^* mice, the levels of *Lcn2* were significantly reduced to the baseline of the control *Dnm1l^+/+^* group (Fig. 1c). Moreover, in a separate aging study, *Lcn2* was upregulation by 3.9-fold in 18-24 months old *Dnm1l^+/+^* mice compared to 3-months old WT mice, but significantly less in their age-matched *Dnm1l^+/-^* littermates (Fig. 1d). Taken together, these data indicate that a partial DRP1 inhibition is effective in reducing the levels of neuroinflammatory genes, especially *Lcn2* in multiple mouse models. Given that aging is a risk factor for chronic neurodegenerative diseases such as PD and AD, these data also suggest the potential involvement of lipocalin 2 in age-related brain disorders. Indeed, increased levels of lipocalin 2 were observed in both PD and AD patients^18, 20^, consistent with the critical role of neuroinflammation in these disorders.

### Reduced proinflammatory gene expression in LPS-treated *Dnm1l^+/-^* mice

To validate the NanoString data and to further investigate whether the alterations of *Lcn2* levels correlate with the levels of proinflammatory genes, we quantified mRNA levels of select proinflammatory genes in the VMB of the LPS-treated mice. These genes include the ones that can be regulated by *Lcn2*^14^, such as *Nlrp3*, *Il1b*, as well as the ones that can control the expression of *Lcn2*^38, 39^, such as *Rela*, *Il6*, *Tnfα*. Our results showed that all the detected genes were significantly upregulated by LPS in *Dnm1l^+/+^* mice but attenuated in their *Dnm1l^+/-^* counterparts (Fig. 2a). These observations were consistent with the altered *Lcn2* levels in these animals (Fig. 1a-b).

**Fig. 2:**
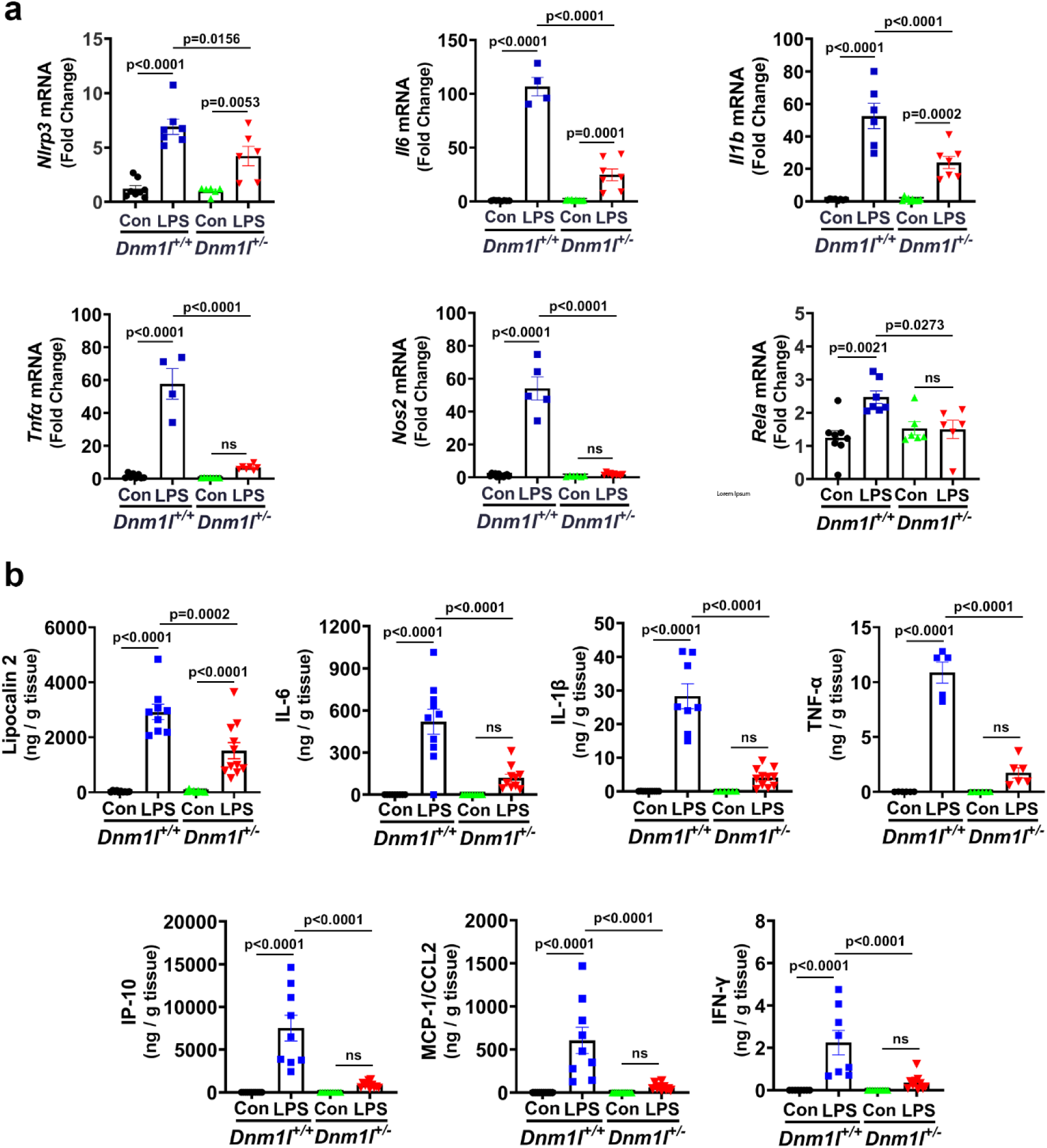
Reduced proinflammatory gene expression in the LPS-treated *Dnm1l^+/-^* mice. **a,b**, LPS (5 mg/kg, i.p.) or saline control was injected to 3-month-old *Dnm1l^+/-^* and *Dnm1l^+/+^* mice and the VMB was collected 6h later for qPCR analysis (**a**) and MSD multiplex immunoassays (**b**). **a**, Relative gene expression of *Nlrp3*, *Il6*, *Il1b*, *Tnfα*, *Nos2* and *Rela* was quantified using *Gapdh* as a reference gene. n=4-10 (2-5F & 1-5M) mice per group, two-way ANOVA. **b**, Data were normalized to total sample protein concentration, n=10-11 (n=5-6F and 5-6M mice), two-way ANOVA. Tukey’s post hoc test was used following ANOVA analysis for all data.

To corroborate the gene expression data obtained from the NanoString and qPCR analysis, we used MSD multiplex immunoassay to examine the protein levels of proinflammatory genes in tissue lysates extracted from the VMB of the LPS-treated *Dnm1l^+/+^* and *Dnm1l^+/−^* mice. Consistent with the alterations observed at the mRNA levels, in addition to lipocalin-2, LPS significantly upregulated other proinflammatory protein levels in the *Dnm1l^+/+^* mice when compared to control *Dnm1l^+/+^* mice (Fig. 2b). However, such increases were drastically attenuated in their *Dnm1l^+/-^* littermates (Fig. 2b). Morphologically, microglia in the *Dnm1l^+/-^* animals also appeared less activated (more ramified structures) as quantified using Sholl analysis of Iba-1 immunostaining (Supplementary Fig. S1).

### Expression of proinflammatory genes and inducibility of *Lcn2* in different cell types

Since the data obtained from the whole VMB tissue (Figs. 1-2) do not provide the resolution of *Lcn2* gene expression with cell-type specificity, and *in vivo Lcn2* expression in different brain cell types has not been evaluated in prior research, we used laser microdissection to remove individual microglia, astrocytes, nigral dopamine (DA) neurons and gamma-aminobutyric acid (GABA) neuron in the substantia nigra of *Dnm1l^+/-^* and *Dnm1l^+/+^* mice injected with LPS. Fig. 3a shows that *Lcn2* was constitutively expressed in all types of cells that were examined. Moreover, no differences were detected in *Lcn2* basal expression level across these four different types of cells (Fig. 3b). Interestingly, although LPS significantly increased the *Lcn2* gene expression in microglia, astrocytes and DA neurons from *Dnm1l^+/+^* mice, no such change was observed in the GABA neurons (Fig. 3a). Consistent with the observation of reduced *Lcn2* level in whole VMB tissue of *Dnm1l^+/-^* mice (Fig. 1b), LPS did not increase the expression level of *Lcn2* in microglia, astrocytes and DA neurons in the *Dnm1l^+/-^* mice when compared to their counterparts isolated from *Dnm1l^+/+^* littermates (Fig. 3a). Furthermore, to compare the inducibility of *Lcn2* in different types of cells upon exposure to LPS in *Dnm1l^+/+^* mice, the *Lcn2* gene expression levels in each cell type were normalized to the *Lcn2* level detected in microglia from *Dnm1l^+/+^* mice without LPS treatment. The results showed that *Lcn2* in astrocytes and microglia were more inducible than its counterpart in DA neurons upon exposure to LPS (Fig. 3b). Our results suggest that the observed LPS-induced upregulation of *Lcn2* as seen in Figure 1 mainly arises from glia cells (microglia and astrocytes) and to a lesser extent from DA neurons, but not from GABA neurons.

**Fig. 3:**
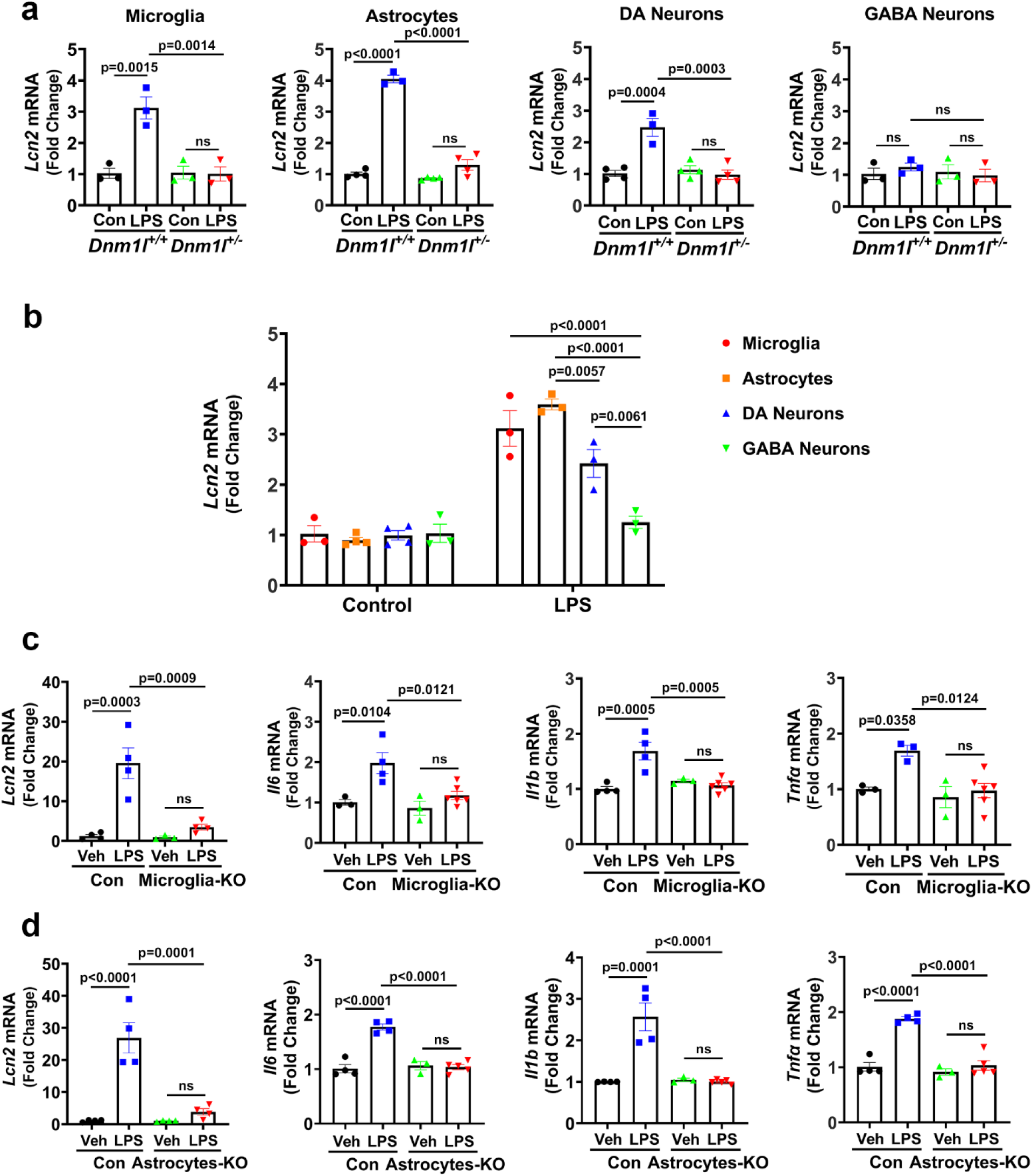
Expression of proinflammatory genes and inducibility of *Lcn2* in different cell types in *Dnm1l* mutant mice. **a,b**, *Dnm1l^+/-^* mice (3-month-old) were treated with LPS (5 mg/kg, i.p.) or saline control and 6h later brains were snap-frozen, VMB was coronally sectioned (10 µm) and immunostained for microglia, astrocytes, DA and GABA neurons using antibodies against IBA1, GFAP, TH and GAD67, respectively. Different cell types were individually captured using LMD6 (Leica). **a**, Micro-dissected cells (40 cells / each type) from the nigra of each animal were processed for cDNA and pre-amplified using “Smart-seq2” method, followed by qPCR analysis of *Lcn2* using *Gapdh* as a reference gene. N=3-4 (2F & 1-2M) mice per group. **b**, Inducibility of *Lcn2* upon LPS exposure in different cell types was quantified in WT (*Dnm1l^+/+^*) mice. **c,d**, Tamoxifen injected 3-month old *Cx3Cr1-Cre^ERT+/^*^-^:*Dnm1l-Loxp*^+/-^ (denoted as “Microglia-KO”), *Aldh1L1^Cre/ERT2+/^*^-^:*Dnm1l-Loxp*^+/-^ (Astrocytes-KO), and *Dnm1l-Loxp*^+/-^ (control) mice were injected with LPS (5 mg/kg, i.p.) and brain tissue was collected 6h later and prepared for laser microdissection to capture individual cell type as described above. “Smart-seq2” method was used to pre-amplify cDNA of 40 microglia and astrocytes from the nigra of each animal. **c**, qPCR analysis of *Lcn2*, *Il6*, *Il1b*, and *Tnfα* (normalized to *Gapdh*) of microglia. N=3-6 (1-2 F & 1-4 M) mice per group. **d**, qPCR analysis of *Lcn2*, *Il6*, *Il1b*, and *Tnfα* (normalized to *Gapdh*) of astrocytes. N=3-5 (1-2 F & 1-3 M) mice per group. Two-way ANOVA, followed by Tukey’s post hoc test was used for all data analysis.

To complement the results obtained from the microdissected cells in the heterozygous global DRP1-KO (*Dnm1l^+/-^*) mouse model (Figs. 3a-b), we further used mice with conditional knockdown of DRP1 in microglia or astrocytes by crossing our recently generated floxed-*Dnm1l* mice with *Cx3Cr1^CreERT2^* or *Aldh1L1^CreERT2^* mice, respectively. To confirm the tamoxifen induced *Dnm1l* gene knockdown in the microglia of “Microglia-KO” mice and astrocytes of “Astrocyte-KO” mice, microglia and astrocytes were isolated by laser microdissection and subjected to smart-seq2 linked qPCR analysis 3 weeks post-tamoxifen injection. Approximately 32% and 44% reduction in *Dnm1l* levels were observed in the “Microglia-KO” mice and “Astrocyte-KO” mice, respectively (Supplementary Fig. S2). Such reduction was not observed in other cell types that we did not target to knockout *Dnm1l* in these mutant mice (Supplementary Fig. S2), indicating the loss of *Dnm1l* was cell type specific as intended. In line with the observations in global DRP1-KO mouse model, LPS significantly increased the expression levels of proinflammatory genes, such as *Lcn2*, *Il6*, *Il1b*, and *Tnfα* in both microglia and astrocytes of control (*Dnm1l-Loxp*^+/-^) mice in comparison to those from their littermates without LPS treatment (Fig. 3c-d). However, such upregulation was abolished in the mutant mice with heterozygous deletion of *Dnm1l* in these cell types (Fig. 3c-d). These results suggest that the reduction of both *Dnm1l* in microglia and astrocytes plays important roles in combating LPS-induced neuroinflammation.

### DRP1 inhibition prevented neuroinflammation *via* the NF-ĸB pathway

Based on our results showing that heterozygous DRP1-KO was protective against neuroinflammation-induced by LPS and based on the known function of DRP1 in mitochondrial function, we initially asked whether the protective mechanism was mediated by preventing mitochondrial dysfunction induced by LPS. To this end, we first evaluated whether there was mitochondrial impairment in microglia in the animal model that we used. We injected adult mice with LPS, then isolated and enriched microglia for the measurement of mitochondrial respiration. No mitochondrial impairment was detectable with this regimen (Supplementary Fig. S3a and S3b). These data suggest the reduced neuroinflammation observed in Figs. 1-3 was not mediated through attenuating mitochondrial deficits induced by LPS. we treated primary microglia from *Dnm1l^+/+^* and *Dnm1l^+/-^* with LPS to induce neuroinflammation.

To further investigate the mechanism by which DRP1 inhibition (*Dnm1l^+/-^*) reduces LPS-induced neuroinflammation and microglia activation, we treated primary microglia with LPS for 6h (the same timepoint we used for the *in vivo* study), followed by chromatin immunoprecipitation (ChIP) assay to determine whether *Lcn2* is transcriptionally regulated by NF-ĸB as previously reported^14^. Consistent with the presence of ĸB site at the promoter region of *Lcn2* gene (Fig. 4a), we detected an enrichment of NF-ĸB p65 at the promoter of *Lcn2* gene in WT microglia treated with LPS (Fig. 4b). However, in microglia with heterozygous DRP1-KO, this binding was significantly reduced (p=0.0012) (Fig. 4b), indicating DRP1 inhibition prevented neuroinflammation by reducing NF-ĸB recruitment to its target proinflammatory gene (*Lcn2)*. Next, we examined the levels of proinflammatory cytokines in the conditioned medium of microglia treated with LPS in the absence or presence of an NF-ĸB inhibitor, Bay11-7082^40^. This compound drastically inhibited the release of proinflammatory cytokines (p<0.0001) in the LPS-treated *Dnm1l^+/+^* microglia (Fig. 4c). The effects of this pharmacological inhibition were further corroborated genetically using *Dnm1l^+/-^* microglia (Fig. 4c). These *in vitro* results further support the *in vivo* data showing reduced levels of proinflammatory proteins in the VMB of LPS-treated *Dnm1l^+/-^* mice compared to their WT littermates (Fig. 2b). In combination, these observations are consistent with the role of NF-ĸB as a major regulator of inflammation and they support our hypothesis that DRP1 plays a critical role in neuroinflammation *via* the NF-ĸB pathway.

**Fig. 4:**
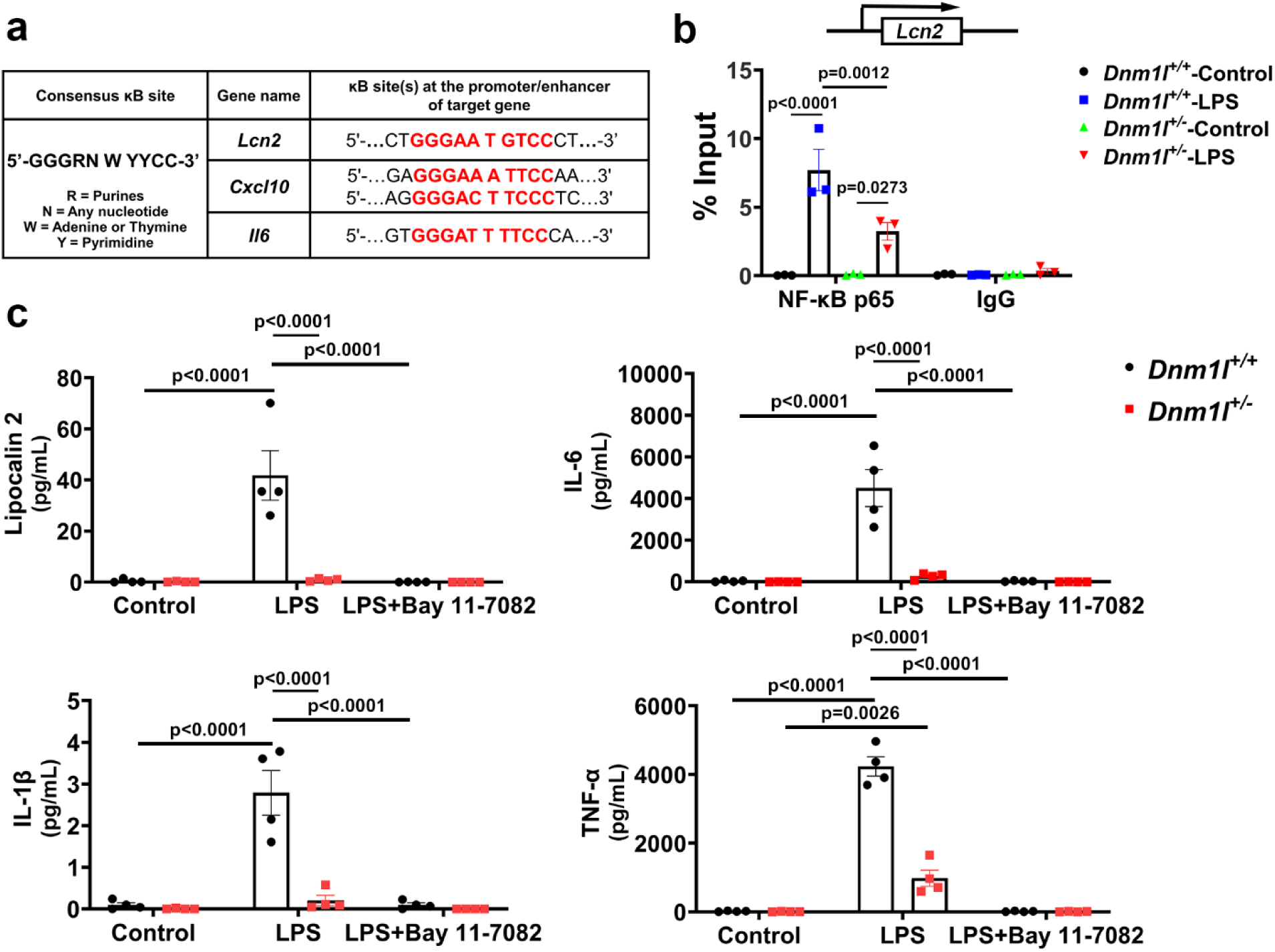
DRP1 inhibition prevents neuroinflammation *via* NF-ĸB pathway. **a**, Consensus NF-ĸB binding sequence (ĸB site) as well as its binding site(s) at the promoter of *Lcn2*, *Cxcl10* and *Il6*. **b**, *Dnm1l^+/-^* and *Dnm1l^+/+^* primary microglia were treated with 100 ng/ml LPS for 6h then subjected to ChIP assay using rabbit anti-NF-ĸB p65 antibodies or rabbit IgG antibodies. The “NF-ĸB p65” or “IgG” bound DNA were recovered for qPCR using primers that targeted the ĸB site at the promoter of *Lcn2* gene. Data represent mean±SEM from three independent experiments. Three-way ANOVA, Tukey’s post hoc analysis. **c**, *Dnm1l^+/+^* and *Dnm1l^+/-^* microglia were treated with LPS (100 ng/ml, 6h) in the absence or presence of Bay11-7082 (5 µM), an NF-ĸB inhibitor. At the end of treatment, the conditioned medium was collected for the detection of secreted cytokines using MSD multiplex immunoassay. Data represent mean±SEM from four independent experiments. Two-way ANOVA, Tukey’s post hoc analysis.

To determine the effects of LPS on mitochondria in our primary microglia model, we performed time-course studies. At the 6h time-point, which induced neuroinflammation, we did not detect mitochondrial fragmentation (Supplementary Fig. S3c and Sd) nor dysfunction (Supplementary Fig. S3e-h) in WT microglia at this time point or longer (24h). However, at an earlier point (2h), we did detect shorter mitochondria equally in both *Dnm1l^+/+^* and *Dnm1l^+/-^* microglia (Supplementary Fig. S3d). These data indicate that first, LPS can activate microglia without impairing mitochondria. Second, the effects of LPS on mitochondrial morphology at an early time point is reversible with no detectable functional deficits. Third, it further supports that a partial DRP-KO protects against neuroinflammation *via* a mechanism independent of mitochondrial dysfunction.

### *Lcn2*-KO reduces proinflammatory cytokines released from microglia

Lipocalin 2 is an emerging prominent inducer of neuroinflammation^12, 14, 20^. Our data shown that DRP1 deficiency is effective in reducing NF-κB and lipocalin 2 levels, indicating the involvement of the DRP1-NF-κB-lipocalin 2 axis in neuroinflammation. To determine whether the levels of *Lcn2* expression correlate with proinflammatory cytokines, primary microglia from *Lcn2^+/+^*, *Lcn2^+/-^*, and *Lcn2^-/-^* were treated with LPS, and proinflammatory cytokines in the medium was analyzed using multiplex ELISA immunoassay. As shown in Figure 5A, basal secretion of lipocalin 2 from *Lcn2^+/+^* microglia was 5.8-fold higher than that of *Lcn2^+/-^* microglia (p=0.011), and not detectable from the *Lcn2^-/-^* group. Such genotypic differences were not observed in the basal levels of IL-6, IL-1β, CXCL-10/IP10, MCP-1/CCL2, and IFN-γ (Fig. 5a), supporting the specific effects of *Lcn2* deletion. As expected, when treated with LPS, the secretion of proinflammatory cytokines was increased in *Lcn2^+/+^* microglia (Fig. 5a). *Lcn2* deletion reduced the levels of proinflammatory cytokines in a gene-dose dependent manner (Fig. 5a). Although a partial-knockdown of *Lcn2* (*Lcn2^+/-^*) was sufficient to reduce lipocalin 2 (p<0.001), IL-6 (p=0.0009) and IL-1β (p=0.0015) (Fig. 5a), a complete *Lcn2* deletion (*Lcn2^-/-^*) was required for a more widespread reducing additional cytokines (Fig. 5a).

**Fig. 5:**
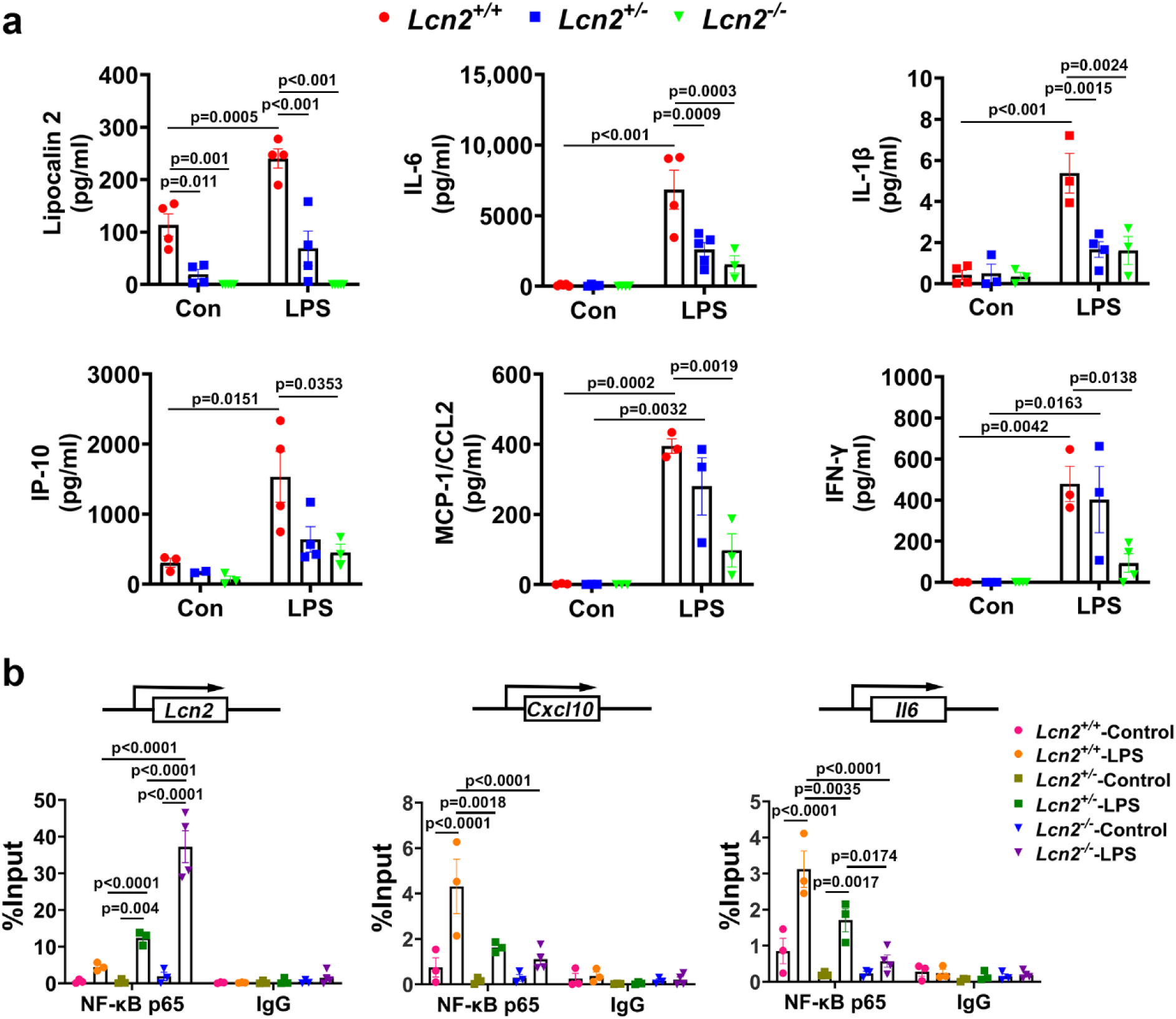
*Lcn2*-KO reduces proinflammatory cytokines released from microglia upon LPS induction. **a**, Microglia isolated from P0-P1 pups using CD11b immunomagnetic beads were grown in 96-well plates and treated with LPS (100 ng/ml) for 6h, the conditioned media were then collected and analyzed with MSD U-PLEX Custom Biomarker multiplex assay. N=3-5 independent experiments for each analyte with 6 replicates per experiment, two-way ANOVA followed by Tukey’s post hoc analysis. **b**, *Lcn2^+/+^*, *Lcn2^+/-^*, and *Lcn2^-/-^* microglia were subjected to LPS treatment (100 ng/ml, 6h) followed by ChIP assay as described. The “NF-ĸB p65” or “IgG” bound DNA were recovered for qPCR using primers that targeted the ĸB site(s) at the promoter of *Lcn2, Cxcl10* and *Il6*, respectively. Ct values were recorded and used for data analysis. N=3 independent experiments, three-way ANOVA followed by Tukey’s post hoc analysis.

Given that DRP1-KO prevented neuroinflammation by reducing NF-ĸB recruitment to its target proinflammatory genes, we asked whether reduced neuroinflammation in *Lcn2*-KO microglia upon LPS induction also resulted from decreased NF-ĸB recruitment. To address this question, we conducted ChIP assay to examine the recruitment of NF-ĸB p65 to the promoter of *Lcn2* gene using *Lcn2^+/+^*, *Lcn2^+/-^*, and *Lcn2^-/-^* microglia. Surprisingly, we found that the recruitment of NF-ĸB p65 to the promoter of *Lcn2* gene was inversely correlated to the *Lcn2* gene expression levels (Fig. 5b). We hypothesized that more NF-ĸB p65 was recruited to the promoter of *Lcn2* gene in *Lcn2*-KO microglia as a compensatory attempt to increase the expression of this gene. However, due to the deletion of exons 2-5 of the *Lcn2* gene in the *Lcn2*-KO microglia, the enrichment of NF-ĸB p65 failed to induce *Lcn2* gene expression, which further led to the accumulation and hijacking of NF-ĸB p65 at the promoter of *Lcn2*. If this were true, such hijacking of NF-ĸB p65 at the promoter of *Lcn2* would lead to reduced recruitment of NF-ĸB p65 to other target genes. To test this hypothesis, we further measured the recruitment of NF-ĸB p65 to other proinflammatory genes that were transcriptional controlled by NF-ĸB, such as *Cxcl10* and *Il6*, using ChIP. Indeed, NF-ĸB p65 binding to the ĸB sites at the promoter of these genes (Fig. 4a) was decreased in *Lcn2*-KO microglia when compared to *Lcn2*-WT microglia (Fig. 5b). These results confirmed that the reduced neuroinflammation in *Lcn2*-KO microglia was due to, in least in part, the hijacking of NF-ĸB at the promoter of *Lcn2*, which in turn prevented its recruitment to other target proinflammatory genes and ultimately led to reduced neuroinflammation.

### LPS induces upregulation and nuclear translocation of DRP1

Based on our observations that DRP1-KO reduced the expression of proinflammatory genes (Figs. 1-2), we hypothesized that the reduced binding of NF-ĸB p65 to the promoter of *Lcn2* was due to lower expression of *Rela*, the gene that encodes NF-ĸB p65. Indeed, qPCR analysis of this gene showed lower expression of *Rela* in the *Dnm1l^+/-^* mice as compared to the WT littermate group (Fig. 2a). Moreover, it has been shown that nuclear DRP1 is highly expressed in lung adenocarcinomas, which might contribute to drug resistance during hypoxia that mediates poor prognosis^41^. DRP1 is a well-documented cytosolic protein^29^, its nuclear expression suggests that DRP1 could translocate to the nucleus under certain circumstances. In combination, such prior research prompted us to formulate the novel hypothesis that DRP1 could transcriptionally regulate the expression of proinflammatory genes. To test this hypothesis, we performed immunofluorescence, followed by Imaris 3D rendering images to quantify nuclear translocation of DRP1 (red) and those that remain in the cytoplasm (yellow) in primary microglia and mice treated with LPS (Fig. 6a-d). In the vehicle control group, we detected some basal nuclear localization of DRP1. This is interesting because to our knowledge, this localization has not been previously reported, and it raises the question of why DRP1 is in the nucleus. More importantly, when microglia were activated, levels of DRP1 in the nucleus (DRP1^Nuc^) increased by 3.8-fold in the LPS-treated primary microglia (p=0.0084) (Fig. 6a and b). Interestingly, the levels of DRP1 in the cytoplasm (DRP1^Cyto^) also increased by 2.3-fold in these microglia (p=0.0001). Consistent with these *in vitro* data, DRP1^Nuc^ and DRP1^Cyto^ levels in the nigral microglia of mice treated with LPS increased by 3-fold (p=0.0009) and 1.5-fold (p=0.0479), respectively, when compared to those in vehicle-treated animals (Fig. 6c and d). These results indicate that there was a drastic translocation of DRP1 into the nucleus as well as an overall upregulation of DRP1 when microglia were activated.

**Fig. 6:**
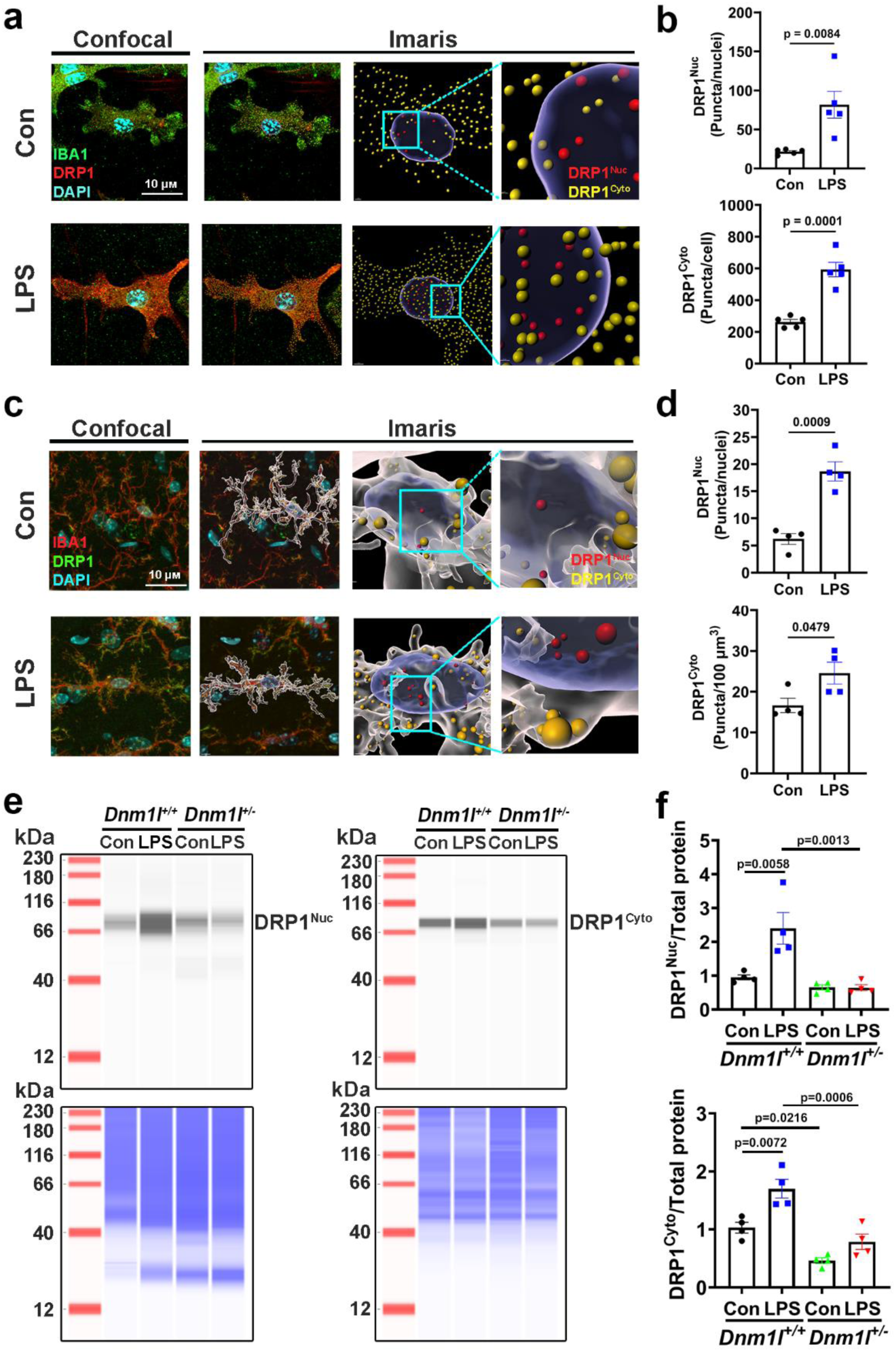
LPS induces upregulation and nuclear translocation of DRP1. **a,b**, Primary microglia cultured from *Dnm1l^+/+^* pups were grown on chamber slides and treated with LPS (100 ng/ml) or vehicle for 6h. **a**, Representative confocal images of IBA1 and DRP1 immunofluorescence, followed by Imaris 3D rendering. **b**, Nuclear DRP1 (red) and cytoplasmic DRP1 (yellow) were quantified using Imaris. N=5 experiments with 14-17 cells/experiment and analyzed using t-tests. **c,d**, 3-month old *Dnm1l^+/+^* mice were injected i.p. with a single dose of LPS (5 mg/kg) or saline. **c**, Midbrain sections were immunostained for SNpc microglia and DRP1, followed by 3D rendering of the nigral microglia and DRP1. **d**, Nuclear and cytosolic DRP1 puncta were quantified using Imaris, N=4 (2F & 2M) with 10-15 cells/animal, t-tests. **e,f**, *Dnm1l^+/+^* and *Dnm1l^+/-^* primary microglia were treated with 100 ng/ml LPS for 6h followed by nuclear and cytoplasmic fractionation. **e**, Immunoblotting of DRP1 in the nuclear and cytoplasmic fractions (top panels). Total proteins per lane (bottom panels) were used as loading control and quantified for the levels of DRP1^Nuc^ and DRP1^Cyto^ in **f**. N=4 independent experiments, two-way ANOVA followed by Tukey post-hoc test.

To further quantitatively measure the nuclear enrichment of DRP1 upon the LPS exposure, capillary immunoblotting was performed to quantify the levels of DRP1 in the nuclear and cytosolic fractions of *Dnm1l^+/+^* and *Dnm1l^+/-^* microglia with or without LPS treatment. To ensure the nuclear fractions were not contaminated with the cytoplasmic fractions and *vice versa*, the nuclear fraction was probed for GAPDH (a cytoplasmic marker) and the cytoplasmic fraction was probed for Histone H3 (a nuclear marker). The results show no cross contamination in these samples (Supplementary Fig. S4b). We then proceed to assessing nuclear translocation of DRP1 upon LPS exposure. Consistent with the immunofluorescence data (Fig. 6a-d), capillary immunoblotting shows the levels of DRP1^Nuc^ and DRP1^Cyto^ were 2.5-fold (p=0.0058) and 1.7-fold (p=0.0072) higher, respectively, in the LPS-treated *Dnm1l^+/+^* microglia (Fig. 6e-f). In contrast, such increases were not observed in *Dnm1l^+/-^* microglia treated with LPS (Fig. 6e-f). Collectively, these results confirm the nuclear enrichment of DRP1 in the WT microglia but not in the heterozygous DRP1-KO microglia after the LPS exposure.

### DRP1 is recruited to the promoter region of *Rela* gene upon LPS exposure

Given that *Rela* levels were lower in *Dnm1l^+/-^* mice when compared to their *Dnm1l^+/+^* littermates after LPS treatment (Fig. 2a), we reasoned that *Rela* might be transcriptionally regulated by DRP1. To test this hypothesis, we further conducted ChIP assay using WT microglia to examine whether DRP1 could be recruited to the promoter of *Rela* after treatment with LPS. However, because DRP1 was not a canonical transcription factor whose binding consensus motif was previously identified, we designed 7 pairs of primers (P1-P7) that could detect the enrichment of DRP1 spanning ∼2000 bp of the promoter region of *Rela* gene (Fig. 7a and Supplementary Table 2). Strikingly, we found that the enrichment of DRP1 was significantly elevated at a 255 bp region that could be amplified by P7 in LPS-treated microglia when compared to control microglia (p<0.0001) (Fig. 7b). Conversely, such enrichment was not observed in IP using IgG (Fig. 7b). In addition, we did not detect any enrichment of DRP1 to other examined regions that could be amplified by P1-P6 (Fig. 7). These results indicate that DRP1 was recruited to the promoter region that was in proximity to the transcription start site of *Rela* gene upon LPS exposure.

**Fig. 7:**
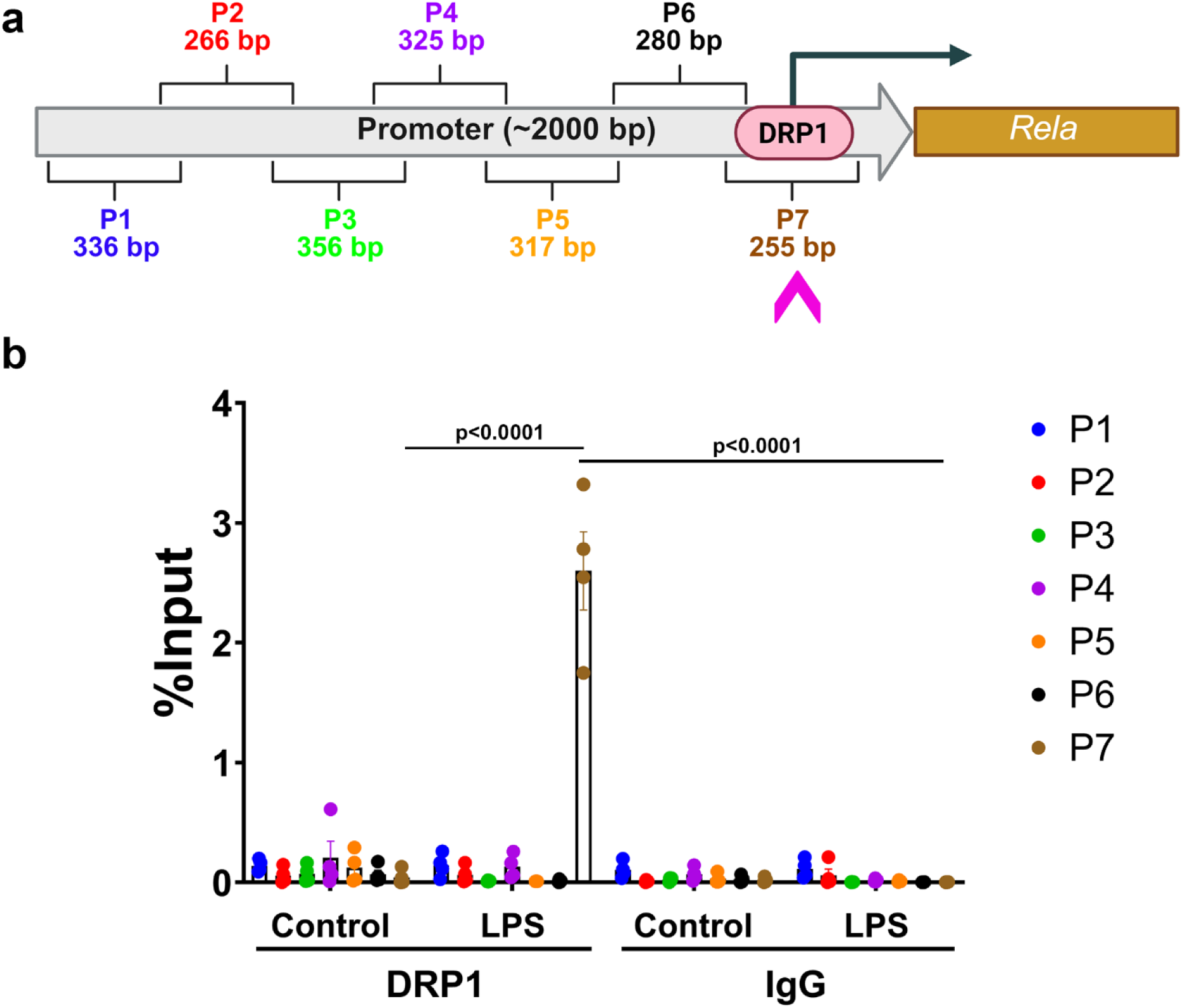
Enrichment of DRP1 at the promoter region of *Rela* gene upon LPS exposure. **a**, Schematic presentation showing the ∼2000 bp promoter region of the *Rela* gene that can be targeted by seven designed primers (P1-7) and where DRP1 was recruited to the ∼255 bp region proximity to the transcription site of *Rela* after LPS exposure. **b**, Primary WT microglia were treated with 100 ng/ml LPS or vehicle control for 6h. ChIP assay was performed to detect the enrichment of DRP1 at the promoter region of *Rela*. All ‘% input’ were obtained from four independent experiments and expressed as mean±SEM, two-way ANOVA with Tukey’s multiple comparison post hoc analysis.

## Discussion

Neuroinflammation is the pathological cornerstone of neurodegenerative diseases. Such negative impact of neuroinflammation underscores the need to learn more about how this process is regulated in order to identify novel therapeutic targets. To date, NF-κB is well-established as a major signaling pathway involved in the regulation of inflammatory responses and its activation has been linked to numerous disorders, including neurodegenerative diseases^9, 42^. As a “master regulator” of inflammation, NF-ĸB regulates numerous pro-inflammatory genes^3, 10^. For example, NF-κB plays a critical role in the formation of NOD-like receptor protein 3 (NLRP3) inflammasome by transcriptionally regulating the expression of its components NLRP3, Pro-IL-1β and Pro-IL-18, pro-caspase-1, and apoptosis associated speck-like protein containing a CARD (ASC)^43^. Once assembled and activated, NLRP3 inflammasome generates mature and proinflammatory cytokines^44^. NF-ĸB-NLRP3 inflammasome represents a major inflammatory pathway.

In the present study we provide evidence showing that NF-ĸB is transcriptionally regulated by DRP1. We demonstrate that DRP1, a well-documented cytosolic protein, was also present in the nucleus at low levels under normal physiological condition. However, upon being challenged with the pro-inflammatory cue (LPS), a significant portion of DRP1 translocated into the nucleus, and subsequently bound to the promoter region of NF-ĸB to activate the expression of proinflammatory genes such as *Rela*, which in turn, upregulated the expression of *Nlrp3*, additional inflammasome related genes, and other proinflammatory genes including *Lcn2,* which has emerged as a critical player in neuroinflammation^11, 13, 45^. The role of lipocalin 2 in NF-κB-mediated activation of the NLRP3 inflammasome was initially investigated in experimental models of heart failure, in which lipocalin 2 induced NLRP3 inflammasome activation via the release of the danger-associated molecular pattern (DAMP) protein and subsequent TLR4-dependent signaling^46^. Recent evidence suggests that lipocalin 2 can directly upregulate the NLRP3 inflammasome complex in macrophages by NF-κB activation in response to LPS^14^. Our study show that in a gene-dose dependent manner, *Lcn2* deletion strongly suppressed the production and secretion of proinflammatory cytokines in LPS-stimulated primary microglia. Interestingly, this anti-inflammatory mechanism was mediated by, at least in part, hijacking of NF-κB at the promoter of *Lcn2*. We hypothesized that the accumulation of NF-ĸB p65 at the promoter of the *Lcn2* gene is a compensatory attempt for the loss of *Lcn2* function. In doing so, NF-ĸB p65 was hijacked at the promoter of *Lcn2*, resulting in reduced recruitment of NF-ĸB p65 to other target genes. Combined with other prior research, our data indicate that in addition to putting a brake on NLRP3 activation, a loss of *Lcn2* function also reduces the availability of NF-κB to activate other pro-inflammatory genes. These results highlight the role of lipocalin 2 in NF-κB-NLRP3 inflammasome signaling pathway.

We detected elevated levels of *Lcn2* in not only LPS models but also transgenic mice over expressing α-synuclein and 18-24 months old mice, demonstrating the involvement of lipocalin 2 in neuroinflammation is most likely a common pathological alteration. However, the most robust increase was detected in the LPS models that we used, most likely due to the temporal changes of a more acute induction of inflammation by LPS. To keep the study focused and for technical feasibility, we used LPS to induce neuroinflammation. The LPS regimen that we selected for the *in vivo* and *in vitro* studies were a balance between high levels of cytokines and no detectable mitochondrial dysfunction. This is an important consideration as a previous study has shown that in primary rodent microglia, mitochondrial impairment and fragmentation induced by a combination of LPS and nigericin for 24h are required for the induction of neuroinflammation ^34^. Although this is an important mechanism, we sought to address the question of whether DRP1 could induce neuroinflammation in the absence of such overt mitochondrial abnormalities. The LPS *in vivo* mouse model that we used was partly based on previous studies^37, 47, 48^. In mice treated with 5mg/kg LPS i.p. for 6h, the levels of pro-inflammatory cytokines in the VMB were substantially increased (Figs. 1-2). However, this treatment did not impair mitochondrial function in microglia (Supplementary Fig. S3a-b). Similarly, in primary microglia treated with LPS for 6h, significant neuroinflammation was detected (Fig. 4c) without alterations in mitochondrial morphology and function (Supplementary Fig. S3c-e and S3g). Although mitochondrial fragmentation was detectable at 2h post LPS treatment without mitochondrial dysfunction, such morphological change was reversible to normal phenotype by 6h. These results indicate that by using neuroinflammatory models without mitochondrial dysfunction, we demonstrate that DRP1 induces neuroinflammation *via* a mechanism independent of mitochondria as illustrated in Fig. 8. In addition to activating neuroinflammation, lipocalin 2 also impairs autophagy flux^46, 49^. Since neuroinflammation induced by LPS also impairs autophagy^50-52^, the combination of lipocalin 2 and LPS would exacerbate each other’s negative impact on autophagy. One significant implication of this effect is its potential link to the accumulation of toxic proteins (such as α-synuclein, tau and β-amyloid) and damaged organelles (mitochondria) as seen in neurodegenerative diseases. Relevant to this study, the turnover of DRP1 is also mediated by autophagy^53^. We detected increased levels of DRP1 in microglia exposed to LPS (Fig. 6). Such an increase may further elevate the pro-inflammatory effects of DRP1, resulting in a vicious cycle of impacted autophagy alterations and neuroinflammation, the two major pathogenic mechanisms in neurodegenerative disorders.

**Fig. 8:**
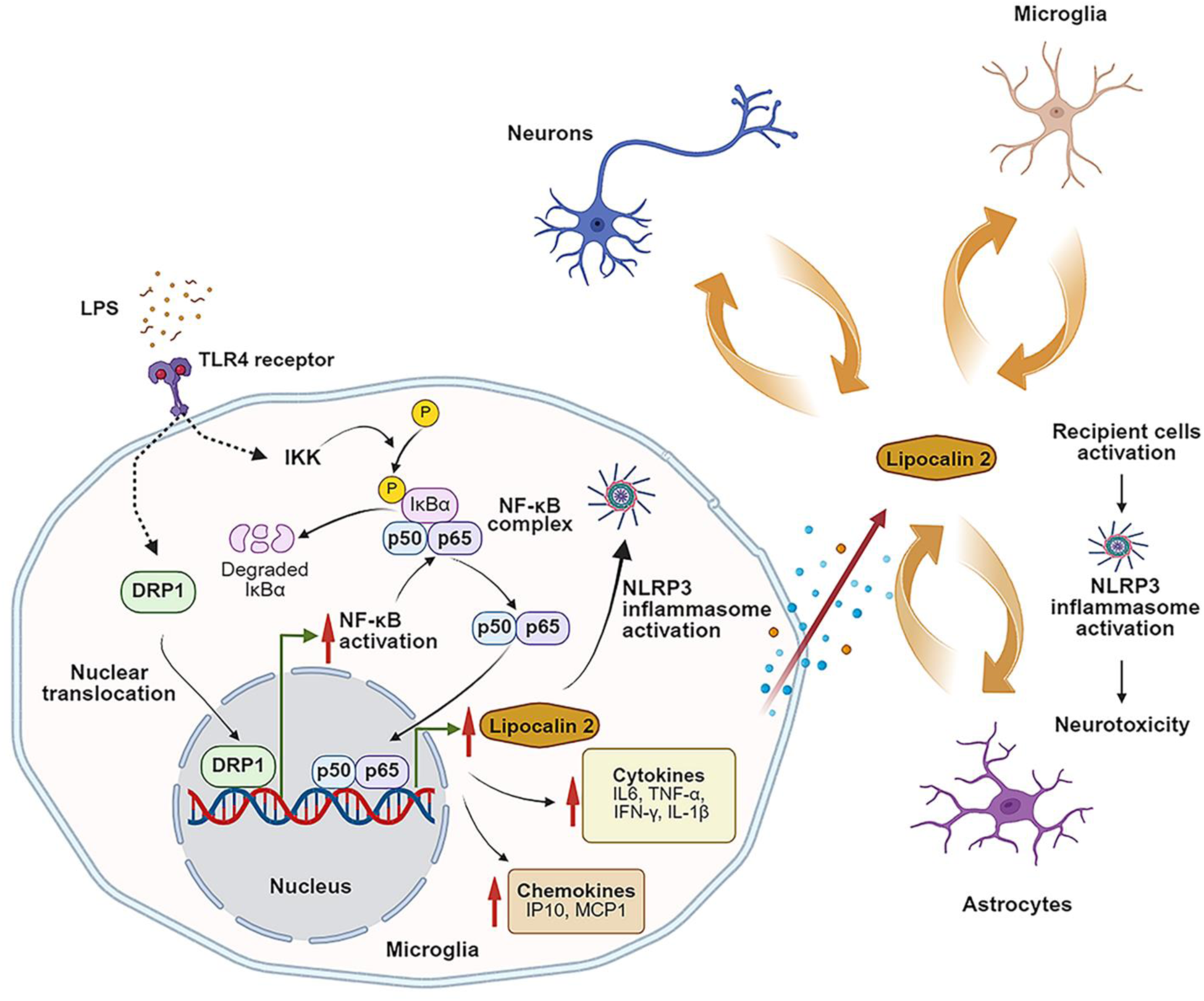
Diagram of how DRP1 induces neuroinflammation *via* the NF-κB-lipocalin 2 axis. When challenged with pro-inflammatory cues such as LPS, DRP1 translocates from the cytosol to the nucleus where it binds to the promoter region of *Rela* to activate its expression and other downstream inflammatory cytokines that can ultimately lead to neurotoxicity. Among the pro-inflammatory genes, *Lcn2* (which encodes lipocalin 2) is the most inducible and cell-type dependent. Once secreted, lipocalin 2 induces neuroinflammation in the neighboring cells.

To investigate the role of DRP1 in neuroinflammation, we used genetic rather than pharmacological approaches to reduce DRP1 function due to the concerns of the off-target effects and other limitations of DRP1 inhibitors. For example, mitochondrial division inhibitor (mdivi-1) is reported to have an off-target effect by reversibly inhibiting of complex I^54^. Dynasore is not a specific DRP1 inhibitor because it also blocks dynamin 1 and dynamin 2^55^. More recently, through virtual and in silico screenings, Drpitor1 was identified as a DRP1 inhibitor. This molecule has been shown to have antitumor activity most likely through DNA intercalation and topisomerase II inhibition^56^. P110 is a peptide-based inhibitor designed to interfere with the interaction of DRP1 with FIS1, a mitochondrial outer membrane fission protein involved in DRP1 recruitment to mitochondria^57^. P110 also partially inhibits the GTPase activity of DRP1^57^. One limitation of using P110 is its poor oral bioavailability due to the unfavorable physiological environment (pH, enzymes) and physical barriers in the gastrointestinal tract. Indeed, in animal studies, P110-TAT is delivered through subcutaneous mini-osmotic pumps^58, 59^ to bypass the gastric issue. In studies where aging animals will be needed, P110 is not feasible for long-term treatment. However, this peptide would be a great tool for *in vitro* or acute *in vivo* studies to complement the heterozygous DRP1-KO models in future studies.

In summary, this study demonstrates that DRP1 is a transcription factor that regulates the expression of proinflammatory genes such as *Rela*, which in turn, upregulates the expression of *Lcn2* and other proinflammatory genes. Since its discovery almost 40 years ago, the role of NF-ĸB in inflammation has been extensively studied. It is clear now that as a “master regulator” of inflammation, NF-ĸB regulates numerous pro-and anti-inflammatory genes.^3^ The complex function of NF-ĸB is also cell type dependent. In neurons, this protein is critical to synaptic plasticity and neuronal survival.^4^ For PD, blocking NF-ĸB reduces neuroprotection and *vice versa* for activating NF-ĸB.^4^ In contrast, as discussed, blocking *Lcn2* has been reported so far to not only reduce neuroinflammation but also increase neuroprotection. Because *Lcn2* is downstream of NF-ĸB, it may not have as many targets as NF-ĸB. Therefore, *Lcn2* may be a more specific target against neuroinflammation. By regulating the NF-κB-lipocalin 2 axis, a highly critical pathway to inflammation, DRP1 is most likely involved in a wide range of diseases.

## RESOURCE AVAILABILITY

### Lead contact

Further information and requests for resources and reagents should be directed to and will be fulfilled by the lead contact, Dr. Kim Tieu.

### Materials availability

Proprietary material is available upon request from the authors.

### Data and code availability

- No new code was required for this study.
- Any additional information required to reanalyze the data reported in this paper is available from the lead contact upon request.

## ACKNOWLEDGMENTS

This study was supported in part by the National Institute of Environmental Health Sciences (NIEHS) of the National Institutes of Health (NIH) under Award Number R35ES030523 (to KT). The content is solely the responsibility of the authors and does not necessarily represent the official views of the NIH. Graphic abstract was created with BioRender.com.

## AUTHORCONTRIBUTIONS

KT, YL and RF contributed to the study conception and design. Experiments and data analysis were performed by YL, RF, HB, SS. and ET. YL and KT were involved in drafting and revision of the manuscript. All the other authors commented on previous versions of the manuscript and approved of the final version.

## Methods

### Animal models and LPS treatment

All mice in this study were bred, maintained, and characterized at the animal care facility of Florida International University (FIU). Animal care and procedures were approved and conducted in accordance with the Institutional Animal Care and Use Committee at FIU.

*Dnm1l*-deficient mouse models. The generation of *Dnm1l*^+/-^ mice with germline deletion of Exon 2 of the *Dnm1l* gene has been described in our recent publication^28^. For conditional DRP1-KO mouse models, we also contracted the European Conditional Mouse Mutagenesis Consortium (EUCOMM) to generate the *Dnm1l-Loxp*^+/+^ mice. These animals were generated using “knockout-first” technology^60^. The ‘knockout-first’ (tm1a) mice, which contain an IRES:*lacZ* trapping cassette and a floxed promoter-driven *neo* cassette inserted into the intron of the *Dnm1l* gene, were crossed with non-specific Flp mouse strain to converts the ‘knockout-first’ allele to the conditional allele (tm1c)^28^. *Cx3Cr1-Cre^ERT+/^*^-^ mice (#020940, Jackson Laboratory) and *Aldh1L1^Cre/ERT2+/^*^-^ mice (#031008, Jackson Laboratory) were subsequently crossed with *Dnm1l-Loxp*^+/+^ mice (tm1c) to generate *Cx3Cr1-Cre^ERT+/^*^-^:*Dnm1l-Loxp*^+/-^ mice (“Microglia-KO) and *Aldh1L1^Cre/ERT2+/^*^-^:*Dnm1l-Loxp*^+/-^ (“Astrocytes-KO”) mice, respectively. *Dnm1l-Loxp*^+/-^ mice were used as the control. These mice were injected with 75 mg/kg body weight of tamoxifen (dissolved in corn oil, cat. no. T2859, Sigma) for 5 consecutive days for the induction of Cre. The LPS treatment was performed for 3 weeks post-tamoxifen injection to knockout *Dnm1l*.

C57BL/6N-Tg(Thy1-SNCA)15Mjff/J (*Dnml1^+/+^*:*SNCA^+/-^*) mice were purchased from the Jackson Laboratory (#017682). *Dnml1^+/+^*:*SNCA^+/-^* mice were crossed with *Dnm1l*^+/-^:*SNCA^-/-^* (heterozygous DRP1-KO) mice to generate the *Dnm1l*^+/-^:*SNCN^+/-^* mice (double heterozygotes). *Dnml1^+/+^*:*SNCA^-/-^* (wild-type) mice were used as the control.

B6.129P2-*Lcn2tm1Aade*/AkiJ (*Lcn2^-/-^*) mice were purchased from the Jackson Laboratory (#024630). *Lcn2^-/-^* mice were crossed with C57BL/6 mice to generate *Lcn2^+/-^* mice. *Lcn2^+/-^* mice were paired with *Lcn2^+/-^* mice to maintain the colony.

LPS treatment. 3-month-old mice were injected intraperitoneally (i.p.) with a single dose of LPS (cat# L4391, Sigma-Aldrich) at the concentration of 5 mg/kg or saline (control). Mice were killed 6h later. Brains were micro-dissected by regions, snap-frozen or fixed for subsequent analysis.

### NanoString codeset

The reporter codeset used was the nCounter Mouse Neuroinflammation Panel (cat. no. 115000237, NanoString Technologies), which measures 757 target genes covering the core neuroinflammatory pathways and processes, together with 13 internal reference genes for data normalization.

### RNA extraction and NanoString nCounter analysis

Total RNA was extracted from mouse ventral midbrain using TRIzol (Thermo Fisher Scientific) following manufacturer’s instructions. Total RNA (75 ng per sample) was used for mRNA expression analysis on the NanoString nCounter SPRINT Profiler (NanoString Technologies). Target mRNA was hybridized with reporter-capture probes at 65 °C for 20 h according to the manufacturer’s protocol. The probe-target complexes were aligned and immobilized in the nCounter cartridge, which was then placed in the nCounter profiler to acquire images and process data. Gene expression levels were then tabulated in a comma-separated value format. The raw digital counts of target gene expression were subject to data analysis using nSolver v4.0 software (NanoString Technologies).

### Quantitative RT-PCR

For reverse transcription, 1 μg of total RNA was converted to complementary DNA (cDNA) using iScript Reverse Transcription Supermix (Bio-Rad). qPCR was assembled using TaqMan Fast Advanced Master Mix (Thermo Fisher Scientific) and Taqman assays prior to detection with QuantStudio 6 real-time PCR system (Thermo Fisher Scientific). The Taqman assays used were *Lcn2* (mouse, assay ID: Mm01324470_m1, Thermo Fisher Scientific), *Nlrp3* (mouse, assay ID: Mm00840904_m1, Thermo Fisher Scientific), *Il6* (mouse, assay ID: Mm00446190_m1, Thermo Fisher Scientific), *Il1b* (mouse, assay ID: Mm00434228_m1, Thermo Fisher Scientific), *Tnfα* (mouse, assay ID: Mm00443258_m1, Thermo Fisher Scientific), *Nos2* (mouse, assay ID: Mm00440502_m1, Thermo Fisher Scientific), *Rela* (mouse, assay ID: Mm00501346_m1, Thermo Fisher Scientific), and *Gapdh* (mouse, assay ID: Mm99999915_g1, Thermo Fisher Scientific). *Gapdh* served as internal control. The reaction conditions were as follows: 50 °C for 2 min, 95 °C for 2 min and 40 cycles of 95 °C for 1 s and 60 °C for 20 s. Relative quantification was performed using the 2^-ΔΔCT^ method.

### Meso Scale Discovery (MSD) multiplex immunoassay

*Dmn1l^+/-^* and *Dmn1l^+/+^* mice (3-month old) were treated with LPS (5 mg/kg, i.p.) or saline control for 6h prior to harvesting the ventral midbrains. Tissues were weighed and resuspended in 1X Tris lysis buffer (150 mM NaCl, 1 mM EDTA, 1 mM EGTA, 1% Triton X-100, and 20 mM Tris-HCl, pH 7.5) together with proteinase inhibitors (75mg tissue /0.5ml ice cold lysis buffer). Samples were briefly sonicated and further incubated on ice for 30 min followed by centrifugation at 20,000xg for 10 min at 4°C. Supernatant were collected for the assay and protein concentration was determined using BCA. For *in vitro*, primary microglia isolated from newborn pups were plated in 96-well plates and treated with 100 ng/ml LPS or vehicle control for 6 h. At the end of the treatment, conditioned media, which contained released cytokines, were collected from each well and transferred into a new plate. The levels of proinflammatory protein in the tissue lysates or secreted cytokines in the conditioned media were measured using a custom U-PLEX mouse panel for the following analytes, lipocalin 2, IL-6, IL-1β, TNF-α, IP-10, MCP-1/CCL2, and IFN-γ on the MSD platform following the manufacturer’s instructions. Plates were read on MSD QuickPlex SQ 120 and analyzed using the Discovery Workbench software.

### DAB staining and Sholl analysis

Mice were perfused and fixed as previously reported^61^. The brains were removed postfixed with 4% paraformaldehyde (PFA) overnight at 4 °C. Sequential overnight incubations in 15% and 30% sucrose in 0.1 M phosphate buffer were performed prior to snap-frozen of the brains in 2-methylbutane at -55 °C. Frozen brains were then mounted in optimal cutting temperature (OCT) medium, and cryo-sectioned at 30 µm using cryostat (CryoStar NX70, Thermo Fisher Scientific). Coronal sections were collected free-floating in 0.1 M tris-buffered saline (TBS). Sections from midbrain were washed in 0.1 M TBS before quenching endogenous peroxidases by incubation with 10% methanol and 0.3% H_2_O_2_ for 10 min at room temperature. Blocking was conducted in buffer containing 5% normal goat serum (NGS) (Vector Laboratories) and 0.1% TritonX-100 in 0.1M TBS for 1 h at room temperature, followed by overnight incubation at 4 °C with primary antibody solution containing antibody against IBA1 (1:1000, cat. no. 019-19741, FUJIFILM Wako Chemicals U.S.A. Corporation), 2% NGS, 0.1% Triton X-100 in 0.1M TBS. After washing with 0.1M TBS, sections were incubated with secondary antibody solution containing biotinylated goat anti-rabbit IgG antibody (1:200, cat. no. BA-1100, Vector Laboratories), 2% NGS in 0.1M TBS for 1 h at room temperature. Brain sections were washed in 0.1M TBS and subjected to incubation with ABC solution (Vectastain ABC kit, cat. no. PK-4000, Vector Laboratories) for 1 h. 3,3′-Diaminobenzidine (DAB peroxidase substrate kit, cat. no. SK-4100, Vector Laboratories) was applied to the slides for 10 min. Slides were then placed in water and ethanol of increasing concentrations (70%, 90%,100%). Coverslips were applied using DPX mounting medium (Sigma). Brightfield images were acquired using Nikon microscope with 100x objective. Individual cell branching was analyzed using the Sholl analysis plug-in for Fiji (http://fiji.sc/Sholl_Analysis) as previously reported^62^. Thirty microglia residing in the nigra were analyzed per group.

### Immunofluorescence-laser microdissection (IF-LMD)

Serial coronal nigral sections (10 µm) were cut on a standard cryostat (CryoStar NX70, Thermo Fisher Scientific), and eight sections from each brain (one hemisphere) were mounted on PEN membrane slides (4µm, cat. no. 11600288, Leica) prior to IF-LMD as previously reported^28^. Briefly, the brain sections were fixed in cold acetone for 5 min and subjected to wash with diethyl pyrocarbonate (DEPC)-treated 1x phosphate-buffered saline (PBS). The sections were then blocked with 5% normal goat serum (NGS) (Vector Laboratories) for 30 min, followed by incubation with primary antibodies for 1 h and secondary antibodies for 30 min at room temperature. Sections were rinsed with DEPC-treated 1x PBS between each step. After counterstaining with DAPI (10 µg/ml) for 5 min at room temperature, the sections were dehydrated in graded alcohols (30 seconds each) and air-dried. The primary antibodies used were chicken anti-tyrosine hydroxylase (1:200, cat. no. 76442, Abcam) and rabbit anti-IBA1 (1:100, cat. no. 019-19741, FUJIFILM Wako Chemicals U.S.A. Corporation) or mouse anti-GAD67 (1:50, cat. no. MAB5406, Millipore) and chicken anti-GFAP (1:200, cat. no. PA1-10004, Thermo Fisher Scientific). The secondary antibodies used were: Alexa Fluor 568 goat anti-chicken (cat. no. A11041, Thermo Fisher Scientific), Alexa Fluor 488 goat anti-rabbit (cat. no. A11008, Thermo Fisher Scientific), and Alexa Fluor 488 goat anti-mouse (cat. no. A11029, Thermo Fisher Scientific). All secondary antibodies were used at 1:100 dilution. The immunostained DA neurons in the substantia nigra *pars compacta*, and GABA neurons in the substantia nigra *pars reticulata* as well as microglia and astrocytes in the substantia nigra were subsequently collected by laser microdissection (LMD6, Leica). A total of 40 cells per cell type were collected for subsequent analysis.

### Reverse transcription and cDNA preamplification using Smart-seq2 method

Laser microdissected cells (40 cells per cell type) were collected into a 0.2 ml thin-wall PCR tube containing 2 μl of a mild hypotonic lysis buffer composed of 0.2% Triton X-100 and 2 U/μl of RNase inhibitor (Clontech). Total RNA in cell lysates was subjected to reverse transcription and cDNA preamplification following Smart-seq2 method^63, 64^. Briefly, cell lysates were mixed with 1 μl of oligo-dT primer (10 μM, Integrated DNA Technologies) and 1 μl of dNTP mix (10 mM, Thermo Fisher Scientific), denatured at 72 °C for 3 min and immediately placed on ice afterwards. Reverse transcription reaction was carried out with Superscript II reverse transcriptase (Thermo Fisher Scientific) by incubation at 42 °C for 90 min, 10 cycles of 50 °C for 2 min and 42 °C for 2 min, followed by enzyme inactivation at 70 °C for 15 min. Subsequently, cDNA preamplification was performed using KAPA HiFi HotStart Ready Mix (Roche). The program used was 98 °C for 3 min, 24 cycles of 98 °C for 15 s, 67 °C for 20 s and 72 °C for 6 min, with a final extension at 72 °C for 5 min. PCR products were purified using AMPure XP beads (Beckman Coulter) following manufacturer’s instructions and eluted in 15 μl of TE buffer. The pre-amplified cDNA was then subjected to qPCR as described earlier.

### Primary mouse microglia

Primary mouse microglia were prepared from post-natal 0-1 day old mouse pups as previously described^65^. Briefly, following decapitation, whole brains and their meninges were removed, then placed in 0.25% trypsin and incubated at 37 °C for 15 min with gentle inversion every 5 min. The mixture was then triturated and passed through a 70 μm strainer prior to culturing in flasks for 16 days in Dulbecco’s modified Eagle’s medium/F12 (DMEM/F12) supplemented with 10% heat-inactivated fetal bovine serum (HI-FBS), 2 mM L-glutamine, 1 mM sodium pyruvate, 1% non-essential amino acids, and 1% penicillin/streptomycin. Full medium changes were performed on day 6 after the initial plating. On day 16, microglia were separated from other cell types by a CD11b immunomagnetic positive selection kit (cat. no. 18970, STEMCELL Technologies) and plated for experiments. HI FBS was reduced to 2% for all treatments. Cell purity (>95%) was confirmed by immunofluorescence using Iba1 as a marker for microglia.

### Adult microglia isolation and mitochondrial respiration assay

WT mice (2-4 month-old) were injected with LPS (5mg/kg, i.p.) or saline control. Six hours post-injection, microglia enrichment was performed using gradient centrifugation as previously described with some modifications.^66, 67^ Mice were perfused with ice-cold sterile 0.9% saline, and brains were quickly removed following decapitation and collected in ice cold Hirbernate A medium containing B27, Glutamax, penicillin-streptomycin and DNase I (HABG)^67^. Brains were then placed on top of sterile filter paper pre-wet with HABG on the cooled stage, and finely mince with a sterile scalpel. The minced brains were subsequently transferred into dissociation DMEM/F12 medium containing 1mg/ml papain, 1.2U/ml dispase II and 20 U/mL Dnase I, and incubated at 30°C for 20 min with gentle inversion every 5 min. Neutralization solution was added prior to the transfer of brains into new tubes, followed by addition of 2ml HABG. The brains were then gently triturated with a 9-in Pasteur pipette with fire-polished tip, and the resulting supernatants were transferred into a new tube with neutralization solution. The supernatants were filtered using a 70 µm strainer and subjected to centrifugation at 400xg for 5min at 4°C. The pellets were then gently resuspended in 6ml HABG and carefully applied on top of a freshly prepared 4-layer Opti-prep density ingredient (top to bottom: 7.4%, 9.9%, 12.4%, and 17.4% diluted in Hibernate A/B27 medium)^66^. The gradients were centrifuged at 800×g for 30 min at 22 °C (with mild acceleration and deceleration rate set at 0). The top layers were aspirated carefully and the bottom fractions enriched with microglia^66^ were collected and diluted into 5 ml cold HABG medium, followed by centrifugation at 400×g for 10 min to remove Opti-Prep. The pellets were resuspended, counted, and plated onto assay plate. The plate was further centrifuged at 1000xg for 15min to attach the cells before topping up with assay buffer and climatized in the non-CO2 incubator for 30min. A standard mito-stress assay was performed as described below using XF^e^ 96 Extracellular Flux Analyzer with extended measurement for basal level.

### Mitochondrial respiration assay

Mitochondrial respiration was assessed using Agilent XF^e^ 96 Extracellular Flux Analyzer (Agilent Inc) as previously reported^65^. Briefly, primary microglia from *Dnm1l*^+/+^ and *Dnm1l*^+/-^ mouse pups were seeded into XF Cell Culture Microplate and treated with LPS (100 ng/ml) the next day for 2h, 6h and 24h, respectively. On the day of the assay, cells were washed two times with XF assay media containing DMEM, 1 mM pyruvate, 2 mM Glutamax, 2 mM glucose and 2 mM HEPES, pH 7.2. The plate was then acclimatized in a non-CO_2_ incubator for 30 min and placed in the Analyzer. A standard XF cell mito-stress test with sequential injections of oligomycin, carbonyl cyanide-p-trifluoromethoxyphenylhydrazone (FCCP) and rotenone/antimycin was conducted.

### Immunostaining

Primary mouse microglia were grown on borosilicate cover slips pre-coated with poly-D-lysine in 24-well plates. After treatment with LPS (100 ng/ml) for 2, 6 or 24 h, cells were fixed and subjected to immunostaining as described previously^65^. Primary antibodies used were rabbit anti-TOM20 (1:1000, cat. no. ab186735, Abcam), mouse anti-DRP1 (1:500, cat. no. 611113, BD Biosciences), rabbit anti-DRP1 (1:500, cat. no. NB110-55288, Novus Biologicals), and goat anti-IBA1 antibody (1:500, cat. no. NB100-1028, Novus Biologicals). Corresponding Alexa Fluor 488, 594, and 633 conjugate secondary antibodies (Invitrogen) were used at 1:1000 dilution. Samples were incubated on an orbital shaker with primary antibodies overnight at 4°C, and in secondary antibodies at room temperature for 1h. Cover slips were then mounted onto slides using Prolong™ gold anti-fade mount with DAPI (Thermo Fisher Scientific).

Mice were immediately perfused with 0.9% saline followed by 4% PFA (prepared in 0.1M phosphate buffer) after exposure to LPS for 6h. Saline-treated age-matched littermates were employed as the vehicle control. Brains were removed and subjected to post-fixation and snap-frozen with 2-methylbutane. Frozen brains were mounted in OCT medium, and cryo-sectioned at 30µm using a cryostat. Coronal sections were collected free-floating in 0.1M TBS prior to immunostaining. Primary antibodies used were rabbit anti-DRP1 (1:500, cat. no. NB110-55288, Novus Biologicals), and goat anti-IBA1 antibody (1:500, cat. no. NB100-1028, Novus Biologicals). Alexa Fluor secondary antibodies were used at 1:1000 dilution (Invitrogen). Following immunostaining with antibodies, sections were stained with DAPI (5 ug/ml, cat. no. 62247, Thermo Fisher Scientific) for 10 min and washed three times with 0.1M TBS before mounted onto microscope slides, cover glassed with Prolong gold anti-fade mount (Thermo Fisher Scientific). All fluorescent images were captured using Olympus Fluoview (FV10i) confocal microscope with 60x objective.

### Quantification of mitochondrial morphology and network

Mitochondrial morphology/network was quantified using the Mitochondrial Network Analysis (MiNA) plugin of Fiji as described previously^28^. Mitochondria were visualized by immunofluorescence using TOM20 (Abcam, ab186735) and imaged with the Olympus Fluoview microscope. The Z-stack images were subjected to MiNA analysis. “Mean rod/branch length” indicates the average length of all rods and branches, whereas “Mean branches per Network” represents the average number of branches per network. In addition, for individual mitochondrial morphology: roundness: 4 x ([Area])/(π × [Major axis]^2^ and aspect ratio (major / minor axes) were quantified using Fiji as previously described^65^. Both of these values approach 1 as the particle becomes circular.

### Quantification using Imaris

The analyses of DRP1 nuclear translocation in primary mouse microglia or microglia in the nigra of adult mouse brains were performed using IMARIS (Oxford instruments). Z-stack confocal microscopy images using 60x objective lens, 3X optical zoom. In brief, the 3D construct of nuclei and cell surfaces were rendered from blue and cyan channels using the “Surfaces” module, respectively. The puncta for DRP1 were generated from the red fluorescence channel using the “Spots” module using appropriate threshold that was maintained between treatment groups. The number of DRP1^Nuc^ was determined by quantification of red puncta inside the nuclei volume, whereas DRP1^Cyto^ was defined by total number of puncta inside the cellular volume, subtracting the number of DRP1^Nuc^.

### Nuclear and cytoplasmic fractionation

To quantitatively measure the enrichment of DRP1 in the nucleus upon LPS exposure, the nuclear and cytoplasmic extracts were prepared from LPS-treated primary mouse microglia, respectively. Vehicle-treated cells were used as the control. Briefly, cells were harvested into 100 μl of “Cytoplasmic Extract Buffer” containing 10mM HEPES, pH 7.6, 60mM KCl, 1mM EDTA, and 0.075% NP-40, 1mM DTT, 1mM PMSF and incubated on ice for 10 min to release cytoplasm. Cell lysates were then subjected to centrifugation at 200x g for 4 min at 4°, allowing for the separation of cytoplasmic extracts from the precipitated nucleus. The supernatant was collected as the crude cytoplasmic lysates, whereas the nucleus pellets were washed with the “Cytoplasmic Extract Buffer” in the absence of NP-40. Following centrifugation at 200x g for 4 min at 4°, the nucleus pellets were resuspended in “Nuclear Extract Buffer” composed of 20mM Tris-HCl, pH 8.0, 420 mM NaCl, 1.5 mM MgCl_2_, 0.2 mM EDTA, 0.5 mM DTT, 25% glycerol and 1mM PMSF. The nucleus was homogenized using a Qsonica sonicator at 4°C for 3 cycles of 30 s ON and 30 s OFF at 60% amplitude. Both crude cytoplasmic and nuclear lysates were then subjected to centrifugation at 21,000x g for 30 min at 4°C. The supernatant, i.e., cytoplasmic extract and nuclear extract, was aliquoted and stored at -80°C for subsequent analysis.

### Capillary gel-free immunoblotting

Capillary immunoblotting was conducted with Jess™ system (ProteinSimple) using 12-230 kDa Fluorescence Separation Module (cat. no. SM-FL004, ProteinSimple) as described previously^28^. After protein quantification using Micro BCA kit (cat. no. 23235, Thermo Fisher Scientific), cytoplasmic extract and nuclear extract were diluted in 0.1X sample buffer and mixed with 5x Fluorescent Master Mix (cat. no. PS-ST01EZ, ProteinSimple) to a final concentration of 0.5µg/µl and 1.2µg/µl, respectively. The samples were then boiled at 95 °C for 5 min, and 3 µl of protein were loaded for each sample. The primary antibodies used were rabbit anti-DRP1 (1:100, cat. no. NB110-55288, Novus Biologicals), rabbit anti-Histone H3 (1:100, cat. no. ab1791, Abcam) and rabbit anti-β tubulin (1:1000, cat. no. NBP1-48320, Novus Biologicals). Anti-rabbit secondary antibodies were prepared using corresponding detection modules (cat. no. DM001, ProteinSimple) and Protein Normalization Module (cat. no. DM-PN02, ProteinSimple) was used for normalization. Data were recorded and analyzed using Compass v6.0.

### Chromatin immunoprecipitation (ChIP) assay

A total of 6 x 10^6^ primary microglia were treated with 100 ng/ml LPS for 6 h. Primary microglia treated with medium only were used as control. At the end of the treatment, cells were washed twice using PBS containing proteinase inhibitors and subjected to ChIP assay. Briefly, primary microglia were supplied with medium containing 1% (w/v) formaldehyde (Sigma-Aldrich) and incubated at 37°C for 15 min (for NF-ĸB p65) or 30 min (for DRP1) for crosslinking reaction that was subsequently quenched by 125 mM glycine for 5 min with shaking. Cells were collected through scraping and centrifugation. Cell pellets were washed twice with cold PBS containing protease inhibitors (Pierce-Thermo Fisher Scientific) and suspended in cell lysis buffer with 1% (w/v) SDS, 10 mM EDTA, and 50 mM Tris HCl, pH=8.0, and protease inhibitors. Cell suspension was then incubated on ice for 10 min and subjected to lysis by sonication with Qsonica sonicator at 4°C for 30 cycles of 10 s ON and 30 s OFF at 60% amplitude for cell lysis and DNA fragmentation. Cell lysates were centrifuged at 18,000 x g for 10 min at 4°C to separate the supernatant from cell debris. Subsequently, the supernatant was diluted 10-fold with ice-cold ChIP dilution buffer containing contained 1% (v/v) Triton X-100, 1.2 mM EDTA, 167 mM NaCl and 16.7 mM Tris–HCl, pH 8.0, with protease inhibitors. Supernatant (100 µl) was taken as “Input”.

The remaining supernatant was incubated with sheared salmon sperm DNA-coated protein A agarose (Pierce-Thermo Fisher Scientific) for 2h at 4°C with rotation. The supernatant was then divided into equal aliquots that were incubated overnight at 4°C with 2 µg IgG (Cell Signaling Technology, #2729) (IgG control) or 2 µg anti-NF-ĸB p65 antibody (Cell Signaling Technology, #8242), and anti-DRP1 antibody (Cell Signaling Technology, #8570). Immunoprecipitations (IPs) of NF-ĸB p65, DRP1 and IgG were incubated with sheared salmon sperm DNA-coated protein A agarose beads for 2 h at 4°C with rotation. IPs bound to the beads were centrifuged at 500x g for 2 min and washed twice with low salt washing buffer containing 150 mM NaCl, followed by washing with high salt buffer with 500 mM NaCl, and finally washed with TE buffer. The IPs were eluted from the beads with freshly made elution buffer containing 1% SDS and 0.1 M NaHCO_3_. The IPs were incubated with 0.2M NaCl at 65°C for overnight for DNA-protein crosslink reversal, leading to the release of the crosslinked DNA from the protein. The IPs were then subjected to proteinase K digestion at 45°C for 2h to remove protein. The released DNA was treated with phenol/chloroform extraction, precipitated with ethanol, and redissolved in TE buffer for subsequent quantitative PCR amplification.

### Quantitative real-time PCR for ChIP assay

The quantitative real-time PCR reaction was assembled with KAPA SYBR FAST qPCR Master Mix (Sigma), purified DNA bound by NF-ĸB p65 or DRP1 and IgG, the forward and reverse PCR primers (Supplementary Table 2) in 10 µl reaction mixture. The PCR amplification was performed using QuantStudio 6 detection system (Thermo Fisher Scientific) with the PCR program: 95°C for 3 min for 1 cycle, 95°C for 3 s, Ta (Table S2) for 20 s, 72°C for 1 s for 40 cycles. Ct values recorded by the QuantStudio Software v1.3 (Thermo Fisher Scientific) during PCR amplification was used to calculate the fold-difference between the amount of DNA bound by NF-ĸB p65 or DRP1 and IgG as well as Input. ΔCt [normalized ChIP] (normalized to the Input) = Ct [ChIP]-(Ct [Input] -log2 [Input dilution factor]), where the Input dilution factor= the fraction volume used for each IP/the fraction volume of Input saved for further analysis. In this study, the fraction volume of Input was 100 μl, and the fraction volume for each IP was 900 μl. Thus, the Input dilution factor was 9. For our experiments, the equation was ΔCt [normalized ChIP] = Ct [ChIP]-(Ct [Input] -log2 [9]). The Input %= 2^-ΔCt [normalized ChIP]^ × 100. The “Input %” indicates the enrichment of NF-ĸB p65 or DRP1 and IgG on the promoter region of their target gene(s).

### Statistics

All values are expressed as mean ± SEM. Differences between means were analyzed using t-test, one-, two-, or three-way ANOVA with different mouse genotypes, treatment or time as independent factors. When ANOVA shows significant differences, Tukey’s post-hoc test was used for pair-wise comparisons between means. All data sets were subjected to a normality test and an equality of variance test. When these criteria were not met, nonparametric analysis was applied (Kruskal-Wallis ANOVA test with Dunn’s post-hoc test). In all analyses, the null hypothesis (H_0_) or the alternative hypothesis (H_1_) was accepted with an α-error (false-positive) ≤ 5% and a β-error (false-negative) ≤ 20%, respectively. F values, degree of freedom and their p values for all data analysis are provided in Supplementary Table S3.

## REFERENCES

1. Kwon, H.S. & Koh, S.H. Neuroinflammation in neurodegenerative disorders: the roles of microglia and astrocytes. Transl Neurodegener 9, 42 (2020).

2. Ransohoff, R.M. How neuroinflammation contributes to neurodegeneration. Science 353, 777–783 (2016).

3. Mitchell, J.P. & Carmody, R.J. NF-kappaB and the Transcriptional Control of Inflammation. Int Rev Cell Mol Biol 335, 41–84 (2018).

4. Dresselhaus, E.C. & Meffert, M.K. Cellular Specificity of NF-kappaB Function in the Nervous System. Front Immunol 10, 1043 (2019).

5. Hayden, M.S. & Ghosh, S. NF-kappaB, the first quarter-century: remarkable progress and outstanding questions. Genes Dev 26, 203–234 (2012).

6. Vallabhapurapu, S. & Karin, M. Regulation and function of NF-kappaB transcription factors in the immune system. Annu Rev Immunol 27, 693–733 (2009).

7. Hayden, M.S. & Ghosh, S. Shared principles in NF-kappaB signaling. Cell 132, 344–362 (2008).

8. Lawrence, T. The nuclear factor NF-kappaB pathway in inflammation. Cold Spring Harb Perspect Biol 1, a001651 (2009).

9. Oeckinghaus, A. & Ghosh, S. The NF-kappaB family of transcription factors and its regulation. Cold Spring Harb Perspect Biol 1, a000034 (2009).

10. Pahl, H.L. Activators and target genes of Rel/NF-kappaB transcription factors. Oncogene 18, 6853–6866 (1999).

11. Jaberi, S.A., et al. Lipocalin-2: Structure, function, distribution and role in metabolic disorders. Biomed Pharmacother 142, 112002 (2021).

12. Xiao, X., Yeoh, B.S. & Vijay-Kumar, M. Lipocalin 2: An Emerging Player in Iron Homeostasis and Inflammation. Annu Rev Nutr 37, 103–130 (2017).

13. Jha, M.K., et al. Diverse functional roles of lipocalin-2 in the central nervous system. Neurosci Biobehav Rev 49, 135–156 (2015).

14. Kim, S.L., Shin, M.W. & Kim, S.W. Lipocalin 2 activates the NLRP3 inflammasome via LPS-induced NF-kappaB signaling and plays a role as a pro-inflammatory regulator in murine macrophages. Mol Med Rep 26 (2022).

15. Ip, J.P., et al. Lipocalin 2 in the central nervous system host response to systemic lipopolysaccharide administration. J Neuroinflammation 8, 124 (2011).

16. Chakraborty, S., Kaur, S., Guha, S. & Batra, S.K. The multifaceted roles of neutrophil gelatinase associated lipocalin (NGAL) in inflammation and cancer. Biochim Biophys Acta 1826, 129–169 (2012).

17. Xiong, M., Qian, Q., Liang, X. & Wei, Y.D. Serum levels of lipocalin-2 in patients with Parkinson’s disease. Neurol Sci 43, 1755–1759 (2022).

18. Kim, B.W., et al. Pathogenic Upregulation of Glial Lipocalin-2 in the Parkinsonian Dopaminergic System. The Journal of neuroscience : the official journal of the Society for Neuroscience 36, 5608–5622 (2016).

19. Eruysal, E., Ravdin, L., Kamel, H., Iadecola, C. & Ishii, M. Plasma lipocalin-2 levels in the preclinical stage of Alzheimer’s disease. Alzheimers Dement (Amst) 11, 646–653 (2019).

20. Naude, P.J., et al. Lipocalin 2: novel component of proinflammatory signaling in Alzheimer’s disease. FASEB J 26, 2811–2823 (2012).

21. Dekens, D.W., et al. Lipocalin 2 contributes to brain iron dysregulation but does not affect cognition, plaque load, and glial activation in the J20 Alzheimer mouse model. J Neuroinflammation 15, 330 (2018).

22. Lee, S., et al. A dual role of lipocalin 2 in the apoptosis and deramification of activated microglia. J Immunol 179, 3231–3241 (2007).

23. Bi, F., et al. Reactive astrocytes secrete lcn2 to promote neuron death. Proc Natl Acad Sci U S A 110, 4069–4074 (2013).

24. Kim, J.H., et al. Aberrant activation of hippocampal astrocytes causes neuroinflammation and cognitive decline in mice. PLoS Biol 22, e3002687 (2024).

25. Knott, A.B., Perkins, G., Schwarzenbacher, R. & Bossy-Wetzel, E. Mitochondrial fragmentation in neurodegeneration. Nat Rev Neurosci 9, 505–518 (2008).

26. Oliver, D. & Reddy, P.H. Dynamics of Dynamin-Related Protein 1 in Alzheimer’s Disease and Other Neurodegenerative Diseases. Cells 8 (2019).

27. Brown, H.J., et al. Imbalanced mitochondrial dynamics in human PD and alpha-synuclein mouse brains. Neurobiol Dis 212, 106976 (2025).

28. Fan, R.Z., et al. A partial Drp1 knockout improves autophagy flux independent of mitochondrial function. Mol Neurodegener 19, 26 (2024).

29. Smirnova, E., Griparic, L., Shurland, D.L. & van der Bliek, A.M. Dynamin-related protein Drp1 is required for mitochondrial division in mammalian cells. Mol Biol Cell 12, 2245–2256 (2001).

30. Ji, W.K., Hatch, A.L., Merrill, R.A., Strack, S. & Higgs, H.N. Actin filaments target the oligomeric maturation of the dynamin GTPase Drp1 to mitochondrial fission sites. Elife 4, e11553 (2015).

31. Itoh, K., et al. A brain-enriched Drp1 isoform associates with lysosomes, late endosomes, and the plasma membrane. J Biol Chem 293, 11809–11822 (2018).

32. Yoon, Y., Pitts, K.R., Dahan, S. & McNiven, M.A. A novel dynamin-like protein associates with cytoplasmic vesicles and tubules of the endoplasmic reticulum in mammalian cells. J Cell Biol 140, 779–793 (1998).

33. Koch, A., et al. Dynamin-like protein 1 is involved in peroxisomal fission. J Biol Chem 278, 8597–8605 (2003).

34. Joshi, A.U., et al. Fragmented mitochondria released from microglia trigger A1 astrocytic response and propagate inflammatory neurodegeneration. Nat Neurosci 22, 1635–1648 (2019).

35. Park, J., et al. Mitochondrial dynamics modulate the expression of pro-inflammatory mediators in microglial cells. J Neurochem 127, 221–232 (2013).

36. Qin, L., et al. Systemic LPS causes chronic neuroinflammation and progressive neurodegeneration. Glia 55, 453–462 (2007).

37. Zhao, Z., et al. A novel role of NLRP3-generated IL-1β in the acute-chronic transition of peripheral lipopolysaccharide-elicited neuroinflammation: implications for sepsis-associated neurodegeneration. J Neuroinflammation 17, 64 (2020).

38. Zhao, P., Elks, C.M. & Stephens, J.M. The induction of lipocalin-2 protein expression in vivo and in vitro. J Biol Chem 289, 5960–5969 (2014).

39. Kim, S.L., Shin, M.W., Seo, S.Y. & Kim, S.W. Lipocalin 2 potentially contributes to tumorigenesis from colitis via IL-6/STAT3/NF-kappaB signaling pathway. Biosci Rep 42 (2022).

40. Lee, J., Rhee, M.H., Kim, E. & Cho, J.Y. BAY 11-7082 is a broad-spectrum inhibitor with anti-inflammatory activity against multiple targets. Mediators Inflamm 2012, 416036 (2012).

41. Chiang, Y.Y., et al. Nuclear expression of dynamin-related protein 1 in lung adenocarcinomas. Mod Pathol 22, 1139–1150 (2009).

42. Shih, R.H., Wang, C.Y. & Yang, C.M. NF-kappaB Signaling Pathways in Neurological Inflammation: A Mini Review. Front Mol Neurosci 8, 77 (2015).

43. Wen, H., Miao, E.A. & Ting, J.P. Mechanisms of NOD-like receptor-associated inflammasome activation. Immunity 39, 432–441 (2013).

44. Sutterwala, F.S., Haasken, S. & Cassel, S.L. Mechanism of NLRP3 inflammasome activation. Ann N Y Acad Sci 1319, 82–95 (2014).

45. Dekens, D.W., et al. Lipocalin 2 as a link between ageing, risk factor conditions and age-related brain diseases. Ageing Res Rev 70, 101414 (2021).

46. Song, E., et al. Lipocalin-2 induces NLRP3 inflammasome activation via HMGB1 induced TLR4 signaling in heart tissue of mice under pressure overload challenge. Am J Transl Res 9, 2723–2735 (2017).

47. Qin, L., et al. Systemic LPS causes chronic neuroinflammation and progressive neurodegeneration. Glia 55, 453–462 (2007).

48. Skrzypczak-Wiercioch, A. & Sałat, K. Lipopolysaccharide-Induced Model of Neuroinflammation: Mechanisms of Action, Research Application and Future Directions for Its Use. Molecules 27 (2022).

49. Gupta, U., et al. Increased LCN2 (lipocalin 2) in the RPE decreases autophagy and activates inflammasome-ferroptosis processes in a mouse model of dry AMD. Autophagy 19, 92–111 (2023).

50. Lee, J.W., et al. TLR4 (toll-like receptor 4) activation suppresses autophagy through inhibition of FOXO3 and impairs phagocytic capacity of microglia. Autophagy 15, 753–770 (2019).

51. Francois, A., et al. Impairment of autophagy in the central nervous system during lipopolysaccharide-induced inflammatory stress in mice. Mol Brain 7, 56 (2014).

52. Russell, A.E., et al. Intermittent Lipopolysaccharide Exposure Significantly Increases Cortical Infarct Size and Impairs Autophagy. ASN Neuro 13, 1759091421991769 (2021).

53. Purnell, P.R. & Fox, H.S. Autophagy-mediated turnover of dynamin-related protein 1. BMC Neurosci 14, 86 (2013).

54. Bordt, E.A., et al. The Putative Drp1 Inhibitor mdivi-1 Is a Reversible Mitochondrial Complex I Inhibitor that Modulates Reactive Oxygen Species. Dev. Cell 40, 583–594 (2017).

55. Macia, E., et al. Dynasore, a cell-permeable inhibitor of dynamin. Developmental Cell 10, 839–850 (2006).

56. Nishiyama, T., et al. Concise synthesis and antiproliferative activity evaluation of ellipticine quinone and its analogs. Eur J Med Chem 136, 1–13 (2017).

57. Qi, X., Qvit, N., Su, Y.C. & Mochly-Rosen, D. A novel Drp1 inhibitor diminishes aberrant mitochondrial fission and neurotoxicity. J. Cell Sci 126, 789–802 (2013).

58. Guo, X., et al. Inhibition of mitochondrial fragmentation diminishes Huntington’s disease-associated neurodegeneration. J. Clin. Invest 123, 5371–5388 (2013).

59. Filichia, E., Hoffer, B., Qi, X. & Luo, Y. Inhibition of Drp1 mitochondrial translocation provides neural protection in dopaminergic system in a Parkinson’s disease model induced by MPTP. in Sci Rep 32656 (2016).

60. Skarnes, W.C., et al. A conditional knockout resource for the genome-wide study of mouse gene function. Nature 474, 337–342 (2011).

61. Rappold, P.M., et al. Drp1 inhibition attenuates neurotoxicity and dopamine release deficits in vivo. Nat Commun 5, 5244 (2014).

62. Ferreira, T.A., et al. Neuronal morphometry directly from bitmap images. Nat Methods 11, 982–984 (2014).

63. Picelli, S., et al. Smart-seq2 for sensitive full-length transcriptome profiling in single cells. Nat Methods 10, 1096–1098 (2013).

64. Picelli, S., et al. Full-length RNA-seq from single cells using Smart-seq2. Nat Protoc 9, 171–181 (2014).

65. Fan, R.Z., Guo, M., Luo, S., Cui, M. & Tieu, K. Exosome release and neuropathology induced by alpha-synuclein: new insights into protective mechanisms of Drp1 inhibition. Acta neuropathologica communications 7, 184 (2019).

66. Brewer, G.J. & Torricelli, J.R. Isolation and culture of adult neurons and neurospheres. Nat Protoc 2, 1490–1498 (2007).

67. Zheng, J., et al. Single-cell RNA-seq analysis reveals compartment-specific heterogeneity and plasticity of microglia. iScience 24, 102186 (2021).

