## Supplemental Information for "DRP1 induces neuroinflammation *via* transcriptional regulation of NF-ĸB"

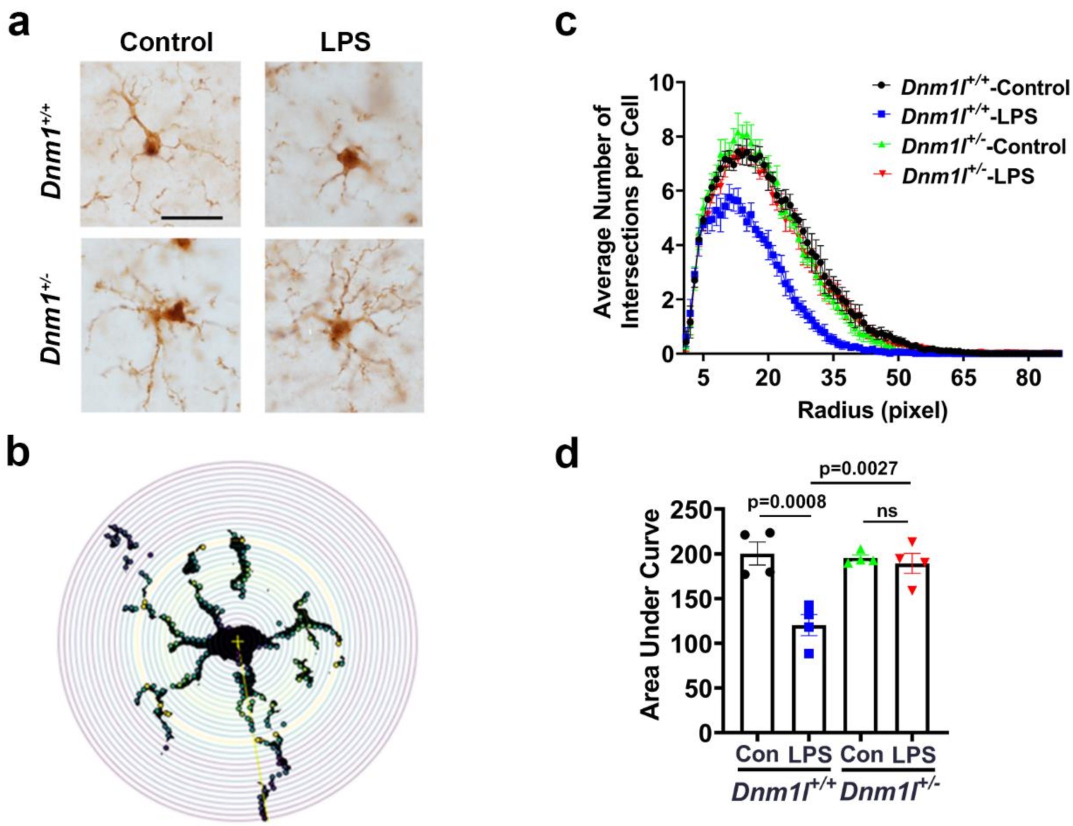


**Figure S1. Reduced microglia activation in LPS-treated *Dnm1l*$^{+/-}$ mice.**

LPS (5 mg/kg, i.p.) or saline control was delivered to 3-month-old *Dnm1l*$^{+/-}$ and *Dnm1l*$^{+/+}$ mice and the VMB was collected 6h later for DAB staining (**a**) and Sholl analysis (**b**-**d**). **a**, Coronal midbrain sections were immunostained for IBA-1 and visualized with DAB. Images of 30 microglia from the nigra of each animal were captured at 100X, n =4 (2M & 2F). **b**, Morphological changes were assessed using Sholl analysis performed by investigators blinded to experimental treatments. **c**, Average number of intersections per cell was obtained from Sholl analysis. **d**, Area under the curve was generated from Sholl analysis and quantified. Data were analyzed using two-way ANOVA with Tukey's post hoc analysis.

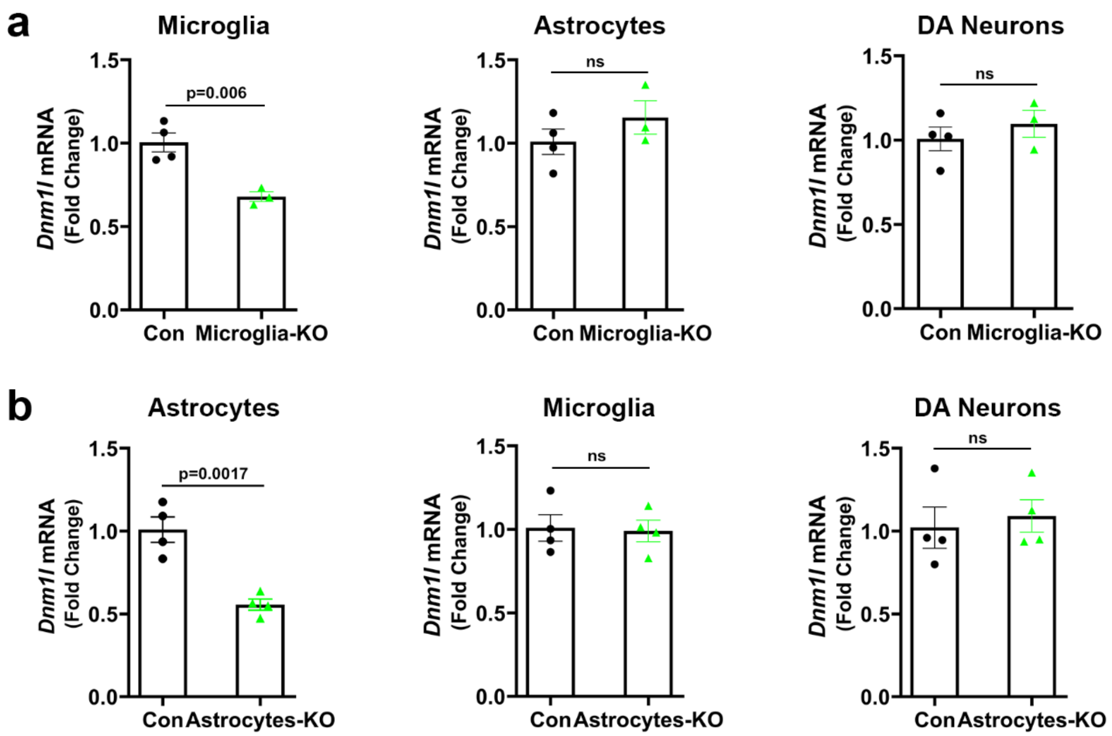


**Figure S2. *Dnm1l* expression in different cell types in conditional *Dnm1l*-KO mouse models.**

Brain tissue was collected from 3-month old "Microglia-KO" (**a**) and "Astrocytes-KO" (**b**) mice 3 weeks post-tamoxifen injection. Tamoxifen injected *Dnm1l-Loxp*$^{+/-}$ mice were used as the control. Snap-frozen brains were sectioned at 10 µm, collected on PEN membrane slides, immunostained with antibodies against IBA1, GFAP and TH. Microglia, astrocytes and DA neurons were captured using laser microdissection. "Smart-seq2" method was used to pre-amplify cDNA of 40 cells from each cell type in the nigra of each animal for qPCR analysis of *Dnm1l* (normalized to *Gapdh*). N=3-4 (1-2 F & 2 M) mice per group. Data represent mean ± SEM, unpaired t-tests.

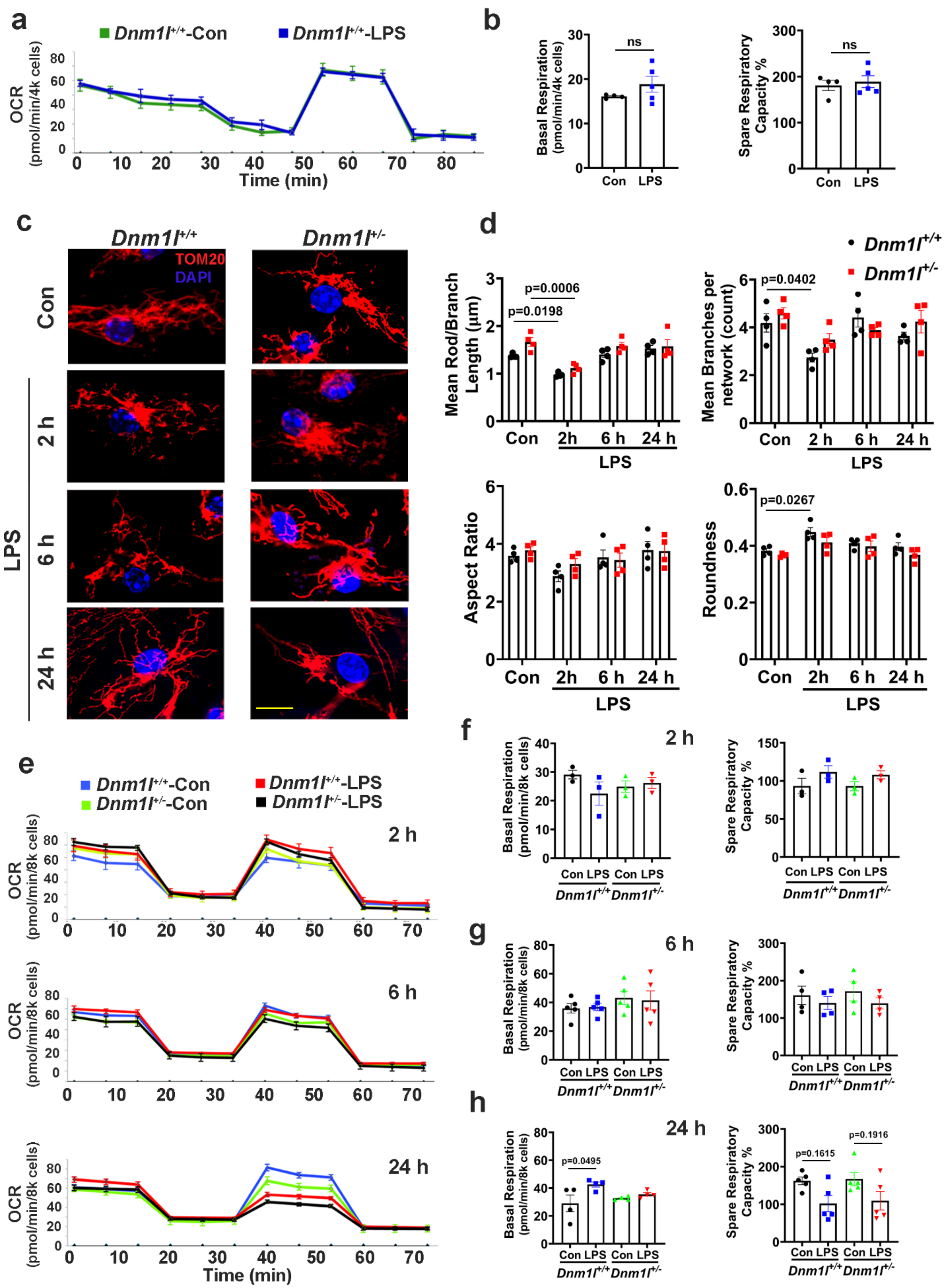
a
Dnm1l+/+-Con
Dnm1l+/+-LPS
OCR (pmol/min/4k cells)
Time (min)
b
Basal Respiration (pmol/min/4k cells)
Spare Respiratory Capacity %
ns
Con LPS
c
Dnm1l+/+
Dnm1l+/-
TOM20
DAPI
Con
LPS
2 h
6 h
24 h
d
Dnm1l+/+
Dnm1l+/-
Mean Rod/Branch Length (µm)
p=0.0006
p=0.0198
Mean Branches per network (count)
p=0.0402
Aspect Ratio
Roundness
p=0.0267
Con 2h 6 h 24 h
LPS
e
Dnm1l+/+-Con
Dnm1l+/--Con
Dnm1l+/+-LPS
Dnm1l+/--LPS
OCR (pmol/min/8k cells)
2 h
6 h
24 h
Time (min)
f
g
h
Basal Respiration (pmol/min/8k cells)
Spare Respiratory Capacity %
Con LPS Con LPS
Dnm1l+/+ Dnm1l+/-
p=0.0495
p=0.1615
p=0.1916

**Figure S3. Mitochondrial morphology and function in LPS-treated primary microglia.** Microglia from *Dnm1l*$^{+/+}$ and *Dnm1l*$^{+/-}$ pups were treated with LPS (100 ng/ml) or vehicle control for 2h, 6h and 24h, respectively. All data were analyzed using two-way ANOVA, followed by Tukey's post hoc test. **a**,**b**, 2-4 month old *Dnm1l*$^{+/+}$ mice were injected with LPS (5 mg/kg, i.p.) or saline control for 6h. Brain microglia were isolated and enriched for the measurement of mitochondrial respiration. Representative kinetic graphs of mito-stress assay (**a**) and the corresponding quantified data (**b**). Data represent mean±SEM, N=4-5 (2 F & 2-3 M) mice per group and analyzed using t-tests. **c**, Representative confocal images of TOM20 immunostained for mitochondrial morphology. **d,** Mitochondrial morphology/network was quantified using the Mitochondrial Network Analysis plugin of Fiji. N=4 independent experiments with 20 cells/experiment. **e**, Mitochondrial respiration was assessed using the XFe96 Extracellular Flux Analyzer. Representative kinetic graphs of mito-stress assay for 2h, 6h and 24h time-points, respectively. **f** Quantified data for oxygen consumption rate (OCR) 2h after LPS treatment. N=3 independent experiments with 6-8 replicates / experiment. **g**, Quantified data for OCR 6h after LPS treatment. N=4-5 independent experiments with 6-8 replicates / experiment. **h**, Quantified data for OCR 24h after LPS treatment. N=4-5 independent experiments with 6-8 replicates / experiment.

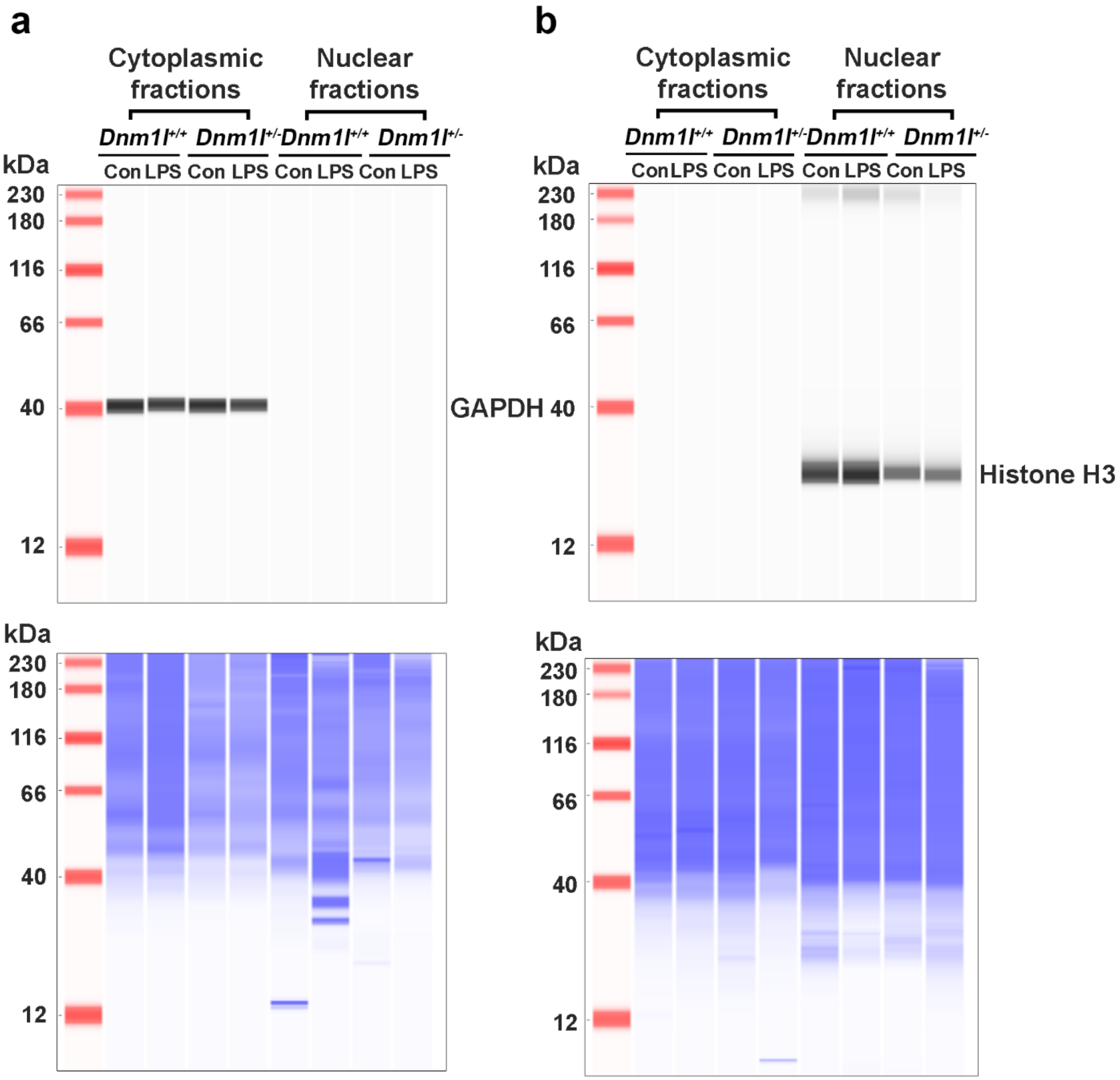


**Figure S4. Purity of cytoplasmic and nuclear fractions.**

*Dnm1l*$^{+/+}$ and *Dnm1l*$^{+/-}$ primary microglia were treated with 100 ng/ml LPS for 6h followed by nuclear and cytoplasmic fractionation. **a**, Immunoblotting of GAPDH in the cytoplasmic and nuclear fractions (top panel). **b**, Immunoblotting of Histone H3 in the cytoplasmic and nuclear fractions (top panel). Total proteins per lane (bottom panels) were used as loading control.

**Supplementary Table S1. NanoString ratio data.**

**Supplementary Table S2. Oligonucleotides used for ChIP-qPCR.**

**Supplementary Table S3. F statistics for ANOVA.**
