## Supplemental Table 3 for "DRP1 induces neuroinflammation *via* transcriptional regulation of NF-ĸB"

| Panel | Mitochondrial morphology quantification | Two-way ANOVA |
|---|---|---|
| b | Mean Rod/Branch Length | Treatment |
| | | Genotpye |
| | | Interaction |
| | Mean Branches per network | Treatment |
| | | Genotpye |
| | | Interaction |
| | Aspect Ratio | Treatment |
| | | Genotpye |
| | | Interaction |
| | Roundness | Treatment |
| | | Genotpye |
| | | Interaction |

| Panel | Mitochondrial respiration quantification | Two-way ANOVA |
|---|---|---|
| d | Basal Respiration | Treatment |
| | | Genotpye |
| | | Interaction |
| | Spare Respiratory Capacity % | Treatment |
| | | Genotpye |
| | | Interaction |
| e | Basal Respiration | Treatment |
| | | Genotpye |
| | | Interaction |
| | Spare Respiratory Capacity % | Treatment |
| | | Genotpye |
| | | Interaction |
| f | Basal Respiration | Treatment |
| | | Genotpye |
| | | Interaction |
| | Spare Respiratory Capacity % | Treatment |
| | | Genotpye |
| | | Interaction |

| F (DFn, DFd) | p value |
|---|---|
| F (3, 24) = 20.05 | p<0.0001 |
| F (1, 24) = 9.363 | p=0.0054 |
| F (3, 24) = 0.9424 | p=0.4357 |
| F (3, 24) = 6.848 | p=0.0017 |
| F (1, 24) = 2.043 | p=0.1658 |
| F (3, 24) = 1.844 | p=0.1662 |
| F (3, 24) = 3.737 | p=0.0246 |
| F (1, 24) = 0.6184 | p=0.4393 |
| F (3, 24) = 0.5765 | p=0.6360 |
| F (3, 24) = 7.373 | p=0.0011 |
| F (1, 24) = 6.321 | p=0.0190 |
| F (3, 24) = 0.4178 | p=0.7418 |

| F (DFn, DFd) | p value |
|---|---|
| F (1, 8) = 1.066 | p=0.3320 |
| F (1, 8) = 0.007385 | p=0.9336 |
| F (1, 8) = 2.354 | p=0.1635 |
| F (1, 8) = 4.811 | p=0.0596 |
| F (1, 8) = 0.06412 | p=0.8065 |
| F (1, 8) = 0.05683 | p=0.8176 |
| F (1, 16) = 0.006571 | p=0.9364 |
| F (1, 16) = 1.689 | p=0.2121 |
| F (1, 16) = 0.09005 | p=0.7680 |
| F (1, 12) = 1.566 | p=0.2346 |
| F (1, 12) = 0.051 | p=0.8251 |
| F (1, 12) = 0.07898 | p=0.7835 |
| F (1, 12) = 6.434 | p=0.0261 |
| F (1, 12) = 0.2827 | p=0.6046 |
| F (1, 12) = 2.790 | p=0.1207 |
| F (1, 16) = 9.369 | p=0.0075 |
| F (1, 16) = 0.1016 | p=0.7540 |
| F (1, 16) = 0.005007 | p=0.9445 |
