## Supplemental Table 2 for "DRP1 induces neuroinflammation *via* transcriptional regulation of NF-ĸB"

**Table S2. Oligonucleotides used for ChIP-qPCR, related to STAR Methods**

| Gene Name | Primer Name | Forward Primer | Reverse Primer | Size (bp) |
|---|---|---|---|---|
| *Lcn2* | − | 5'-GAG TGG ACA GGC AGT CCA GAT C-3' | 5'-AAG ATT TCT GTC CCT CTC TCC CCC-3' | 170 |
| *Cxcl10* | − | 5'-CAT CTG ATT TCT CAA ACA GCT CAC G-3' | 5'-GGC TGC TGA GGA GTA TTT ATC TG-3' | 243 |
| *Il6* | − | 5'-CCT CAA GGA TGA CTT AAG CAC AC-3' | 5'-ATT AGG AGT CAA CTC TCT AAT TTT GAG ACT C-3' | 199 |
| *Rela* | P1 | 5'-CTG CAT CCC CTT CGT TTT C-3' | 5'-GAT AAA TGT GAG TTG TGT TTG GTA G-3' | 336 |
| | P2 | 5'-CCT GCA CTA CCA AAC ACA AC-3' | 5'-TCA GAC GTT CAA CGT CAT C-3' | 266 |
| | P3 | 5'-ATG ACG TTG AAC GTC TGA C-3' | 5'-CAG GAA GAC AAA AGT GAA AAT AGT TAC-3' | 356 |
| | P4 | 5'-GCG TAA CTA TTT TCA CTT TTG TCT TC-3' | 5'-CCG TGA ACA TCT CCT TCA AG-3' | 325 |
| | P5 | 5'-GGA GAT GTT CAC GGT GTG-3' | 5'-CAG TGA CTG AAT TCC ACA CG-3' | 317 |
| | P6 | 5'-ACG CTT AGG AAA ACG TGT G-3' | 5'-GAC TAC AAG CTC CGC AG-3' | 280 |
| | P7 | 5'-TGC GGA GCT TGT AGT CG-3' | 5'-GCT AAA GTA AAG CCA TTC GCC-3' | 255 |
