## Supplemental Table 1 for "DRP1 induces neuroinflammation *via* transcriptional regulation of NF-ĸB"

| Probe Name | Accession # | Class Name | Analyte Type | WT-LPS | StDev of WT-LPS |
|---|---|---|---|---|---|
| AI464131 | NM_001085515.2 | Endogenous | mRNA | 15.05 | 7.2 |
| Abcc3 | NM_029600.3 | Endogenous | mRNA | 2.89 | 1.56 |
| Abcc8 | NM_011510.3 | Endogenous | mRNA | 10.8 | 2.5 |
| Abl1 | NM_009594.4 | Endogenous | mRNA | 38.9 | 5.62 |
| Adamts16 | NM_172053.2 | Endogenous | mRNA | 10.32 | 6.01 |
| Ago4 | NM_153177.3 | Endogenous | mRNA | 5.08 | 2.93 |
| Agt | NM_007428.3 | Endogenous | mRNA | 107.36 | 19.52 |
| Ak1 | NM_001198790.1 | Endogenous | mRNA | 374.73 | 253.45 |
| Akt1 | NM_001165894.1 | Endogenous | mRNA | 111.8 | 14.75 |
| Akt2 | NM_001110208.1 | Endogenous | mRNA | 38.14 | 24.19 |
| Aldh1l1 | NM_027406.1 | Endogenous | mRNA | 79.19 | 12.36 |
| Ambra1 | NM_001080754.1 | Endogenous | mRNA | 193.63 | 87.04 |
| Amigo2 | NM_178114.4 | Endogenous | mRNA | 53.01 | 13.04 |
| Anapc15 | NM_027532.3 | Endogenous | mRNA | 35.55 | 8.81 |
| Anxa1 | NM_010730.2 | Endogenous | mRNA | 7.81 | 6.3 |
| Apc | NM_007462.3 | Endogenous | mRNA | 362.57 | 112.66 |
| Apex1 | NM_009687.2 | Endogenous | mRNA | 12.47 | 8.7 |
| Apoe | NM_001305844.1 | Endogenous | mRNA | 742.65 | 155.93 |
| Arc | NM_018790.2 | Endogenous | mRNA | 18.39 | 10.3 |
| Arhgap24 | NM_029270.2 | Endogenous | mRNA | 31.07 | 16.28 |
| Arid1a | NM_001080819.1 | Endogenous | mRNA | 81.11 | 35.63 |
| Asb2 | NM_023049.1 | Endogenous | mRNA | 3.39 | 2.15 |
| Ash2l | NM_001080793.1 | Endogenous | mRNA | 94.09 | 16.76 |
| Asph | NM_001177849.1 | Endogenous | mRNA | 201.81 | 35.79 |
| Atf3 | NM_007498.3 | Endogenous | mRNA | 4.05 | 2.14 |
| Atg14 | NM_172599.4 | Endogenous | mRNA | 48.2 | 24.34 |
| Atg3 | NM_026402.3 | Endogenous | mRNA | 223.16 | 61.64 |
| Atg5 | NM_001314013.1 | Endogenous | mRNA | 101.77 | 48.62 |
| Atg7 | NM_028835.1 | Endogenous | mRNA | 24.13 | 7.72 |
| Atg9a | NM_001003917.3 | Endogenous | mRNA | 197.32 | 95 |
| Atm | NM_007499.2 | Endogenous | mRNA | 39.97 | 20.06 |
| Atp6v0e | NM_025272.2 | Endogenous | mRNA | 81.07 | 47.68 |
| Atp6v1a | NM_007508.5 | Endogenous | mRNA | 422.17 | 88.36 |
| Atr | NM_019864.1 | Endogenous | mRNA | 40.66 | 33.01 |
| Axl | NM_009465.3 | Endogenous | mRNA | 76.94 | 47.79 |
| B3gnt5 | NM_001159407.1 | Endogenous | mRNA | 3.52 | 2.7 |
| Bad | NM_007522.3 | Endogenous | mRNA | 54.31 | 21.47 |
| Bag3 | NM_013863.4 | Endogenous | mRNA | 20.05 | 9.89 |
| Bag4 | NM_026121.3 | Endogenous | mRNA | 92.81 | 25.27 |
| Bak1 | NM_007523.2 | Endogenous | mRNA | 54.37 | 29.51 |
| Bard1 | NM_007525.3 | Endogenous | mRNA | 8.59 | 4.38 |
| Bax | NM_007527.3 | Endogenous | mRNA | 18.26 | 7.05 |
| Bbc3 | NM_133234.1 | Endogenous | mRNA | 7.59 | 2.07 |
| Bcas1 | NM_029815.2 | Endogenous | mRNA | 779.57 | 265.53 |
| Bcl10 | NM_009740.1 | Endogenous | mRNA | 33.45 | 7.74 |
| Bcl2 | NM_009741.3 | Endogenous | mRNA | 10.26 | 5.8 |

| | | | | | |
|---|---|---|---|---|---|
| Bcl2a1a | NM_009742.3 | Endogenous | mRNA | 63.94 | 41.84 |
| Bcl2l1 | NM_009743.4 | Endogenous | mRNA | 242.35 | 96.69 |
| Bcl2l11 | NM_001284410.1 | Endogenous | mRNA | 4.75 | 3.14 |
| Bcl2l2 | NM_007537.1 | Endogenous | mRNA | 373.99 | 243.14 |
| Bdnf | NM_007540.4 | Endogenous | mRNA | 15.5 | 8.07 |
| Becn1 | NM_019584.3 | Endogenous | mRNA | 262.44 | 87 |
| Bid | NM_007544.3 | Endogenous | mRNA | 37.47 | 7.95 |
| Bik | NM_007546.2 | Endogenous | mRNA | 2.87 | 1.56 |
| Bin1 | NM_001083334.1 | Endogenous | mRNA | 292.72 | 180.19 |
| Birc2 | NM_007465.2 | Endogenous | mRNA | 91.86 | 32.31 |
| Birc3 | NM_007464.3 | Endogenous | mRNA | 6.92 | 1.62 |
| Birc5 | NM_009689.2 | Endogenous | mRNA | 7 | 3.07 |
| Blk | NM_007549.2 | Endogenous | mRNA | 3.71 | 2.64 |
| Blm | NM_001042527.2 | Endogenous | mRNA | 19.42 | 19.94 |
| Blnk | NM_008528.4 | Endogenous | mRNA | 3.89 | 1.89 |
| Bmi1 | NM_007552.4 | Endogenous | mRNA | 36.55 | 16.73 |
| Bnip3 | NM_009760.4 | Endogenous | mRNA | 209.63 | 69.67 |
| Bnip3l | NM_009761.3 | Endogenous | mRNA | 244.6 | 77.41 |
| Bok | NM_016778.2 | Endogenous | mRNA | 13.45 | 4.04 |
| Bola2 | NM_175103.3 | Endogenous | mRNA | 331.68 | 119.73 |
| Braf | NM_139294.5 | Endogenous | mRNA | 148.22 | 56.3 |
| Brca1 | NM_009764.3 | Endogenous | mRNA | 3.61 | 1.56 |
| Brd2 | NM_010238.3 | Endogenous | mRNA | 167.38 | 24.61 |
| Brd3 | NM_001113573.1 | Endogenous | mRNA | 54.12 | 16.07 |
| Brd4 | NM_001286630.1 | Endogenous | mRNA | 131.96 | 52.73 |
| Btk | NM_013482.2 | Endogenous | mRNA | 3.67 | 1.65 |
| C1qa | NM_007572.2 | Endogenous | mRNA | 93.77 | 50.05 |
| C1qb | NM_009777.2 | Endogenous | mRNA | 241.3 | 140.26 |
| C1qc | NM_007574.2 | Endogenous | mRNA | 81.03 | 48.62 |
| C3 | XM_011246258.1 | Endogenous | mRNA | 13.73 | 9.15 |
| C3ar1 | NM_009779.2 | Endogenous | mRNA | 22.04 | 12.54 |
| C4a | NM_011413.2 | Endogenous | mRNA | 103.58 | 44.97 |
| C5ar1 | NM_007577.3 | Endogenous | mRNA | 25.32 | 10.9 |
| C6 | NM_016704.2 | Endogenous | mRNA | 3.3 | 2.19 |
| Cables1 | NM_001146287.1 | Endogenous | mRNA | 85.16 | 21.51 |
| Calcoco2 | NM_001271018.1 | Endogenous | mRNA | 3.09 | 1.65 |
| Calr | NM_007591.3 | Endogenous | mRNA | 1174.65 | 589.4 |
| Camk4 | NM_009793.3 | Endogenous | mRNA | 54.52 | 13.33 |
| Casp1 | NM_009807.2 | Endogenous | mRNA | 7.3 | 4.65 |
| Casp2 | NM_007610.1 | Endogenous | mRNA | 20.53 | 9.75 |
| Casp3 | NM_009810.2 | Endogenous | mRNA | 17.06 | 5.32 |
| Casp4 | NM_007609.2 | Endogenous | mRNA | 21.56 | 10.88 |
| Casp6 | NM_009811.3 | Endogenous | mRNA | 2.89 | 1.56 |
| Casp7 | NM_007611.2 | Endogenous | mRNA | 33.47 | 27.54 |
| Casp8 | NM_009812.2 | Endogenous | mRNA | 4.3 | 2.24 |
| Casp9 | NM_015733.4 | Endogenous | mRNA | 25.56 | 17.82 |
| Cass4 | NM_001080820.2 | Endogenous | mRNA | 3.97 | 2.45 |

| | | | | | |
|---|---|---|---|---|---|
| Ccl2 | NM_011333.3 | Endogenous | mRNA | 211.36 | 264.93 |
| Ccl3 | NM_011337.1 | Endogenous | mRNA | 17.52 | 11.53 |
| Ccl4 | NM_013652.1 | Endogenous | mRNA | 5.12 | 3.15 |
| Ccl5 | NM_013653.1 | Endogenous | mRNA | 20.97 | 15.49 |
| Ccl7 | NM_013654.3 | Endogenous | mRNA | 16.44 | 14.68 |
| Ccng2 | NM_007635.4 | Endogenous | mRNA | 44.55 | 25.76 |
| Ccni | NM_017367.3 | Endogenous | mRNA | 190.58 | 46.28 |
| Ccr2 | NM_009915.2 | Endogenous | mRNA | 7.72 | 7.43 |
| Ccr5 | NM_009917.5 | Endogenous | mRNA | 4.93 | 2.17 |
| Cd109 | NM_153098.3 | Endogenous | mRNA | 6.35 | 1.89 |
| Cd14 | NM_009841.3 | Endogenous | mRNA | 24.83 | 8.85 |
| Cd163 | NM_053094.2 | Endogenous | mRNA | 2.87 | 1.56 |
| Cd19 | NM_009844.2 | Endogenous | mRNA | 3.61 | 1.56 |
| Cd209e | NM_130905.2 | Endogenous | mRNA | 5.9 | 1.96 |
| Cd244 | NM_018729.2 | Endogenous | mRNA | 14.65 | 11.08 |
| Cd24a | NM_009846.2 | Endogenous | mRNA | 52.69 | 21.17 |
| Cd300lf | NM_001169153.1 | Endogenous | mRNA | 6.33 | 4.09 |
| Cd33 | NM_001111058.1 | Endogenous | mRNA | 4.9 | 3.96 |
| Cd36 | NM_007643.3 | Endogenous | mRNA | 5.22 | 1.87 |
| Cd3d | NM_013487.2 | Endogenous | mRNA | 4.36 | 2.26 |
| Cd3e | NM_007648.4 | Endogenous | mRNA | 2.87 | 1.56 |
| Cd3g | NM_009850.2 | Endogenous | mRNA | 6.02 | 3.68 |
| Cd40 | NM_011611.2 | Endogenous | mRNA | 13.22 | 10.85 |
| Cd44 | NM_009851.2 | Endogenous | mRNA | 3.3 | 2.19 |
| Cd47 | NM_010581.3 | Endogenous | mRNA | 323.38 | 132.58 |
| Cd6 | NM_001037801.2 | Endogenous | mRNA | 3.61 | 1.56 |
| Cd68 | NM_009853.1 | Endogenous | mRNA | 14.32 | 13.69 |
| Cd69 | NM_001033122.3 | Endogenous | mRNA | 5.49 | 4.4 |
| Cd70 | NM_011617.1 | Endogenous | mRNA | 15.62 | 19.89 |
| Cd72 | NM_001110320.1 | Endogenous | mRNA | 11.82 | 13.13 |
| Cd74 | NM_001042605.1 | Endogenous | mRNA | 32.21 | 28.64 |
| Cd83 | NM_009856.2 | Endogenous | mRNA | 192.84 | 71.8 |
| Cd84 | NM_013489.2 | Endogenous | mRNA | 10.87 | 13.09 |
| Cd86 | NM_019388.3 | Endogenous | mRNA | 9.15 | 1.64 |
| Cd8a | NM_001081110.2 | Endogenous | mRNA | 2.87 | 1.56 |
| Cd8b1 | NM_009858.2 | Endogenous | mRNA | 3.86 | 1.87 |
| Cdc25a | NM_007658.3 | Endogenous | mRNA | 22.42 | 6.22 |
| Cdc7 | NM_001271566.1 | Endogenous | mRNA | 23.35 | 8.52 |
| Cdk20 | NM_053180.2 | Endogenous | mRNA | 23.9 | 10.41 |
| Cdkn1a | NM_007669.4 | Endogenous | mRNA | 341.48 | 148.37 |
| Cdkn1c | NM_009876.3 | Endogenous | mRNA | 5.12 | 4.4 |
| Ceacam3 | NM_054059.1 | Endogenous | mRNA | 2.87 | 1.56 |
| Cflar | NM_207653.3 | Endogenous | mRNA | 28.02 | 9.59 |
| Ch25h | NM_009890.1 | Endogenous | mRNA | 8.62 | 2.88 |
| Chek1 | NM_007691.5 | Endogenous | mRNA | 10.77 | 5.2 |
| Chek2 | NM_016681.3 | Endogenous | mRNA | 13.05 | 6.81 |
| Chn2 | NM_001163640.1 | Endogenous | mRNA | 52.16 | 16.53 |

| | | | | | |
|---|---|---|---|---|---|
| Chst8 | NM_175140.4 | Endogenous | mRNA | 32.42 | 14.66 |
| Chuk | NM_001162410.1 | Endogenous | mRNA | 75.54 | 13.49 |
| Cidea | NM_007702.2 | Endogenous | mRNA | 3.36 | 2.66 |
| Cideb | NM_009894.3 | Endogenous | mRNA | 23.38 | 28.36 |
| Cks1b | NM_016904.1 | Endogenous | mRNA | 9.39 | 5.35 |
| Clcf1 | NM_019952.3 | Endogenous | mRNA | 8.58 | 5.57 |
| Cldn5 | NM_013805.4 | Endogenous | mRNA | 40.47 | 18.17 |
| Clec7a | NM_020008.2 | Endogenous | mRNA | 6.95 | 3.28 |
| Clic4 | NM_013885.2 | Endogenous | mRNA | 320.78 | 218.23 |
| Cln3 | NM_001146311.1 | Endogenous | mRNA | 22.59 | 7.84 |
| Clstn1 | NM_023051.4 | Endogenous | mRNA | 271.28 | 66.97 |
| Cnn2 | NM_007725.2 | Endogenous | mRNA | 17.52 | 13.15 |
| Cnp | NM_009923.2 | Endogenous | mRNA | 908.39 | 281.11 |
| Cntnap2 | NM_001004357.2 | Endogenous | mRNA | 93.43 | 17.57 |
| Coa5 | NM_198006.4 | Endogenous | mRNA | 119.06 | 58.24 |
| Col6a3 | XM_897036.2 | Endogenous | mRNA | 15.49 | 16.91 |
| Cotl1 | NM_028071.3 | Endogenous | mRNA | 125.2 | 34.11 |
| Cox5b | NM_009942.2 | Endogenous | mRNA | 1294.57 | 423.04 |
| Cp | NM_001042611.1 | Endogenous | mRNA | 158.92 | 58.29 |
| Cpa3 | NM_007753.2 | Endogenous | mRNA | 2.87 | 1.56 |
| Creb1 | NM_001037726.1 | Endogenous | mRNA | 36.81 | 17.85 |
| Crebbp | NM_001025432.1 | Endogenous | mRNA | 122.26 | 40.31 |
| Crem | NM_001110853.1 | Endogenous | mRNA | 20.56 | 4.49 |
| Crip1 | NM_007763.3 | Endogenous | mRNA | 16.56 | 3.68 |
| Cryba4 | NM_021351.1 | Endogenous | mRNA | 3.64 | 1.82 |
| Csf1 | NM_001113530.1 | Endogenous | mRNA | 160.81 | 72.12 |
| Csf1r | NM_001037859.1 | Endogenous | mRNA | 60.09 | 25.36 |
| Csf2rb | NM_007780.4 | Endogenous | mRNA | 4.67 | 4.88 |
| Csf3r | NM_001252651.1 | Endogenous | mRNA | 5.22 | 3.06 |
| Csk | NM_007783.2 | Endogenous | mRNA | 36.21 | 12.61 |
| Cst7 | NM_009977.3 | Endogenous | mRNA | 4.23 | 1.91 |
| Ctse | NM_007799.3 | Endogenous | mRNA | 6.04 | 4.95 |
| Ctsf | NM_019861.1 | Endogenous | mRNA | 86.53 | 36.86 |
| Ctss | NM_021281.2 | Endogenous | mRNA | 107.19 | 53.85 |
| Ctsw | NM_009985.4 | Endogenous | mRNA | 2.87 | 1.56 |
| Cx3cl1 | NM_009142.3 | Endogenous | mRNA | 55.31 | 29.42 |
| Cx3cr1 | NM_009987.3 | Endogenous | mRNA | 46.7 | 16.82 |
| Cxcl10 | NM_021274.1 | Endogenous | mRNA | 125.08 | 108.38 |
| Cxcl9 | NM_008599.2 | Endogenous | mRNA | 13.34 | 9.45 |
| Cycs | NM_007808.4 | Endogenous | mRNA | 4.77 | 2.78 |
| Cyp27a1 | NM_024264.3 | Endogenous | mRNA | 33.16 | 26.03 |
| Cyp7b1 | NM_007825.4 | Endogenous | mRNA | 22.93 | 11.85 |
| Cytip | NM_139200.4 | Endogenous | mRNA | 8.96 | 13.78 |
| Dab2 | NM_023118.2 | Endogenous | mRNA | 13.95 | 12.59 |
| Dapk1 | NM_134062.1 | Endogenous | mRNA | 48.29 | 9.72 |
| Ddb2 | NM_028119.5 | Endogenous | mRNA | 21.56 | 10.93 |
| Ddx58 | NM_172689.3 | Endogenous | mRNA | 96.92 | 84.5 |

| | | | | | |
|---|---|---|---|---|---|
| Dicer1 | NM_148948.2 | Endogenous | mRNA | 46.46 | 6.8 |
| Dlg1 | NM_001252433.1 | Endogenous | mRNA | 132.14 | 67.29 |
| Dlg4 | NM_001109752.1 | Endogenous | mRNA | 322.42 | 162.6 |
| Dlx1 | NM_010053.1 | Endogenous | mRNA | 5.71 | 3.84 |
| Dlx2 | NM_010054.2 | Endogenous | mRNA | 5.73 | 6.37 |
| Dna2 | NM_177372.3 | Endogenous | mRNA | 7.03 | 3.54 |
| Dnmt1 | NM_010066.3 | Endogenous | mRNA | 20.66 | 6.85 |
| Dnmt3a | NM_007872.4 | Endogenous | mRNA | 45.31 | 27.8 |
| Dnmt3b | NM_001003960.3 | Endogenous | mRNA | 3.09 | 1.65 |
| Dock1 | NM_001033420.2 | Endogenous | mRNA | 66.09 | 16.78 |
| Dock2 | NM_033374.3 | Endogenous | mRNA | 4.73 | 1.89 |
| Dot1l | NM_199322.1 | Endogenous | mRNA | 11.07 | 10.04 |
| Dst | NM_010081.2 | Endogenous | mRNA | 247.47 | 87.85 |
| Duoxa1 | NM_145395.2 | Endogenous | mRNA | 3.54 | 1.59 |
| Dusp7 | NM_153459.4 | Endogenous | mRNA | 64.64 | 19.17 |
| E2f1 | NM_007891.4 | Endogenous | mRNA | 13.01 | 10.68 |
| Eed | NM_021876.3 | Endogenous | mRNA | 75.39 | 34.53 |
| Eef2k | NM_007908.3 | Endogenous | mRNA | 48.77 | 17.7 |
| Egfr | NM_207655.2 | Endogenous | mRNA | 12.15 | 6.65 |
| Egr1 | NM_007913.5 | Endogenous | mRNA | 62.05 | 19.24 |
| Ehmt2 | NM_145830.1 | Endogenous | mRNA | 144.61 | 31.53 |
| Eif1 | NM_011508.1 | Endogenous | mRNA | 530.59 | 146.83 |
| Emcn | NM_001163522.1 | Endogenous | mRNA | 9.33 | 6.51 |
| Emp1 | NM_010128.4 | Endogenous | mRNA | 18.86 | 11.11 |
| Enpp6 | NM_177304.3 | Endogenous | mRNA | 18.31 | 16.25 |
| Entpd2 | NM_009849.2 | Endogenous | mRNA | 11.45 | 3.32 |
| Eomes | NM_010136.2 | Endogenous | mRNA | 4.55 | 4.01 |
| Ep300 | NM_177821.6 | Endogenous | mRNA | 127.13 | 58.77 |
| Epcam | NM_008532.2 | Endogenous | mRNA | 3.86 | 1.87 |
| Epg5 | NM_001195633.1 | Endogenous | mRNA | 25.94 | 10.31 |
| Epsti1 | NM_029495.2 | Endogenous | mRNA | 3.29 | 1.46 |
| Erbb3 | NM_010153.1 | Endogenous | mRNA | 15.48 | 16.33 |
| Ercc2 | NM_007949.4 | Endogenous | mRNA | 64.86 | 35.1 |
| Esam | NM_027102.3 | Endogenous | mRNA | 53.87 | 28.87 |
| Ets2 | NM_011809.2 | Endogenous | mRNA | 58.71 | 13.41 |
| Exo1 | NM_012012.4 | Endogenous | mRNA | 2.87 | 1.56 |
| Ezh1 | NM_007970.1 | Endogenous | mRNA | 36.98 | 15.47 |
| Ezh2 | NM_007971.2 | Endogenous | mRNA | 44.53 | 25.26 |
| F3 | NM_010171.3 | Endogenous | mRNA | 209.57 | 51.07 |
| Fa2h | NM_178086.3 | Endogenous | mRNA | 44.02 | 24.07 |
| Fabp5 | NM_010634.3 | Endogenous | mRNA | 307.6 | 82.76 |
| Fadd | NM_010175.5 | Endogenous | mRNA | 4.51 | 2.52 |
| Fancc | NM_007985.2 | Endogenous | mRNA | 12.84 | 6.94 |
| Fancd2 | NM_001033244.3 | Endogenous | mRNA | 4.27 | 3.71 |
| Fancg | NM_053081.2 | Endogenous | mRNA | 16.05 | 6.7 |
| Fas | NM_007987.2 | Endogenous | mRNA | 13.78 | 6.07 |
| Fasl | NM_010177.3 | Endogenous | mRNA | 2.87 | 1.56 |

| | | | | | |
|---|---|---|---|---|---|
| Fbln5 | NM_011812.4 | Endogenous | mRNA | 14.5 | 8.91 |
| Fcer1g | NM_010185.4 | Endogenous | mRNA | 19.49 | 5.77 |
| Fcgr1 | NM_010186.5 | Endogenous | mRNA | 28.12 | 23.65 |
| Fcgr2b | NM_001077189.1 | Endogenous | mRNA | 74.12 | 26.55 |
| Fcgr3 | NM_010188.5 | Endogenous | mRNA | 9.21 | 6.09 |
| Fcrla | NM_145141.2 | Endogenous | mRNA | 6.87 | 1.78 |
| Fcrlb | NM_001029984.2 | Endogenous | mRNA | 4.36 | 2.26 |
| Fcrls | NM_030707.3 | Endogenous | mRNA | 19.95 | 14.25 |
| Fdxr | NM_007997.1 | Endogenous | mRNA | 4.58 | 2.16 |
| Fen1 | NM_001271614.1 | Endogenous | mRNA | 20.1 | 6.07 |
| Fgd2 | NM_001159538.1 | Endogenous | mRNA | 3.71 | 2.64 |
| Fgf13 | NM_010200.2 | Endogenous | mRNA | 189.09 | 57.33 |
| Fgl2 | NM_008013.2 | Endogenous | mRNA | 16.61 | 16.8 |
| Fkbp5 | NM_010220.3 | Endogenous | mRNA | 88.11 | 45.37 |
| Flt1 | NM_010228.3 | Endogenous | mRNA | 6.89 | 2.09 |
| Fos | NM_010234.2 | Endogenous | mRNA | 34 | 11.86 |
| Foxp3 | NM_054039.2 | Endogenous | mRNA | 3.59 | 2.35 |
| Fpr1 | NM_013521.2 | Endogenous | mRNA | 3.93 | 2.49 |
| Fscn1 | NM_007984.2 | Endogenous | mRNA | 100.75 | 18.02 |
| Fyn | NM_008054.2 | Endogenous | mRNA | 148.81 | 50.87 |
| Gadd45a | NM_007836.1 | Endogenous | mRNA | 69.3 | 35.68 |
| Gadd45g | NM_011817.2 | Endogenous | mRNA | 43.69 | 22.85 |
| Gal3st1 | NM_001177691.1 | Endogenous | mRNA | 12.65 | 1.57 |
| Gba | NM_001077411.1 | Endogenous | mRNA | 37.36 | 15.06 |
| Gbp2 | NM_010260.1 | Endogenous | mRNA | 162.56 | 117.49 |
| Gclc | NM_010295.2 | Endogenous | mRNA | 27.88 | 10.75 |
| Gdpd2 | NM_023608.3 | Endogenous | mRNA | 34.27 | 8.15 |
| Gja1 | NM_010288.3 | Endogenous | mRNA | 807.43 | 412.83 |
| Gjb1 | NM_008124.2 | Endogenous | mRNA | 61.34 | 20.77 |
| Gna15 | NM_010304.3 | Endogenous | mRNA | 7.89 | 5.12 |
| Gpr183 | NM_183031.2 | Endogenous | mRNA | 8.14 | 6.44 |
| Gpr34 | NM_011823.4 | Endogenous | mRNA | 2.87 | 1.56 |
| Gpr62 | NM_001159652.1 | Endogenous | mRNA | 34.59 | 8.71 |
| Gpr84 | NM_030720.1 | Endogenous | mRNA | 22.5 | 8.41 |
| Grap | NM_027817.3 | Endogenous | mRNA | 5.64 | 3.07 |
| Gria1 | NM_001252403.1 | Endogenous | mRNA | 133.01 | 25.98 |
| Gria2 | NM_001039195.1 | Endogenous | mRNA | 359.69 | 116.48 |
| Gria4 | NM_001113180.1 | Endogenous | mRNA | 128.05 | 60.38 |
| Grin2a | NM_008170.2 | Endogenous | mRNA | 7.61 | 3.65 |
| Grin2b | NM_008171.3 | Endogenous | mRNA | 104.79 | 55.85 |
| Grm2 | NM_001160353.1 | Endogenous | mRNA | 11.06 | 3.85 |
| Grm3 | NM_181850.2 | Endogenous | mRNA | 56.44 | 10.38 |
| Grn | NM_008175.3 | Endogenous | mRNA | 31.83 | 13.39 |
| Gsn | NM_146120.3 | Endogenous | mRNA | 132.99 | 43.81 |
| Gstm1 | NM_010358.5 | Endogenous | mRNA | 15.71 | 9.61 |
| Gzma | NM_010370.2 | Endogenous | mRNA | 3.3 | 2.19 |
| Gzmb | NM_013542.2 | Endogenous | mRNA | 2.87 | 1.56 |

| | | | | | |
|---|---|---|---|---|---|
| H2-T23 | NM_010398.3 | Endogenous | mRNA | 8.08 | 7.01 |
| H2afx | NM_010436.2 | Endogenous | mRNA | 89.66 | 29.85 |
| Hat1 | NM_026115.4 | Endogenous | mRNA | 106.44 | 46.34 |
| Hcar2 | NM_030701.1 | Endogenous | mRNA | 5.64 | 5.22 |
| Hdac1 | NM_008228.2 | Endogenous | mRNA | 29.29 | 10.77 |
| Hdac2 | NM_008229.2 | Endogenous | mRNA | 132.09 | 42.07 |
| Hdac4 | NM_207225.1 | Endogenous | mRNA | 23.64 | 6.6 |
| Hdac6 | NM_010413.3 | Endogenous | mRNA | 34.2 | 6.24 |
| Hdc | NM_008230.4 | Endogenous | mRNA | 3.21 | 1.87 |
| Hells | NM_008234.3 | Endogenous | mRNA | 12.41 | 5.27 |
| Hif1a | NM_010431.2 | Endogenous | mRNA | 262.31 | 61.62 |
| Hilpda | NM_023516.5 | Endogenous | mRNA | 3.53 | 1.44 |
| Hira | NM_010435.2 | Endogenous | mRNA | 41.05 | 9.64 |
| Hist1h1d | NM_145713.3 | Endogenous | mRNA | 5.33 | 2.49 |
| Hmgb1 | NM_010439.3 | Endogenous | mRNA | 176.95 | 67.45 |
| Hmox1 | NM_010442.2 | Endogenous | mRNA | 11.8 | 7.11 |
| Homer1 | NM_147176.2 | Endogenous | mRNA | 63.25 | 14.21 |
| Hpgds | NM_019455.4 | Endogenous | mRNA | 8.16 | 5.73 |
| Hprt | NM_013556.2 | Endogenous | mRNA | 95.26 | 27.64 |
| Hps4 | NM_138646.3 | Endogenous | mRNA | 4.44 | 3.16 |
| Hrk | NM_007545.2 | Endogenous | mRNA | 17.1 | 16.31 |
| Hsd11b1 | NM_008288.2 | Endogenous | mRNA | 126 | 47.97 |
| Hspb1 | NM_013560.2 | Endogenous | mRNA | 86.13 | 40.12 |
| Hus1 | NM_008316.2 | Endogenous | mRNA | 28.71 | 8.24 |
| Icam2 | NM_010494.1 | Endogenous | mRNA | 14.86 | 21.03 |
| Ifi30 | NM_023065.3 | Endogenous | mRNA | 15.57 | 7.32 |
| Ifih1 | NM_027835.2 | Endogenous | mRNA | 26.51 | 13.39 |
| Ifitm2 | NM_030694.1 | Endogenous | mRNA | 129.72 | 84.82 |
| Ifitm3 | NM_025378.2 | Endogenous | mRNA | 249.77 | 74.01 |
| Ifnar1 | NM_010508.1 | Endogenous | mRNA | 69.42 | 28.86 |
| Ifnar2 | NM_001110498.1 | Endogenous | mRNA | 35.91 | 17.64 |
| Igf1 | NM_001111274.1 | Endogenous | mRNA | 6.42 | 4.08 |
| Igf1r | NM_010513.2 | Endogenous | mRNA | 22.52 | 10.57 |
| Igf2r | NM_010515.1 | Endogenous | mRNA | 34.39 | 27.71 |
| Igsf10 | NM_001162884.1 | Endogenous | mRNA | 18.65 | 2.47 |
| Igsf6 | NM_030691.1 | Endogenous | mRNA | 12.91 | 7.8 |
| Ikbkb | NM_010546.2 | Endogenous | mRNA | 52.69 | 17.12 |
| Ikbke | NM_019777.3 | Endogenous | mRNA | 4.59 | 7.81 |
| Ikbkg | NM_178590.2 | Endogenous | mRNA | 32.93 | 20.17 |
| Il10rb | NM_008349.5 | Endogenous | mRNA | 8.25 | 4.14 |
| Il15ra | NM_008358.2 | Endogenous | mRNA | 9.84 | 3.97 |
| Il1a | NM_010554.4 | Endogenous | mRNA | 15.39 | 13.05 |
| Il1b | NM_008361.3 | Endogenous | mRNA | 13.14 | 8.8 |
| Il1r1 | NM_001123382.1 | Endogenous | mRNA | 24.89 | 7.02 |
| Il1r2 | NM_010555.4 | Endogenous | mRNA | 4.67 | 2.33 |
| Il1rap | NM_134103.2 | Endogenous | mRNA | 50.65 | 16.61 |
| Il1rl2 | NM_133193.3 | Endogenous | mRNA | 8.27 | 3.76 |

| | | | | | |
|---|---|---|---|---|---|
| Il1rn | NM_031167.5 | Endogenous | mRNA | 12.12 | 2.9 |
| Il21r | NM_021887.1 | Endogenous | mRNA | 6.17 | 5.44 |
| Il2rg | NM_013563.3 | Endogenous | mRNA | 14.65 | 6.7 |
| Il3 | NM_010556.4 | Endogenous | mRNA | 8.51 | 5.3 |
| Il3ra | NM_008369.1 | Endogenous | mRNA | 3.52 | 1.87 |
| Il6ra | NM_010559.2 | Endogenous | mRNA | 14.16 | 4.37 |
| Inpp5d | NM_001110192.1 | Endogenous | mRNA | 9.65 | 6.44 |
| Iqsec1 | NM_001134383.1 | Endogenous | mRNA | 527.42 | 257.78 |
| Irak1 | NM_008363.2 | Endogenous | mRNA | 22.3 | 7.71 |
| Irak2 | NM_001113553.1 | Endogenous | mRNA | 45.3 | 23.83 |
| Irak3 | NM_028679.3 | Endogenous | mRNA | 11.68 | 3.17 |
| Irak4 | NM_029926.5 | Endogenous | mRNA | 34.37 | 29.15 |
| Irf1 | NM_008390.1 | Endogenous | mRNA | 49.77 | 16.64 |
| Irf2 | NM_008391.2 | Endogenous | mRNA | 39.49 | 15.04 |
| Irf3 | NM_016849.4 | Endogenous | mRNA | 17.45 | 8.61 |
| Irf4 | NM_013674.1 | Endogenous | mRNA | 5.14 | 2.55 |
| Irf6 | NM_016851.2 | Endogenous | mRNA | 5.71 | 4.26 |
| Irf7 | NM_016850.2 | Endogenous | mRNA | 50.83 | 31.62 |
| Irf8 | NM_008320.3 | Endogenous | mRNA | 5.33 | 2.87 |
| Islr2 | NM_001161538.1 | Endogenous | mRNA | 4.49 | 1.79 |
| Itga6 | NM_008397.3 | Endogenous | mRNA | 52.35 | 25.14 |
| Itga7 | NM_008398.2 | Endogenous | mRNA | 18.26 | 8.61 |
| Itgam | NM_001082960.1 | Endogenous | mRNA | 13.59 | 6 |
| Itgav | NM_008402.2 | Endogenous | mRNA | 55.66 | 14.2 |
| Itgax | NM_021334.2 | Endogenous | mRNA | 3.43 | 2.74 |
| Itgb5 | NM_001145884.1 | Endogenous | mRNA | 95.59 | 45.73 |
| Jag1 | NM_013822.2 | Endogenous | mRNA | 12.39 | 2.47 |
| Jam2 | NM_023844.4 | Endogenous | mRNA | 113.17 | 54.58 |
| Jarid2 | NM_021878.2 | Endogenous | mRNA | 39.18 | 13.79 |
| Jun | NM_010591.2 | Endogenous | mRNA | 69.67 | 21.51 |
| Kat2a | NM_020004.5 | Endogenous | mRNA | 62.06 | 25.44 |
| Kat2b | NM_020005.3 | Endogenous | mRNA | 78.2 | 23.51 |
| Kcnd1 | NM_008423.1 | Endogenous | mRNA | 10.1 | 6.07 |
| Kcnj10 | NM_001039484.1 | Endogenous | mRNA | 383.26 | 187.49 |
| Kcnk13 | NM_146037.1 | Endogenous | mRNA | 7.75 | 8.52 |
| Kdm1a | NM_133872.1 | Endogenous | mRNA | 120.57 | 59.72 |
| Kdm1b | NM_172262.3 | Endogenous | mRNA | 14.63 | 4.2 |
| Kdm2a | NM_001001984.2 | Endogenous | mRNA | 117.76 | 73.18 |
| Kdm2b | NM_001003953.1 | Endogenous | mRNA | 9.93 | 5.65 |
| Kdm3a | NM_001038695.2 | Endogenous | mRNA | 47.38 | 16.71 |
| Kdm3b | NM_001081256.1 | Endogenous | mRNA | 2.87 | 1.56 |
| Kdm4a | NM_172382.2 | Endogenous | mRNA | 20.81 | 6.46 |
| Kdm4b | NM_172132.1 | Endogenous | mRNA | 57.01 | 19.54 |
| Kdm4c | NM_144787.1 | Endogenous | mRNA | 29.6 | 9.94 |
| Kdm4d | NM_173433.2 | Endogenous | mRNA | 8.54 | 5.08 |
| Kdm5a | XR_377436.1 | Endogenous | mRNA | 83.93 | 29.13 |
| Kdm5b | NM_152895.2 | Endogenous | mRNA | 70.04 | 23.78 |

| | | | | | |
|---|---|---|---|---|---|
| Kdm5c | NM_013668.3 | Endogenous | mRNA | 130.76 | 81.35 |
| Kdm5d | NM_011419.3 | Endogenous | mRNA | 14.98 | 16.01 |
| Kdm6a | NM_009483.1 | Endogenous | mRNA | 14.35 | 5.35 |
| Kif2c | NM_134471.3 | Endogenous | mRNA | 3.54 | 1.47 |
| Kir3dl1 | NM_177749.3 | Endogenous | mRNA | 2.87 | 1.56 |
| Kir3dl2 | NM_177748.2 | Endogenous | mRNA | 2.87 | 1.56 |
| Kit | NM_001122733.1 | Endogenous | mRNA | 13.97 | 5.66 |
| Klrb1 | NM_001099918.1 | Endogenous | mRNA | 6.03 | 4.46 |
| Klrd1 | NM_010654.2 | Endogenous | mRNA | 5.26 | 3.69 |
| Klrk1 | NM_001083322.1 | Endogenous | mRNA | 10.44 | 10.5 |
| Kmt2a | NM_001081049.1 | Endogenous | mRNA | 86.29 | 13.44 |
| Kmt2c | NM_001081383.1 | Endogenous | mRNA | 88.67 | 35.02 |
| Lacc1 | NM_172488.2 | Endogenous | mRNA | 14.34 | 9.43 |
| Lag3 | NM_008479.1 | Endogenous | mRNA | 12.71 | 9.39 |
| Lair1 | NM_001113474.1 | Endogenous | mRNA | 10.6 | 6.07 |
| Lamp1 | NM_010684.2 | Endogenous | mRNA | 791.18 | 324.38 |
| Lamp2 | NM_001017959.1 | Endogenous | mRNA | 345.76 | 87.69 |
| Lcn2 | NM_008491.1 | Endogenous | mRNA | 1440.61 | 518.84 |
| Ldha | NM_010699.2 | Endogenous | mRNA | 237.68 | 96.82 |
| Ldlrad3 | NM_178886.2 | Endogenous | mRNA | 33.69 | 23.81 |
| Lfng | NM_008494.3 | Endogenous | mRNA | 86.9 | 17.74 |
| Lgmn | NM_011175.3 | Endogenous | mRNA | 225.53 | 46.65 |
| Lig1 | NM_001083188.1 | Endogenous | mRNA | 22.39 | 9.33 |
| Lilrb4a | NM_013532.3 | Endogenous | mRNA | 28.48 | 16.83 |
| Lingo1 | NM_181074.4 | Endogenous | mRNA | 63.01 | 8.49 |
| Lmna | NM_001002011.2 | Endogenous | mRNA | 71.45 | 22.28 |
| Lmnb1 | NM_010721.2 | Endogenous | mRNA | 15.08 | 4.26 |
| Lrg1 | NM_029796.2 | Endogenous | mRNA | 4.23 | 2.64 |
| Lrrc25 | NM_153074.3 | Endogenous | mRNA | 9.29 | 8.73 |
| Lrrc3 | NM_145152.4 | Endogenous | mRNA | 10.81 | 3.27 |
| Lsr | NM_001164184.1 | Endogenous | mRNA | 10.65 | 3.25 |
| Lst1 | NM_010734.2 | Endogenous | mRNA | 7.29 | 7.4 |
| Lta | NM_010735.2 | Endogenous | mRNA | 10.03 | 7.44 |
| Ltb | NM_008518.2 | Endogenous | mRNA | 4.36 | 2.59 |
| Ltbr | NM_010736.3 | Endogenous | mRNA | 11.39 | 4.3 |
| Ltc4s | NM_008521.1 | Endogenous | mRNA | 2.96 | 1.56 |
| Ly6a | NM_010738.2 | Endogenous | mRNA | 295.56 | 102.08 |
| Ly6g | XM_909927.2 | Endogenous | mRNA | 3.17 | 1.66 |
| Ly9 | NM_008534.2 | Endogenous | mRNA | 11.04 | 11.27 |
| Lyn | NM_010747.1 | Endogenous | mRNA | 15.19 | 8.21 |
| Mafb | NM_010658.2 | Endogenous | mRNA | 32.03 | 19.9 |
| Maff | NM_010755.3 | Endogenous | mRNA | 16.22 | 8.29 |
| Mag | NM_010758.2 | Endogenous | mRNA | 143.07 | 35.17 |
| Mal | NM_001171187.1 | Endogenous | mRNA | 530.8 | 301.58 |
| Man2b1 | NM_010764.2 | Endogenous | mRNA | 69.06 | 31.6 |
| Map1lc3a | NM_025735.1 | Endogenous | mRNA | 191.29 | 49.04 |
| Map2k1 | NM_008927.3 | Endogenous | mRNA | 195.69 | 31.33 |

| | | | | | |
|---|---|---|---|---|---|
| Map2k4 | NM_009157.4 | Endogenous | mRNA | 314.81 | 77.92 |
| Map3k1 | NM_011945.2 | Endogenous | mRNA | 17.43 | 5.81 |
| Map3k14 | NM_016896.3 | Endogenous | mRNA | 4.67 | 3.4 |
| Mapk10 | NM_001081567.1 | Endogenous | mRNA | 401.06 | 108.54 |
| Mapk12 | NM_013871.3 | Endogenous | mRNA | 3.3 | 2.19 |
| Mapk14 | NM_011951.2 | Endogenous | mRNA | 31.14 | 12.59 |
| Mapt | NM_001038609.2 | Endogenous | mRNA | 200.62 | 49.65 |
| Marco | NM_010766.2 | Endogenous | mRNA | 4.67 | 2.16 |
| Mavs | NM_144888.2 | Endogenous | mRNA | 26.66 | 10.45 |
| Mb21d1 | NM_173386.4 | Endogenous | mRNA | 3.79 | 1.93 |
| Mbd2 | NM_010773.2 | Endogenous | mRNA | 168.01 | 26.34 |
| Mbd3 | NM_013595.2 | Endogenous | mRNA | 97.33 | 34.4 |
| Mcm2 | NM_008564.2 | Endogenous | mRNA | 8.69 | 3.92 |
| Mcm5 | NM_008566.2 | Endogenous | mRNA | 13.78 | 13.04 |
| Mcm6 | NM_008567.1 | Endogenous | mRNA | 5.81 | 3.03 |
| Mdc1 | NM_001010833.2 | Endogenous | mRNA | 12.91 | 1.94 |
| Mdm2 | NM_010786.4 | Endogenous | mRNA | 119.83 | 56.2 |
| Mef2c | NM_001170537.1 | Endogenous | mRNA | 26.24 | 9.34 |
| Mertk | NM_008587.1 | Endogenous | mRNA | 46.18 | 18.82 |
| Mfge8 | NM_008594.2 | Endogenous | mRNA | 24.68 | 11.98 |
| Mgmt | NM_008598.2 | Endogenous | mRNA | 10.01 | 6.93 |
| Mmp12 | NM_008605.3 | Endogenous | mRNA | 7.72 | 5.26 |
| Mmp14 | NM_008608.3 | Endogenous | mRNA | 9.71 | 6.77 |
| Mobp | NM_001039364.2 | Endogenous | mRNA | 1520.49 | 455.1 |
| Mog | NM_010814.2 | Endogenous | mRNA | 95.31 | 17.2 |
| Mpeg1 | NM_010821.1 | Endogenous | mRNA | 32.46 | 16.14 |
| Mpg | NM_010822.3 | Endogenous | mRNA | 10.01 | 6.36 |
| Mr1 | NM_008209.4 | Endogenous | mRNA | 14.61 | 8.15 |
| Mre11a | NM_018736.2 | Endogenous | mRNA | 27.2 | 7.78 |
| Ms4a1 | NM_007641.5 | Endogenous | mRNA | 4.44 | 4.86 |
| Ms4a2 | NM_001276330.1 | Endogenous | mRNA | 2.87 | 1.56 |
| Ms4a4a | XM_003086124.1 | Endogenous | mRNA | 18.16 | 11.49 |
| Msh2 | NM_008628.2 | Endogenous | mRNA | 70.63 | 16.22 |
| Msn | NM_010833.2 | Endogenous | mRNA | 31.3 | 18.72 |
| Msr1 | NM_001113326.1 | Endogenous | mRNA | 21.51 | 14.11 |
| Mvp | NM_080638.2 | Endogenous | mRNA | 43.3 | 16.46 |
| Myc | NM_010849.4 | Endogenous | mRNA | 19.46 | 3.26 |
| Myct1 | NM_026793.2 | Endogenous | mRNA | 26.72 | 46.47 |
| Myd88 | NM_010851.2 | Endogenous | mRNA | 20.51 | 5.59 |
| Myrf | NM_001033481.1 | Endogenous | mRNA | 194.75 | 49.45 |
| Nbn | NM_013752.3 | Endogenous | mRNA | 7.86 | 4.32 |
| Ncaph | NM_144818.3 | Endogenous | mRNA | 8.63 | 12.04 |
| Ncf1 | NM_001286037.1 | Endogenous | mRNA | 28.11 | 14.9 |
| Ncor1 | NM_011308.2 | Endogenous | mRNA | 87.38 | 18.31 |
| Ncor2 | NM_011424.2 | Endogenous | mRNA | 147.02 | 58.15 |
| Ncr1 | NM_010746.3 | Endogenous | mRNA | 3.29 | 1.46 |
| Nefl | NM_010910.1 | Endogenous | mRNA | 383.75 | 173.52 |

| | | | | | |
|---|---|---|---|---|---|
| Nfe2l2 | NR_132727.1 | Endogenous | mRNA | 15.87 | 11.53 |
| Nfkb1 | NM_008689.2 | Endogenous | mRNA | 57.41 | 19.83 |
| Nfkb2 | NM_019408.2 | Endogenous | mRNA | 19.16 | 18.34 |
| Nfkbia | NM_010907.2 | Endogenous | mRNA | 39.37 | 17.63 |
| Nfkbie | NM_008690.3 | Endogenous | mRNA | 8.44 | 6.77 |
| Ngf | NM_001112698.1 | Endogenous | mRNA | 13.41 | 10.25 |
| Ngfr | NM_033217.3 | Endogenous | mRNA | 3.97 | 2.2 |
| Ninj2 | NM_016718.2 | Endogenous | mRNA | 4.55 | 4.01 |
| Nkg7 | NM_024253.4 | Endogenous | mRNA | 4.89 | 9.6 |
| Nlgn1 | NM_138666.3 | Endogenous | mRNA | 105.97 | 21.3 |
| Nlgn2 | NM_198862.2 | Endogenous | mRNA | 246.99 | 67.16 |
| Nlrp3 | NM_145827.3 | Endogenous | mRNA | 9.46 | 2.88 |
| Nod1 | NM_172729.2 | Endogenous | mRNA | 16.39 | 12.45 |
| Nostrin | NM_181547.3 | Endogenous | mRNA | 10.13 | 4.37 |
| Npl | NM_028749.1 | Endogenous | mRNA | 9.61 | 5.84 |
| Npnt | NM_001029836.1 | Endogenous | mRNA | 11.2 | 8.72 |
| Nptx1 | NM_008730.2 | Endogenous | mRNA | 97.64 | 47.94 |
| Nqo1 | NM_008706.5 | Endogenous | mRNA | 21.37 | 11.3 |
| Nrgn | NM_022029.2 | Endogenous | mRNA | 14.81 | 8.23 |
| Nrm | NM_134122.2 | Endogenous | mRNA | 4.3 | 2.04 |
| Nrp2 | NM_001077403.1 | Endogenous | mRNA | 23.4 | 14.03 |
| Nthl1 | NM_008743.2 | Endogenous | mRNA | 9.05 | 3.33 |
| Nwd1 | NM_176940.5 | Endogenous | mRNA | 70.49 | 29.38 |
| Oas1g | NM_011852.2 | Endogenous | mRNA | 9.38 | 5.54 |
| Ogg1 | NM_010957.4 | Endogenous | mRNA | 13.73 | 10.72 |
| Olfml3 | NM_133859.2 | Endogenous | mRNA | 22.62 | 6.59 |
| Opalin | NM_153520.1 | Endogenous | mRNA | 59.78 | 26.23 |
| Optn | NM_181848.4 | Endogenous | mRNA | 97.06 | 24.31 |
| Osgin1 | NM_027950.1 | Endogenous | mRNA | 5.27 | 4.36 |
| Osmr | NM_011019.3 | Endogenous | mRNA | 105.4 | 45.37 |
| P2rx7 | NM_001038839.2 | Endogenous | mRNA | 8.99 | 3.71 |
| P2ry12 | NM_027571.3 | Endogenous | mRNA | 16.07 | 12.28 |
| Pacsin1 | NM_011861.3 | Endogenous | mRNA | 444.22 | 108.97 |
| Padi2 | NM_008812.2 | Endogenous | mRNA | 28.69 | 2.93 |
| Pak1 | NM_011035.2 | Endogenous | mRNA | 552.13 | 190.58 |
| Parp1 | NM_007415.2 | Endogenous | mRNA | 56.34 | 17.04 |
| Parp2 | NM_009632.2 | Endogenous | mRNA | 36.04 | 12.6 |
| Pcna | NM_011045.2 | Endogenous | mRNA | 107.84 | 27.77 |
| Pdpn | NM_010329.2 | Endogenous | mRNA | 66.13 | 11.73 |
| Pecam1 | NM_008816.2 | Endogenous | mRNA | 28.63 | 13.55 |
| Pex14 | NM_019781.2 | Endogenous | mRNA | 67.86 | 8.61 |
| Pik3ca | NM_008839.1 | Endogenous | mRNA | 55.26 | 13.72 |
| Pik3cb | NM_029094.3 | Endogenous | mRNA | 81.67 | 34.04 |
| Pik3cd | XM_003945690.1 | Endogenous | mRNA | 20.45 | 9.93 |
| Pik3cg | NM_020272.2 | Endogenous | mRNA | 4.25 | 2.48 |
| Pik3r1 | NM_001024955.1 | Endogenous | mRNA | 173.97 | 112.27 |
| Pik3r2 | NM_008841.2 | Endogenous | mRNA | 113.49 | 61.23 |

| | | | | | |
|---|---|---|---|---|---|
| Pik3r5 | NM_177320.2 | Endogenous | mRNA | 4.19 | 2.52 |
| Pilra | NM_153510.3 | Endogenous | mRNA | 12.1 | 12.38 |
| Pilrb1 | NM_133209.2 | Endogenous | mRNA | 4.23 | 1.91 |
| Pink1 | NM_026880.2 | Endogenous | mRNA | 736.44 | 119.68 |
| Pla2g4a | NM_008869.2 | Endogenous | mRNA | 12.62 | 7.5 |
| Pla2g5 | NM_001122954.1 | Endogenous | mRNA | 8.6 | 3.7 |
| Plcg2 | NM_172285.1 | Endogenous | mRNA | 7.04 | 4.56 |
| Pld1 | NM_001164056.1 | Endogenous | mRNA | 40.87 | 28.61 |
| Pld2 | NM_008876.2 | Endogenous | mRNA | 12.11 | 6.59 |
| Plekhb1 | NM_001163184.1 | Endogenous | mRNA | 1675.8 | 370.67 |
| Plekhm1 | NM_183034.1 | Endogenous | mRNA | 11.42 | 8.08 |
| Pllp | NM_026385.3 | Endogenous | mRNA | 17.68 | 9.94 |
| Plp1 | NM_011123.2 | Endogenous | mRNA | 2573.27 | 475.85 |
| Plxdc2 | NM_026162.5 | Endogenous | mRNA | 81.12 | 27.05 |
| Plxnb3 | NM_019587.2 | Endogenous | mRNA | 30.95 | 12.62 |
| Pmp22 | NM_008885.2 | Endogenous | mRNA | 89.69 | 55.54 |
| Pms2 | NM_008886.2 | Endogenous | mRNA | 45.69 | 16.07 |
| Pnoc | NM_001205075.1 | Endogenous | mRNA | 7.87 | 6.83 |
| Pole | NM_011132.2 | Endogenous | mRNA | 9.26 | 6.42 |
| Ppfia4 | NM_001144855.1 | Endogenous | mRNA | 36.91 | 13.56 |
| Ppp3ca | NM_008913.4 | Endogenous | mRNA | 249.64 | 57.65 |
| Ppp3cb | NM_008914.2 | Endogenous | mRNA | 195.02 | 33.41 |
| Ppp3r1 | NM_024459.2 | Endogenous | mRNA | 273 | 141.98 |
| Ppp3r2 | NM_001004025.4 | Endogenous | mRNA | 10.3 | 4.64 |
| Prdx1 | NM_011034.4 | Endogenous | mRNA | 22.19 | 6.94 |
| Prf1 | NM_011073.2 | Endogenous | mRNA | 3.61 | 1.83 |
| Prkaca | NM_008854.3 | Endogenous | mRNA | 598.41 | 113.34 |
| Prkacb | NM_011100.3 | Endogenous | mRNA | 328.34 | 47.23 |
| Prkar1a | NM_021880.2 | Endogenous | mRNA | 796.18 | 157.1 |
| Prkar2a | NM_008924.2 | Endogenous | mRNA | 164.24 | 80.76 |
| Prkar2b | NM_011158.3 | Endogenous | mRNA | 115.48 | 39.86 |
| Prkce | NM_011104.2 | Endogenous | mRNA | 74.91 | 26.52 |
| Prkcq | NM_008859.2 | Endogenous | mRNA | 21.9 | 11.19 |
| Prkdc | NM_011159.2 | Endogenous | mRNA | 17.15 | 14.1 |
| Pros1 | NM_011173.2 | Endogenous | mRNA | 34.9 | 16.92 |
| Psen2 | NM_001128605.1 | Endogenous | mRNA | 38.02 | 32.75 |
| Psmb8 | NM_010724.2 | Endogenous | mRNA | 53.81 | 39.9 |
| Pten | NM_008960.2 | Endogenous | mRNA | 116.89 | 35.56 |
| Ptger3 | NM_011196.2 | Endogenous | mRNA | 5.12 | 5.63 |
| Ptger4 | NM_008965.1 | Endogenous | mRNA | 5.28 | 2.48 |
| Ptgs2 | NM_011198.3 | Endogenous | mRNA | 36.77 | 10.67 |
| Ptms | NM_026988.2 | Endogenous | mRNA | 225.01 | 27.33 |
| Ptpn6 | NM_013545.2 | Endogenous | mRNA | 6.9 | 4.77 |
| Ptprc | NM_011210.3 | Endogenous | mRNA | 7.92 | 6.78 |
| Pttg1 | NM_001131054.1 | Endogenous | mRNA | 142.82 | 32.17 |
| Ptx3 | NM_008987.3 | Endogenous | mRNA | 27.18 | 21.65 |
| Pycard | NM_023258.4 | Endogenous | mRNA | 3.29 | 1.46 |

| | | | | | |
|---|---|---|---|---|---|
| Rab6b | NM_173781.4 | Endogenous | mRNA | 514.95 | 106.88 |
| Rab7 | NM_009005.2 | Endogenous | mRNA | 625.99 | 191.46 |
| Rac1 | NM_009007.2 | Endogenous | mRNA | 354.27 | 70.84 |
| Rac2 | NM_009008.3 | Endogenous | mRNA | 4.13 | 2.02 |
| Rad1 | NM_011232.2 | Endogenous | mRNA | 28.28 | 12.46 |
| Rad17 | NM_001044371.1 | Endogenous | mRNA | 20.63 | 15.37 |
| Rad50 | NM_009012.2 | Endogenous | mRNA | 5.26 | 2.85 |
| Rad51 | NM_011234.4 | Endogenous | mRNA | 5.91 | 3.38 |
| Rad51b | NM_009014.3 | Endogenous | mRNA | 6.57 | 7.03 |
| Rad51c | NM_053269.3 | Endogenous | mRNA | 11.58 | 9.22 |
| Rad9a | NM_011237.2 | Endogenous | mRNA | 14.5 | 8.07 |
| Rag1 | NM_009019.2 | Endogenous | mRNA | 3.29 | 1.46 |
| Rala | NM_019491.5 | Endogenous | mRNA | 78.35 | 20.35 |
| Ralb | NM_022327.5 | Endogenous | mRNA | 41.32 | 7.65 |
| Rapgef3 | NM_001177810.1 | Endogenous | mRNA | 89.68 | 7.49 |
| Rb1cc1 | NM_009826.4 | Endogenous | mRNA | 144.19 | 52.82 |
| Rbfox3 | NM_001024931.2 | Endogenous | mRNA | 287.48 | 130.4 |
| Rela | NM_009045.4 | Endogenous | mRNA | 40.69 | 25.59 |
| Relb | NM_009046.2 | Endogenous | mRNA | 21.3 | 6.04 |
| Reln | NM_011261.2 | Endogenous | mRNA | 14.5 | 3.03 |
| Rgl1 | NM_016846.3 | Endogenous | mRNA | 40 | 15.1 |
| Rhoa | NM_016802.4 | Endogenous | mRNA | 190.75 | 32.67 |
| Ripk1 | NM_009068.3 | Endogenous | mRNA | 31.12 | 31.83 |
| Ripk2 | NM_138952.3 | Endogenous | mRNA | 13.06 | 5.42 |
| Rnf8 | NM_021419.2 | Endogenous | mRNA | 29.56 | 8.76 |
| Rpa1 | NM_026653.2 | Endogenous | mRNA | 124.94 | 37.23 |
| Rpl28 | NM_009081.2 | Endogenous | mRNA | 504.92 | 139.92 |
| Rpl29 | NM_009082.2 | Endogenous | mRNA | 333.87 | 94.07 |
| Rpl36al | NM_025589.4 | Endogenous | mRNA | 92.38 | 29.35 |
| Rpl9 | NM_011292.2 | Endogenous | mRNA | 824.68 | 335.37 |
| Rps10 | NM_025963.3 | Endogenous | mRNA | 73.59 | 20.01 |
| Rps2 | NM_008503.5 | Endogenous | mRNA | 613.4 | 193.15 |
| Rps21 | NM_025587.2 | Endogenous | mRNA | 1661.18 | 768.88 |
| Rps3 | NM_012052.2 | Endogenous | mRNA | 42.96 | 22.68 |
| Rps9 | NM_029767.2 | Endogenous | mRNA | 635.87 | 163.45 |
| Rrm2 | NM_009104.1 | Endogenous | mRNA | 9.29 | 7.04 |
| Rsad2 | NM_021384.2 | Endogenous | mRNA | 126.75 | 127.95 |
| Rtn4rl1 | NM_177708.4 | Endogenous | mRNA | 8.57 | 4.45 |
| S100a10 | NM_009112.2 | Endogenous | mRNA | 123.05 | 41.32 |
| S100b | NM_009115.3 | Endogenous | mRNA | 385.52 | 123.02 |
| S1pr3 | NM_010101.3 | Endogenous | mRNA | 14.36 | 7.37 |
| S1pr4 | NM_010102.2 | Endogenous | mRNA | 4.41 | 3.33 |
| S1pr5 | NM_053190.2 | Endogenous | mRNA | 25.17 | 13.13 |
| Sall1 | NM_021390.3 | Endogenous | mRNA | 38.4 | 8.78 |
| Sell | XM_006496716.1 | Endogenous | mRNA | 7.03 | 3.48 |
| Serpina3n | NM_009252.2 | Endogenous | mRNA | 91.98 | 32.96 |
| Serpine1 | NM_008871.2 | Endogenous | mRNA | 5.27 | 3.34 |

| | | | | | |
|---|---|---|---|---|---|
| Serpinf1 | NM_011340.3 | Endogenous | mRNA | 55.26 | 13.63 |
| Serping1 | NM_009776.3 | Endogenous | mRNA | 20 | 8.68 |
| Sesn1 | NM_001013370.2 | Endogenous | mRNA | 85.78 | 38.65 |
| Sesn2 | NM_144907.1 | Endogenous | mRNA | 12.2 | 4.5 |
| Setd1a | NM_178029.3 | Endogenous | mRNA | 43.55 | 13.42 |
| Setd1b | NM_001040398.1 | Endogenous | mRNA | 47.91 | 15.4 |
| Setd2 | NM_001081340.2 | Endogenous | mRNA | 103.41 | 12.97 |
| Setd7 | NM_080793.5 | Endogenous | mRNA | 59.2 | 10.02 |
| Setdb1 | NM_018877.2 | Endogenous | mRNA | 20.28 | 4.8 |
| Sftpd | NM_009160.2 | Endogenous | mRNA | 5.67 | 6.18 |
| Sh2d1a | NR_132588.1 | Endogenous | mRNA | 2.87 | 1.56 |
| Shank3 | NM_021423.3 | Endogenous | mRNA | 24.95 | 10.72 |
| Siglec1 | NM_011426.3 | Endogenous | mRNA | 5.24 | 2.52 |
| Siglecf | NM_145581.1 | Endogenous | mRNA | 8.28 | 6.16 |
| Sin3a | NM_001110350.1 | Endogenous | mRNA | 65.98 | 21.45 |
| Sirt1 | NM_019812.2 | Endogenous | mRNA | 44.71 | 29.05 |
| Slamf8 | NM_029084.3 | Endogenous | mRNA | 5.08 | 3.84 |
| Slamf9 | NM_029612.4 | Endogenous | mRNA | 4.25 | 2.33 |
| Slc10a6 | NM_029415.2 | Endogenous | mRNA | 9.76 | 6.12 |
| Slc17a6 | NM_080853.3 | Endogenous | mRNA | 353 | 95.48 |
| Slc17a7 | NM_182993.2 | Endogenous | mRNA | 119.12 | 62.26 |
| Slc1a3 | NM_148938.3 | Endogenous | mRNA | 353.66 | 64.5 |
| Slc2a1 | NM_011400.3 | Endogenous | mRNA | 55.47 | 17.59 |
| Slc2a5 | NM_019741.3 | Endogenous | mRNA | 5.96 | 2.8 |
| Slc44a1 | NM_001159633.1 | Endogenous | mRNA | 223.95 | 53.6 |
| Slc6a1 | NM_178703.4 | Endogenous | mRNA | 361.36 | 63.7 |
| Slco2b1 | NM_175316.3 | Endogenous | mRNA | 7.06 | 4.17 |
| Slfn8 | NM_181545.4 | Endogenous | mRNA | 21.71 | 19.12 |
| Smarca4 | NM_011417.2 | Endogenous | mRNA | 106.57 | 21 |
| Smarca5 | NM_053124.2 | Endogenous | mRNA | 240.18 | 47.31 |
| Smarcd1 | NM_031842.1 | Endogenous | mRNA | 18.87 | 11.93 |
| Smc1a | NM_019710.2 | Endogenous | mRNA | 36 | 12.96 |
| Snca | NM_009221.2 | Endogenous | mRNA | 45.86 | 14.25 |
| Socs3 | NM_007707.2 | Endogenous | mRNA | 7.86 | 2.46 |
| Sod2 | NM_013671.3 | Endogenous | mRNA | 416.76 | 114.14 |
| Sox10 | XM_128139.6 | Endogenous | mRNA | 20.6 | 3.1 |
| Sox4 | NM_009238.2 | Endogenous | mRNA | 19.52 | 18.91 |
| Sox9 | NM_011448.4 | Endogenous | mRNA | 109.75 | 66.65 |
| Sphk1 | NM_011451.3 | Endogenous | mRNA | 4.53 | 2.34 |
| Spib | NM_019866.1 | Endogenous | mRNA | 3.04 | 1.69 |
| Spint1 | NM_016907.3 | Endogenous | mRNA | 5.03 | 2.47 |
| Spp1 | NM_009263.3 | Endogenous | mRNA | 350.72 | 134.51 |
| Sqstm1 | NM_011018.2 | Endogenous | mRNA | 589.12 | 175.02 |
| Srgn | NM_011157.2 | Endogenous | mRNA | 144.12 | 91.05 |
| Srxn1 | NM_029688.4 | Endogenous | mRNA | 144.85 | 8.22 |
| St3gal6 | NM_018784.2 | Endogenous | mRNA | 33.01 | 17.9 |
| St8sia6 | NM_145838.1 | Endogenous | mRNA | 22.44 | 8.64 |

| | | | | | |
|---|---|---|---|---|---|
| Stat1 | NM_009283.3 | Endogenous | mRNA | 25.11 | 6.33 |
| Steap4 | NM_054098.3 | Endogenous | mRNA | 29.71 | 24.54 |
| Stmn1 | NM_019641.3 | Endogenous | mRNA | 1193.59 | 364.05 |
| Stx18 | NM_026959.2 | Endogenous | mRNA | 23.91 | 10.59 |
| Sumo1 | NM_009460.1 | Endogenous | mRNA | 414.59 | 157.89 |
| Suv39h1 | NM_011514.2 | Endogenous | mRNA | 32.51 | 7.21 |
| Suv39h2 | NM_022724.4 | Endogenous | mRNA | 19.29 | 10.85 |
| Suz12 | NM_199196.1 | Endogenous | mRNA | 157.82 | 74.86 |
| Syk | NM_001198977.1 | Endogenous | mRNA | 9.1 | 3.28 |
| Syn2 | NM_013681.1 | Endogenous | mRNA | 329.07 | 84.28 |
| Syp | NM_009305.2 | Endogenous | mRNA | 387.29 | 54.86 |
| Tarbp2 | NM_001253795.1 | Endogenous | mRNA | 20.3 | 8.68 |
| Tbc1d4 | NM_001081278.2 | Endogenous | mRNA | 20.7 | 7.84 |
| Tbr1 | NM_009322.3 | Endogenous | mRNA | 5.49 | 4.22 |
| Tbx21 | NM_019507.2 | Endogenous | mRNA | 7.6 | 3.45 |
| Tcirg1 | NM_001136091.1 | Endogenous | mRNA | 43.22 | 33.72 |
| Tcl1 | NM_009337.3 | Endogenous | mRNA | 2.94 | 1.56 |
| Tet1 | NM_027384.1 | Endogenous | mRNA | 33.15 | 11.69 |
| Tfg | NM_001252443.1 | Endogenous | mRNA | 297.52 | 121.25 |
| Tgfa | NM_031199.2 | Endogenous | mRNA | 22.05 | 4.13 |
| Tgfb1 | NM_011577.1 | Endogenous | mRNA | 8.57 | 2.45 |
| Tgfbr1 | NM_009370.2 | Endogenous | mRNA | 56.24 | 31.94 |
| Tgm1 | NM_001161714.1 | Endogenous | mRNA | 2.87 | 1.56 |
| Tgm2 | NM_009373.3 | Endogenous | mRNA | 49.91 | 17.42 |
| Tie1 | NM_011587.2 | Endogenous | mRNA | 5.09 | 2.38 |
| Timeless | NM_011589.1 | Endogenous | mRNA | 24.69 | 14.48 |
| Timp1 | NM_011593.2 | Endogenous | mRNA | 20.23 | 14.73 |
| Tle3 | NM_009389.2 | Endogenous | mRNA | 71.02 | 31.33 |
| Tlr2 | NM_011905.2 | Endogenous | mRNA | 19.26 | 18.28 |
| Tlr4 | NM_021297.2 | Endogenous | mRNA | 6.11 | 3.15 |
| Tlr7 | NM_133211.3 | Endogenous | mRNA | 6.88 | 4.42 |
| Tm4sf1 | NM_008536.3 | Endogenous | mRNA | 99.43 | 60.94 |
| Tmc7 | NM_172476.4 | Endogenous | mRNA | 28.51 | 7.53 |
| Tmcc3 | NM_172051.2 | Endogenous | mRNA | 96.58 | 43.99 |
| Tmem100 | NM_026433.2 | Endogenous | mRNA | 17.49 | 5.25 |
| Tmem119 | NM_146162.2 | Endogenous | mRNA | 41.92 | 26.57 |
| Tmem144 | NM_027495.4 | Endogenous | mRNA | 14.63 | 6.54 |
| Tmem173 | NM_028261.1 | Endogenous | mRNA | 4.29 | 3.54 |
| Tmem204 | NM_001001183.1 | Endogenous | mRNA | 15.35 | 15.73 |
| Tmem206 | NM_025864.3 | Endogenous | mRNA | 50.41 | 27.28 |
| Tmem37 | NM_019432.2 | Endogenous | mRNA | 3.52 | 1.87 |
| Tmem64 | NM_181401.3 | Endogenous | mRNA | 127.05 | 70.54 |
| Tmem88b | NM_001033394.3 | Endogenous | mRNA | 72.58 | 47.66 |
| Tnf | NM_013693.2 | Endogenous | mRNA | 3.83 | 3.31 |
| Tnfrsf10b | NM_020275.3 | Endogenous | mRNA | 9.4 | 6.41 |
| Tnfrsf11b | NM_008764.3 | Endogenous | mRNA | 9.67 | 6.7 |
| Tnfrsf12a | NM_001161746.1 | Endogenous | mRNA | 48.04 | 20.08 |

| | | | | | |
|---|---|---|---|---|---|
| Tnfrsf13c | NM_028075.2 | Endogenous | mRNA | 4.9 | 2.9 |
| Tnfrsf17 | NM_011608.1 | Endogenous | mRNA | 5.79 | 3.83 |
| Tnfrsf1a | NM_011609.2 | Endogenous | mRNA | 42.19 | 11.47 |
| Tnfrsf1b | NM_011610.3 | Endogenous | mRNA | 9.6 | 5.34 |
| Tnfrsf25 | NM_033042.3 | Endogenous | mRNA | 5.63 | 5.94 |
| Tnfrsf4 | NM_011659.2 | Endogenous | mRNA | 3.3 | 2.19 |
| Tnfsf10 | NM_009425.2 | Endogenous | mRNA | 11.62 | 9.04 |
| Tnfsf12 | NM_011614.3 | Endogenous | mRNA | 14.04 | 11.01 |
| Tnfsf13b | NM_033622.1 | Endogenous | mRNA | 10.46 | 8.97 |
| Tnfsf4 | NM_009452.2 | Endogenous | mRNA | 4.55 | 4.01 |
| Tnfsf8 | NM_009403.2 | Endogenous | mRNA | 4.79 | 2.06 |
| Top2a | NM_011623.2 | Endogenous | mRNA | 14.73 | 13.36 |
| Topbp1 | NM_176979.5 | Endogenous | mRNA | 27.11 | 15.16 |
| Tpd52 | NM_001025262.1 | Endogenous | mRNA | 232.93 | 69.93 |
| Tpsb2 | NM_010781.3 | Endogenous | mRNA | 4.58 | 2.16 |
| Tradd | NM_001033161.2 | Endogenous | mRNA | 15.94 | 19.75 |
| Traf1 | NM_009421.3 | Endogenous | mRNA | 7.06 | 5.31 |
| Traf2 | NM_009422.2 | Endogenous | mRNA | 31.91 | 17.09 |
| Traf3 | NM_011632.3 | Endogenous | mRNA | 62.68 | 21.61 |
| Traf6 | NM_009424.2 | Endogenous | mRNA | 31.83 | 23.93 |
| Trat1 | NM_198297.3 | Endogenous | mRNA | 5.75 | 4.54 |
| Trem1 | NM_021406.5 | Endogenous | mRNA | 6.47 | 6.24 |
| Trem2 | NM_031254.2 | Endogenous | mRNA | 7.49 | 4.88 |
| Trem3 | NM_021407.3 | Endogenous | mRNA | 3.56 | 1.46 |
| Trim47 | NM_001205081.1 | Endogenous | mRNA | 11.23 | 4.1 |
| Trp53 | NM_011640.1 | Endogenous | mRNA | 15.93 | 14.38 |
| Trp53bp2 | NM_173378.2 | Endogenous | mRNA | 40.28 | 12.83 |
| Trp73 | NM_011642.3 | Endogenous | mRNA | 7.06 | 5.85 |
| Trpa1 | NM_177781.4 | Endogenous | mRNA | 3.23 | 1.74 |
| Trpm4 | NM_175130.4 | Endogenous | mRNA | 8.99 | 4.09 |
| Tspan18 | NM_183180.2 | Endogenous | mRNA | 4.61 | 2.65 |
| Ttr | NM_013697.4 | Endogenous | mRNA | 8 | 1.6 |
| Tubb3 | NM_023279.2 | Endogenous | mRNA | 776.81 | 223.06 |
| Tubb4a | NM_009451.3 | Endogenous | mRNA | 177.82 | 39.75 |
| Txnrd1 | NM_015762.2 | Endogenous | mRNA | 291.24 | 140.69 |
| Tyrobp | NM_011662.2 | Endogenous | mRNA | 21.07 | 11.51 |
| Ugt8a | NM_011674.4 | Endogenous | mRNA | 268.7 | 191.67 |
| Ulk1 | NM_009469.3 | Endogenous | mRNA | 61.1 | 3.42 |
| Ung | NM_001040691.1 | Endogenous | mRNA | 11.37 | 5.3 |
| Uty | NM_009484.2 | Endogenous | mRNA | 15.42 | 14.07 |
| Vamp7 | NM_011515.4 | Endogenous | mRNA | 186.9 | 56.38 |
| Vav1 | NM_011691.4 | Endogenous | mRNA | 11.94 | 7.26 |
| Vegfa | NM_001025250.3 | Endogenous | mRNA | 50.41 | 11 |
| Vim | NM_011701.4 | Endogenous | mRNA | 41.66 | 10.32 |
| Vps4a | NM_126165.1 | Endogenous | mRNA | 90.43 | 17 |
| Vps4b | NM_009190.2 | Endogenous | mRNA | 50.12 | 39.08 |
| Was | NM_009515.2 | Endogenous | mRNA | 3.89 | 1.85 |

| | | | | | |
|---|---|---|---|---|---|
| Wdr5 | NM_080848.2 | Endogenous | mRNA | 25.32 | 9.89 |
| Xcl1 | NM_008510.1 | Endogenous | mRNA | 8.26 | 5.47 |
| Xiap | NM_009688.2 | Endogenous | mRNA | 180.56 | 23.67 |
| Xrcc6 | NM_010247.2 | Endogenous | mRNA | 21.63 | 4.74 |
| Zbp1 | NM_021394.2 | Endogenous | mRNA | 21.6 | 14.29 |
| Zfp367 | NM_175494.4 | Endogenous | mRNA | 48.49 | 4.26 |
| Aars | NM_146217.4 | Housekeeping | mRNA | 70.84 | 31.68 |
| Asb10 | NM_080444.4 | Housekeeping | mRNA | 2.87 | 1.56 |
| Ccdc127 | NM_024201.3 | Housekeeping | mRNA | 109.76 | 11.4 |
| Cnot10 | NM_153585.5 | Housekeeping | mRNA | 56.19 | 10.18 |
| Csnk2a2 | NM_009974.3 | Housekeeping | mRNA | 69.83 | 26.15 |
| Fam104a | NM_138598.5 | Housekeeping | mRNA | 50.35 | 28.36 |
| Gusb | NM_010368.1 | Housekeeping | mRNA | 11.9 | 6.74 |
| Lars | NM_134137.2 | Housekeeping | mRNA | 47.49 | 8.68 |
| Mto1 | NM_026658.2 | Housekeeping | mRNA | 16 | 7.28 |
| Supt7l | NM_028150.1 | Housekeeping | mRNA | 29.93 | 5.03 |
| Tada2b | NM_001170454.1 | Housekeeping | mRNA | 103.06 | 40.01 |
| Tbp | NM_013684.3 | Housekeeping | mRNA | 28.97 | 8.55 |
| Xpnpep1 | NM_133216.3 | Housekeeping | mRNA | 76.34 | 14.53 |

| WT-CTL | StDev of WT-CTL | WT-LPS vs. WT-CTL | P value of: WT-LPS \ | gKO-LPS | StDev of gk |
|---|---|---|---|---|---|
| 26.37 | 8.68 | -1.75 | 0.03084852 | 13.1 | 6.36 |
| 4.39 | 3.79 | -1.52 | 0.16629936 | 4.11 | 2.64 |
| 5.83 | 5.95 | 1.85 | 0.05915449 | 8.02 | 1.38 |
| 40.61 | 11.58 | -1.04 | 0.71182024 | 33.19 | 6.64 |
| 14.27 | 5.96 | -1.38 | 0.258773 | 9.54 | 2.92 |
| 5.45 | 5.26 | -1.07 | 0.84934193 | 8.4 | 4.62 |
| 122.62 | 32.5 | -1.14 | 0.25183392 | 95.77 | 29.58 |
| 454.41 | 214.9 | -1.21 | 0.57663518 | 250.03 | 194.61 |
| 134.14 | 28.97 | -1.2 | 0.08724573 | 103.33 | 25.44 |
| 43.23 | 21.18 | -1.13 | 0.726008 | 24.9 | 15.1 |
| 89.93 | 16.33 | -1.14 | 0.18187299 | 74.89 | 31.02 |
| 210.54 | 76.59 | -1.09 | 0.74412227 | 138.96 | 96.22 |
| 57.09 | 13.73 | -1.08 | 0.58581525 | 42.05 | 43.25 |
| 40.58 | 7.9 | -1.14 | 0.32236671 | 38.43 | 8.33 |
| 9.53 | 11.53 | -1.22 | 0.66106141 | 10.42 | 3.79 |
| 510.39 | 137.13 | -1.41 | 0.05603671 | 349.3 | 112.86 |
| 10.82 | 12.43 | 1.15 | 0.79200161 | 22.38 | 4.06 |
| 1008.81 | 156.02 | -1.36 | 0.01416343 | 703.95 | 182.55 |
| 32.48 | 18.3 | -1.77 | 0.11361358 | 13.94 | 7.34 |
| 45.16 | 15.55 | -1.45 | 0.09557721 | 28.36 | 8.22 |
| 95.49 | 35.5 | -1.18 | 0.47761032 | 62.35 | 34.37 |
| 3.5 | 1.75 | -1.03 | 0.91088647 | 4.11 | 2.64 |
| 112.85 | 27.39 | -1.2 | 0.13862789 | 88.41 | 37.35 |
| 225.36 | 35.65 | -1.12 | 0.2395336 | 228.5 | 41.35 |
| 3.02 | 1.01 | 1.34 | 0.25752336 | 7.08 | 6.64 |
| 57.74 | 27.25 | -1.2 | 0.49339446 | 35.28 | 15.97 |
| 273.61 | 67.54 | -1.23 | 0.20936722 | 206.6 | 37.59 |
| 120.37 | 40.04 | -1.18 | 0.41405094 | 96.1 | 22.55 |
| 26.82 | 16.31 | -1.11 | 0.581806 | 21.67 | 1.98 |
| 236.13 | 84.29 | -1.2 | 0.4488253 | 143.49 | 84.08 |
| 49.27 | 17.13 | -1.23 | 0.41042593 | 35.05 | 25.29 |
| 84.1 | 42.72 | -1.04 | 0.90750575 | 55.7 | 51.5 |
| 442.01 | 76.28 | -1.05 | 0.65650892 | 431.09 | 44.84 |
| 45.15 | 18.22 | -1.11 | 0.76871192 | 28.01 | 21.41 |
| 70.51 | 34.19 | 1.09 | 0.78799444 | 48.46 | 30.05 |
| 4.26 | 3.19 | -1.21 | 0.5596028 | 7.66 | 2.46 |
| 66.78 | 20.17 | -1.23 | 0.27000722 | 49.55 | 19.79 |
| 20.72 | 6.83 | -1.03 | 0.87339664 | 18.01 | 8.72 |
| 126.03 | 37.78 | -1.36 | 0.05766781 | 90.82 | 34.29 |
| 54.64 | 28.33 | -1.01 | 0.98633307 | 38.11 | 26.3 |
| 8.16 | 7.84 | 1.05 | 0.88162172 | 8.53 | 2.42 |
| 22.55 | 8.26 | -1.23 | 0.33840838 | 13.8 | 3.35 |
| 4.46 | 4.34 | 1.7 | 0.07003404 | 6.35 | 1.3 |
| 926.94 | 176.92 | -1.19 | 0.27075249 | 794.82 | 105.42 |
| 27.32 | 9.08 | 1.22 | 0.23791243 | 40.99 | 8.25 |
| 20.67 | 9.9 | -2.01 | 0.02541479 | 14.95 | 10.56 |

| | | | | | |
|---|---|---|---|---|---|
| 14.87 | 8.87 | 4.3 | 0.00220087 | 35.95 | 31.06 |
| 188.44 | 49.92 | 1.29 | 0.19415988 | 199.12 | 74.1 |
| 6.63 | 4.98 | -1.4 | 0.33458093 | 5.35 | 1.72 |
| 436.31 | 198.43 | -1.17 | 0.64250952 | 245.29 | 185.82 |
| 19.06 | 11.02 | -1.23 | 0.43074697 | 12.23 | 3.1 |
| 295.06 | 75.85 | -1.12 | 0.46239972 | 223.94 | 50.21 |
| 30.43 | 20.56 | 1.23 | 0.37680444 | 28.96 | 15.98 |
| 3.1 | 1.04 | -1.08 | 0.71323913 | 5.33 | 1.72 |
| 426.95 | 144.13 | -1.46 | 0.19427772 | 235.63 | 90.6 |
| 83.95 | 29.16 | 1.09 | 0.64119798 | 84.28 | 28.27 |
| 3.18 | 1.13 | 2.18 | 0.00032965 | 6.06 | 4.77 |
| 3.47 | 3.15 | 2.02 | 0.01605231 | 5.83 | 1.63 |
| 3.74 | 4.15 | -1.01 | 0.97597563 | 4.11 | 2.64 |
| 21.72 | 19.53 | -1.12 | 0.81342316 | 13.8 | 6.47 |
| 6.33 | 4.09 | -1.63 | 0.10302139 | 4.45 | 2.74 |
| 57.49 | 18.98 | -1.57 | 0.05924118 | 37.64 | 15.5 |
| 168.79 | 57.97 | 1.24 | 0.23897147 | 269.88 | 65.11 |
| 325.64 | 73.51 | -1.33 | 0.06717487 | 246.33 | 21.02 |
| 17.93 | 9.67 | -1.33 | 0.23935448 | 11.96 | 15.79 |
| 405.24 | 125.99 | -1.22 | 0.28459606 | 269.41 | 73.45 |
| 189.81 | 60.21 | -1.28 | 0.21290019 | 129.27 | 34.19 |
| 3.02 | 1.01 | 1.19 | 0.40905833 | 6.85 | 4.2 |
| 175.49 | 33.45 | -1.05 | 0.5915302 | 158.11 | 31.77 |
| 65.61 | 18.49 | -1.21 | 0.24679965 | 49.6 | 14.83 |
| 155.23 | 71.29 | -1.18 | 0.50481027 | 105.22 | 87.82 |
| 3.53 | 1.73 | 1.04 | 0.86800343 | 5.99 | 1.46 |
| 95.39 | 34.53 | -1.02 | 0.94077915 | 75.68 | 57.62 |
| 225.39 | 85.5 | 1.07 | 0.789469 | 179.37 | 95.71 |
| 97.93 | 24.55 | -1.21 | 0.39286238 | 61.22 | 22.37 |
| 9.22 | 6.62 | 1.49 | 0.22441362 | 14.9 | 5.44 |
| 7.37 | 9.72 | 2.99 | 0.02821302 | 17.24 | 5.59 |
| 71.13 | 63.13 | 1.46 | 0.22439995 | 65.2 | 24.38 |
| 10.46 | 7.56 | 2.42 | 0.01932696 | 18.09 | 16.01 |
| 2.99 | 1.01 | 1.1 | 0.70499188 | 4.11 | 2.64 |
| 90.83 | 41.99 | -1.07 | 0.72844374 | 71.57 | 39.53 |
| 2.99 | 1.01 | 1.03 | 0.88487029 | 4.11 | 2.64 |
| 1515.03 | 560.62 | -1.29 | 0.35037768 | 910.32 | 603.46 |
| 58.66 | 10.42 | -1.08 | 0.5847013 | 59.78 | 33.48 |
| 3.74 | 2.45 | 1.95 | 0.01902986 | 6.15 | 1.96 |
| 22.68 | 10.97 | -1.1 | 0.67671174 | 16.53 | 9.63 |
| 22.01 | 10.07 | -1.29 | 0.17785561 | 18.64 | 5.26 |
| 2.99 | 1.01 | 7.22 | 0.00002221 | 22.67 | 13.52 |
| 3.68 | 2.59 | -1.28 | 0.35665959 | 5.43 | 1.64 |
| 35.3 | 24.26 | -1.05 | 0.90782905 | 19.81 | 15.17 |
| 3.72 | 3.15 | 1.16 | 0.6376763 | 5.6 | 3.59 |
| 31.35 | 14.73 | -1.23 | 0.47456303 | 19.6 | 9.09 |
| 3.18 | 1.13 | 1.25 | 0.39972109 | 4.11 | 2.64 |

| | | | | | |
|---|---|---|---|---|---|
| 5.49 | 7.2 | 38.52 | 0.00012828 | 111.72 | 149.54 |
| 6.57 | 7.55 | 2.67 | 0.02917617 | 11.72 | 20.21 |
| 5.13 | 4.55 | -1 | 0.99517828 | 7.29 | 3.19 |
| 3.6 | 2.5 | 5.82 | 0.00029511 | 19.18 | 19.44 |
| 3.1 | 1.04 | 5.3 | 0.00327979 | 14.41 | 14.8 |
| 53.09 | 27.9 | -1.19 | 0.55165434 | 28.06 | 20.8 |
| 229.51 | 35.73 | -1.2 | 0.11798833 | 194.31 | 39.17 |
| 5.14 | 4.32 | 1.5 | 0.32795462 | 4.85 | 2.19 |
| 2.99 | 1.01 | 1.65 | 0.03322075 | 4.11 | 2.64 |
| 4.58 | 4.18 | 1.39 | 0.23717196 | 6.18 | 2.82 |
| 3.87 | 3.35 | 6.41 | 0.00000504 | 21.33 | 13.89 |
| 2.99 | 1.01 | -1.04 | 0.8469106 | 4.84 | 2.21 |
| 2.99 | 1.01 | 1.21 | 0.38351625 | 4.11 | 2.64 |
| 4.11 | 4.81 | 1.44 | 0.25321952 | 5.18 | 2.42 |
| 16.28 | 15.27 | -1.11 | 0.82385367 | 9.95 | 12.61 |
| 42.7 | 19.85 | 1.23 | 0.39183605 | 41.69 | 18.98 |
| 3.66 | 4.96 | 1.73 | 0.13803697 | 6.8 | 2.18 |
| 8.45 | 6.29 | -1.72 | 0.17419486 | 6.72 | 2.16 |
| 5.12 | 4.65 | 1.02 | 0.94625211 | 5.44 | 3.06 |
| 4.35 | 4 | 1 | 0.99063665 | 4.52 | 2.33 |
| 2.99 | 1.01 | -1.04 | 0.8469106 | 4.11 | 2.64 |
| 6.27 | 9.86 | -1.04 | 0.9283666 | 6.15 | 5.89 |
| 14.21 | 9.34 | -1.08 | 0.86629039 | 13.5 | 10.9 |
| 3.64 | 3.41 | -1.1 | 0.75399244 | 5.07 | 2.34 |
| 363.77 | 123.81 | -1.12 | 0.60124409 | 270.08 | 144.89 |
| 5.92 | 4.53 | -1.64 | 0.1187412 | 6.98 | 2.29 |
| 13.62 | 6.93 | 1.05 | 0.87778056 | 11.38 | 6.09 |
| 4.41 | 4.95 | 1.25 | 0.58676523 | 7.03 | 5.24 |
| 21.9 | 17.72 | -1.4 | 0.54461467 | 10.7 | 29.16 |
| 9.32 | 9.98 | 1.27 | 0.62541056 | 8.31 | 6.02 |
| 22.04 | 19.06 | 1.46 | 0.40625572 | 19.29 | 12.01 |
| 211.06 | 74.41 | -1.09 | 0.65121102 | 161.65 | 66.89 |
| 15.81 | 11.71 | -1.45 | 0.34308669 | 9.87 | 6.19 |
| 4.97 | 4.57 | 1.84 | 0.04666433 | 11.43 | 3.17 |
| 2.99 | 1.01 | -1.04 | 0.8469106 | 4.11 | 2.64 |
| 3.6 | 2.5 | 1.07 | 0.79839844 | 4.94 | 2.14 |
| 27.61 | 7.82 | -1.23 | 0.16274939 | 22.02 | 2.11 |
| 23.46 | 8.26 | -1 | 0.98025107 | 19.21 | 8.03 |
| 22.92 | 15.06 | 1.04 | 0.87670541 | 19.09 | 16.35 |
| 48.85 | 31.12 | 6.99 | 0.00003103 | 153.61 | 186.38 |
| 13.95 | 11.59 | -2.72 | 0.01829427 | 7.21 | 2.95 |
| 2.99 | 1.01 | -1.04 | 0.8469106 | 5.18 | 2.42 |
| 15.44 | 9.2 | 1.82 | 0.10339799 | 33.45 | 14.3 |
| 2.99 | 1.01 | 2.88 | 0.00010899 | 9.43 | 3.31 |
| 13.55 | 9.56 | -1.26 | 0.41629657 | 9.22 | 4.84 |
| 16.34 | 9.49 | -1.25 | 0.48208189 | 9.55 | 6.96 |
| 91.76 | 27.94 | -1.76 | 0.0071152 | 57.74 | 26.23 |

| | | | | | |
|---|---|---|---|---|---|
| 36.61 | 8.6 | -1.13 | 0.51460445 | 23.55 | 5.75 |
| 86.12 | 14.88 | -1.14 | 0.18537056 | 78.76 | 8.07 |
| 3.72 | 3.25 | -1.11 | 0.76500922 | 5.69 | 4.1 |
| 29.54 | 10.3 | -1.26 | 0.50717437 | 16.45 | 15.15 |
| 10.37 | 10.92 | -1.1 | 0.80641246 | 9.73 | 7.55 |
| 7.87 | 7.79 | 1.09 | 0.82653469 | 6.85 | 4.2 |
| 108.9 | 51.26 | -2.69 | 0.00057332 | 43.77 | 7.78 |
| 2.99 | 1.01 | 2.33 | 0.00416947 | 6.98 | 3.07 |
| 229.46 | 92.23 | 1.4 | 0.31607372 | 246.45 | 98.26 |
| 22.09 | 6.16 | 1.02 | 0.88306659 | 17.09 | 6.37 |
| 309.92 | 67.32 | -1.14 | 0.30707687 | 235.01 | 72.7 |
| 19.31 | 15.5 | -1.1 | 0.78005141 | 12.24 | 7.02 |
| 1241.44 | 168.07 | -1.37 | 0.03256321 | 955.58 | 111.74 |
| 102.72 | 15.83 | -1.1 | 0.30648288 | 86.5 | 12.92 |
| 153.14 | 60.86 | -1.29 | 0.36445394 | 87.23 | 67.12 |
| 18.72 | 11.7 | -1.21 | 0.69321156 | 11.74 | 11.27 |
| 130.44 | 44.1 | -1.04 | 0.80306739 | 86.06 | 15.08 |
| 1565.79 | 306.94 | -1.21 | 0.20380482 | 1214.16 | 270.88 |
| 33.29 | 10.58 | 4.77 | 0.00001088 | 124.19 | 76.65 |
| 2.99 | 1.01 | -1.04 | 0.8469106 | 4.11 | 2.64 |
| 45.41 | 14.78 | -1.23 | 0.3486678 | 28.92 | 8.89 |
| 148.98 | 47.8 | -1.22 | 0.30236182 | 106.36 | 59.96 |
| 20.41 | 7 | 1.01 | 0.95999676 | 21.53 | 5.11 |
| 35.82 | 16.62 | -2.16 | 0.00223824 | 20.92 | 7.14 |
| 4.44 | 3.78 | -1.22 | 0.49527276 | 5.24 | 2.18 |
| 94.37 | 43.81 | 1.7 | 0.05043283 | 129.23 | 58.08 |
| 75.86 | 28.83 | -1.26 | 0.33051664 | 45.04 | 21.69 |
| 5.99 | 5.36 | -1.28 | 0.54381531 | 5.71 | 1.92 |
| 10.42 | 10.39 | -2 | 0.07419259 | 9.96 | 5.97 |
| 45.76 | 11.81 | -1.26 | 0.14538264 | 35.25 | 11.14 |
| 3.5 | 1.75 | 1.21 | 0.45480853 | 4.52 | 2.33 |
| 4.28 | 3.94 | 1.41 | 0.39271015 | 6.72 | 2.16 |
| 91.98 | 21.74 | -1.06 | 0.75914413 | 100.91 | 11.88 |
| 112.61 | 32.91 | -1.05 | 0.81325406 | 100.74 | 39.13 |
| 3.02 | 1.01 | -1.05 | 0.80892092 | 5.33 | 1.72 |
| 69.09 | 33.08 | -1.25 | 0.48415563 | 33.25 | 23.72 |
| 71.4 | 30.94 | -1.53 | 0.07309064 | 40.81 | 16.6 |
| 2.99 | 1.01 | 41.86 | 0.00002237 | 81.24 | 136.22 |
| 3.81 | 4.28 | 3.5 | 0.00345347 | 9.45 | 3.58 |
| 5.77 | 4.7 | -1.21 | 0.58486897 | 6.12 | 2.98 |
| 54.42 | 25.57 | -1.64 | 0.14406739 | 30.45 | 8.92 |
| 18.25 | 2.84 | 1.26 | 0.22663999 | 25.26 | 7.9 |
| 4.69 | 6.19 | 1.91 | 0.27846673 | 8.93 | 9.46 |
| 11.19 | 9.66 | 1.25 | 0.59973818 | 11.16 | 7.78 |
| 55 | 15.8 | -1.14 | 0.32070816 | 46.65 | 27.93 |
| 21.77 | 12.42 | -1.01 | 0.97358805 | 13.78 | 11.55 |
| 40.55 | 32.36 | 2.39 | 0.08732109 | 64.4 | 62.93 |

| | | | | | |
|---|---|---|---|---|---|
| 62.03 | 18.46 | -1.34 | 0.04074011 | 53.86 | 22.23 |
| 142.94 | 37.84 | -1.08 | 0.71752566 | 104.53 | 23.66 |
| 362.4 | 127.32 | -1.12 | 0.62343663 | 212.32 | 110.69 |
| 5.48 | 6.25 | 1.04 | 0.91065675 | 5.71 | 1.92 |
| 2.99 | 1.01 | 1.92 | 0.10219369 | 4.85 | 2.19 |
| 7.56 | 4.88 | -1.07 | 0.80618238 | 6.69 | 1.19 |
| 21.62 | 8.87 | -1.05 | 0.81188422 | 15.17 | 9.19 |
| 57.52 | 24.91 | -1.27 | 0.44955593 | 34.03 | 27.89 |
| 3.6 | 2.5 | -1.17 | 0.56218463 | 4.11 | 2.64 |
| 94.16 | 27.49 | -1.42 | 0.04234549 | 69.92 | 12.9 |
| 4.49 | 4.94 | 1.05 | 0.8806898 | 4.6 | 2.3 |
| 13.62 | 7.72 | -1.23 | 0.65009332 | 14.15 | 2.03 |
| 290.65 | 66.95 | -1.17 | 0.34530547 | 212.24 | 61.12 |
| 3.29 | 1.47 | 1.08 | 0.75397336 | 4.11 | 2.64 |
| 60.59 | 11.94 | 1.07 | 0.63234955 | 53.18 | 8.62 |
| 18.32 | 13.37 | -1.41 | 0.35235748 | 10.38 | 8.95 |
| 94.23 | 39.22 | -1.25 | 0.38709676 | 68.53 | 44.83 |
| 42.29 | 20.98 | 1.15 | 0.53256887 | 33.34 | 26.36 |
| 14.43 | 9.64 | -1.19 | 0.58872873 | 9.31 | 3.11 |
| 41.76 | 14.56 | 1.49 | 0.03898151 | 57.86 | 58.46 |
| 157.55 | 23.72 | -1.09 | 0.38763002 | 128.61 | 45.38 |
| 591.65 | 103.81 | -1.12 | 0.39646545 | 506.24 | 101.1 |
| 20.66 | 8.64 | -2.21 | 0.06977661 | 13.6 | 4.03 |
| 11.14 | 8.17 | 1.69 | 0.15643571 | 14.02 | 6.39 |
| 33.67 | 15.17 | -1.84 | 0.06646696 | 15.13 | 3.63 |
| 9.89 | 5.57 | 1.16 | 0.59353358 | 9.79 | 3.92 |
| 2.99 | 1.01 | 1.52 | 0.15250428 | 5.18 | 2.42 |
| 133.5 | 58.37 | -1.05 | 0.85687703 | 87.33 | 64.75 |
| 2.99 | 1.01 | 1.29 | 0.28894269 | 4.11 | 2.64 |
| 27.11 | 8.5 | -1.04 | 0.82343936 | 32.14 | 15.04 |
| 4.66 | 3.76 | -1.42 | 0.23137853 | 4.85 | 2.19 |
| 40.01 | 18.73 | -2.59 | 0.04698095 | 12.15 | 10.96 |
| 89.04 | 39.56 | -1.37 | 0.25405845 | 47.36 | 27.86 |
| 45.9 | 22.67 | 1.17 | 0.59793359 | 44.94 | 24.36 |
| 60.56 | 12.96 | -1.03 | 0.79186803 | 58.92 | 26.92 |
| 3.69 | 2.98 | -1.29 | 0.3568483 | 4.85 | 2.19 |
| 47.68 | 19.34 | -1.29 | 0.21882027 | 48.07 | 10.25 |
| 40.37 | 17.96 | 1.1 | 0.72210413 | 37.94 | 13.7 |
| 188.6 | 53.06 | 1.11 | 0.4522745 | 166.15 | 56.74 |
| 73.42 | 25.11 | -1.67 | 0.02987641 | 36.89 | 6.32 |
| 490.49 | 192.09 | -1.59 | 0.02648109 | 290.48 | 122.44 |
| 6.22 | 4.29 | -1.38 | 0.36582264 | 5.82 | 1.76 |
| 13 | 7.53 | -1.01 | 0.96873349 | 9.67 | 3.26 |
| 3.51 | 3.45 | 1.22 | 0.56659091 | 5.57 | 3.73 |
| 25.44 | 12.93 | -1.58 | 0.08997749 | 13.45 | 9.56 |
| 9.18 | 7.45 | 1.5 | 0.20379491 | 10.66 | 3.93 |
| 3.53 | 4.16 | -1.23 | 0.4792617 | 4.52 | 2.33 |

| | | | | | |
|---|---|---|---|---|---|
| 11.36 | 9.22 | 1.28 | 0.48193771 | 11.86 | 3.34 |
| 6.12 | 8.53 | 3.18 | 0.00828591 | 13.79 | 4.63 |
| 14.06 | 9.51 | 2 | 0.06015198 | 21.82 | 12.51 |
| 17.82 | 11.75 | 4.16 | 0.00220419 | 56.68 | 34.43 |
| 4.09 | 4.62 | 2.25 | 0.02662189 | 9.66 | 5.23 |
| 3.57 | 3.98 | 1.92 | 0.02426722 | 6.65 | 3.2 |
| 4.11 | 2.35 | 1.06 | 0.8080163 | 6.57 | 3 |
| 22.75 | 7.98 | -1.14 | 0.63738048 | 14.71 | 4.99 |
| 5.11 | 4.71 | -1.12 | 0.71425724 | 5.64 | 3.91 |
| 25.49 | 10.49 | -1.27 | 0.22354597 | 21.29 | 4.91 |
| 3.63 | 1.7 | 1.02 | 0.9413045 | 4.85 | 2.19 |
| 243.64 | 92.58 | -1.29 | 0.18614893 | 160.58 | 150.21 |
| 8.84 | 10.22 | 1.88 | 0.24076948 | 12.52 | 10.04 |
| 29.17 | 15.52 | 3.02 | 0.00250425 | 51.22 | 75.69 |
| 8.75 | 6.39 | -1.27 | 0.39086899 | 10.27 | 2.84 |
| 13.68 | 10.35 | 2.48 | 0.01095911 | 29.33 | 26.9 |
| 5.42 | 5.6 | -1.51 | 0.29360527 | 5.09 | 4.18 |
| 2.99 | 1.01 | 1.31 | 0.30344674 | 5.33 | 1.72 |
| 138.29 | 22.76 | -1.37 | 0.00527491 | 92.32 | 20.94 |
| 176.94 | 46.24 | -1.19 | 0.28143412 | 136.68 | 30.02 |
| 55.28 | 16.82 | 1.25 | 0.28064325 | 66.34 | 22.71 |
| 18.33 | 11.19 | 2.38 | 0.01498016 | 24.43 | 14.64 |
| 32.6 | 18.14 | -2.58 | 0.00089588 | 19.02 | 3.96 |
| 38.96 | 11.13 | -1.04 | 0.81001103 | 40.19 | 3.67 |
| 13.23 | 4.88 | 12.29 | 0.00007031 | 100.24 | 94.07 |
| 28 | 9.64 | -1 | 0.98314041 | 35.54 | 9.01 |
| 49.7 | 19.19 | -1.45 | 0.04314789 | 29.53 | 20.59 |
| 1056.8 | 429.42 | -1.31 | 0.35619143 | 652.32 | 377.26 |
| 89.55 | 16.42 | -1.46 | 0.03990737 | 72.41 | 16.34 |
| 5.93 | 5.01 | 1.33 | 0.43777972 | 7.45 | 3.88 |
| 7.32 | 5.63 | 1.11 | 0.78386962 | 7.19 | 4.06 |
| 23.52 | 4.94 | -8.21 | 0.00001726 | 7.88 | 6.55 |
| 52.16 | 17.72 | -1.51 | 0.01834544 | 36.57 | 5.58 |
| 3.85 | 2.5 | 5.84 | 0.0000125 | 13.73 | 8.23 |
| 15.85 | 8.74 | -2.81 | 0.00746841 | 6.22 | 3 |
| 147.66 | 48.04 | -1.11 | 0.43976727 | 137.61 | 46.24 |
| 412.59 | 153.25 | -1.15 | 0.49423128 | 279.93 | 237.98 |
| 123.09 | 58.4 | 1.04 | 0.85808855 | 181.96 | 74.76 |
| 10.74 | 2.64 | -1.41 | 0.17285028 | 8.37 | 10.67 |
| 139.64 | 77.64 | -1.33 | 0.3474144 | 74.09 | 76.22 |
| 7.81 | 10.52 | 1.42 | 0.39722669 | 10.89 | 4.18 |
| 82.23 | 12.34 | -1.46 | 0.002045 | 73.65 | 13.96 |
| 39.46 | 13.99 | -1.24 | 0.30613777 | 51.8 | 7.34 |
| 166.89 | 49.34 | -1.25 | 0.18650968 | 119 | 14.71 |
| 21.41 | 6.67 | -1.36 | 0.26355168 | 14.5 | 13.28 |
| 3.6 | 2.5 | -1.09 | 0.76571542 | 5.18 | 2.42 |
| 3.02 | 1.01 | -1.05 | 0.80892092 | 4.52 | 2.33 |

| | | | | | |
|---|---|---|---|---|---|
| 4.74 | 4.02 | 1.7 | 0.23925231 | 7.7 | 3.46 |
| 115.05 | 37.03 | -1.28 | 0.20747818 | 78.94 | 42.63 |
| 117.89 | 38.25 | -1.11 | 0.65184706 | 83.13 | 30.1 |
| 3.62 | 2.55 | 1.56 | 0.2500132 | 7.47 | 5.09 |
| 31.6 | 4.77 | -1.08 | 0.58424139 | 26.47 | 4.4 |
| 172.91 | 39.4 | -1.31 | 0.117523 | 125.27 | 42.29 |
| 22.41 | 6.61 | 1.05 | 0.72851771 | 20.31 | 6.85 |
| 37.23 | 11.76 | -1.09 | 0.59538585 | 37.52 | 13.02 |
| 2.99 | 1.01 | 1.07 | 0.77423978 | 5.07 | 2.34 |
| 19.06 | 11.9 | -1.54 | 0.10179607 | 8.85 | 2.11 |
| 219.95 | 29.52 | 1.19 | 0.10748392 | 274.84 | 29.13 |
| 3.93 | 2.53 | -1.11 | 0.65981662 | 5.59 | 2.53 |
| 47.45 | 14.3 | -1.16 | 0.30430752 | 41.72 | 8.7 |
| 6.07 | 5.94 | -1.14 | 0.69901717 | 4.52 | 2.33 |
| 217.74 | 67.66 | -1.23 | 0.28731614 | 148.18 | 28.14 |
| 4.85 | 5.57 | 2.43 | 0.01884827 | 11.91 | 6.75 |
| 76.46 | 20.69 | -1.21 | 0.12596869 | 66.93 | 6.12 |
| 13.52 | 9.21 | -1.66 | 0.14938208 | 8.37 | 2.42 |
| 114.93 | 34.63 | -1.21 | 0.2170064 | 88.02 | 29.56 |
| 4.58 | 4.16 | -1.03 | 0.93282658 | 9.11 | 2.22 |
| 18.48 | 11.14 | -1.08 | 0.85361135 | 7.76 | 2.42 |
| 162.11 | 75.04 | -1.29 | 0.28567785 | 117.58 | 21.96 |
| 35.75 | 25.25 | 2.41 | 0.02032563 | 51.41 | 30.35 |
| 29.41 | 7.6 | -1.02 | 0.86719841 | 27.82 | 3.93 |
| 31.33 | 23.78 | -2.11 | 0.1083903 | 14.19 | 5.35 |
| 6.74 | 6.4 | 2.31 | 0.03843252 | 14.09 | 1.78 |
| 8.31 | 6.05 | 3.19 | 0.00479269 | 27.19 | 9.97 |
| 37.92 | 18.18 | 3.42 | 0.00508169 | 71.15 | 55.63 |
| 39.88 | 20.01 | 6.26 | 0.00000035 | 191.53 | 128.95 |
| 88.23 | 40.1 | -1.27 | 0.38380814 | 55.52 | 37.79 |
| 46.11 | 23.44 | -1.28 | 0.38088706 | 37.77 | 26.95 |
| 11.28 | 7.57 | -1.76 | 0.09403177 | 7.86 | 6.1 |
| 27.9 | 12.61 | -1.24 | 0.33905903 | 18.91 | 3.4 |
| 40.66 | 19.3 | -1.18 | 0.60099792 | 22.77 | 14.88 |
| 17.25 | 11.41 | 1.08 | 0.70534253 | 12.86 | 7.28 |
| 4.35 | 4.74 | 2.97 | 0.00456923 | 12.06 | 8 |
| 58.51 | 19.69 | -1.11 | 0.56282407 | 44.73 | 22.35 |
| 3.02 | 1.01 | 1.52 | 0.30027768 | 5.86 | 2.68 |
| 41.12 | 21.23 | -1.25 | 0.47946537 | 22.92 | 22.74 |
| 16.23 | 8.91 | -1.97 | 0.08712049 | 13.43 | 2.31 |
| 7.33 | 3.7 | 1.34 | 0.25710991 | 5.8 | 3.29 |
| 6.91 | 6.12 | 2.23 | 0.09913479 | 11.82 | 11.2 |
| 3.85 | 4.26 | 3.41 | 0.00443421 | 9.2 | 4.08 |
| 4.53 | 3.92 | 5.49 | 0.00008969 | 23.58 | 12.03 |
| 3.39 | 1.91 | 1.38 | 0.21019904 | 5.33 | 1.72 |
| 60.57 | 14.33 | -1.2 | 0.24129723 | 58.09 | 13.15 |
| 6.24 | 6.54 | 1.33 | 0.36990458 | 7.29 | 3.81 |

| | | | | | |
|---|---|---|---|---|---|
| 3.18 | 1.13 | 3.82 | 0.00000465 | 16.03 | 19.62 |
| 3.5 | 1.75 | 1.77 | 0.09858143 | 5.33 | 1.72 |
| 7 | 7.44 | 2.09 | 0.0753521 | 14.85 | 5.01 |
| 6.66 | 6.26 | 1.28 | 0.49998742 | 6.72 | 6.56 |
| 3.29 | 1.47 | 1.07 | 0.78907353 | 5.18 | 2.42 |
| 4.76 | 3.81 | 2.98 | 0.00150253 | 15.38 | 4.79 |
| 5.08 | 4.75 | 1.9 | 0.09060387 | 8.38 | 8.24 |
| 549.68 | 185.89 | -1.04 | 0.86447847 | 370.43 | 167.22 |
| 26.55 | 8.61 | -1.19 | 0.32356676 | 22.06 | 7.18 |
| 28.97 | 10.58 | 1.56 | 0.08321786 | 34.07 | 14.57 |
| 3.24 | 1.28 | 3.61 | 0.00000511 | 11.23 | 4.76 |
| 27.59 | 24.47 | 1.25 | 0.63092488 | 17.16 | 19.76 |
| 8.76 | 6.69 | 5.68 | 0.00015851 | 47.7 | 28.36 |
| 31.71 | 13.26 | 1.25 | 0.28103614 | 40.92 | 9.5 |
| 13.92 | 12.97 | 1.25 | 0.56388372 | 19.64 | 8.42 |
| 3.73 | 2.13 | 1.38 | 0.22973752 | 5.75 | 3.84 |
| 3.5 | 1.75 | 1.63 | 0.12660094 | 4.93 | 2.17 |
| 7.92 | 9.75 | 6.42 | 0.00122186 | 35.32 | 28.01 |
| 5.83 | 7.24 | -1.09 | 0.82057452 | 7.41 | 2.53 |
| 4.95 | 4.86 | -1.1 | 0.75329262 | 7.19 | 4.06 |
| 70.11 | 30.12 | -1.34 | 0.23884527 | 44.47 | 17.25 |
| 17.04 | 8.37 | 1.07 | 0.80427438 | 12.58 | 5.57 |
| 14.38 | 9.54 | -1.06 | 0.83088762 | 9.57 | 6.37 |
| 49.11 | 9.63 | 1.13 | 0.32642233 | 50.45 | 12.82 |
| 2.99 | 1.01 | 1.15 | 0.63546902 | 4.11 | 2.64 |
| 143.59 | 59.51 | -1.5 | 0.1436782 | 73.02 | 18.48 |
| 14.34 | 6.18 | -1.16 | 0.39376074 | 9.47 | 1.91 |
| 146.42 | 43 | -1.29 | 0.20350647 | 107.47 | 20.12 |
| 57.5 | 18.64 | -1.47 | 0.07570125 | 33.79 | 11.96 |
| 68.49 | 20.02 | 1.02 | 0.91902161 | 61.51 | 27.78 |
| 64.14 | 20.99 | -1.03 | 0.87403733 | 44.1 | 19.64 |
| 103.64 | 35.93 | -1.33 | 0.12361629 | 76.76 | 22.02 |
| 11.52 | 1.11 | -1.14 | 0.5405637 | 8.67 | 1.2 |
| 502.91 | 129.98 | -1.31 | 0.17405453 | 355 | 61.87 |
| 9.89 | 5.25 | -1.27 | 0.50090271 | 7.51 | 2.11 |
| 141.22 | 46.09 | -1.17 | 0.49517503 | 97.22 | 63.19 |
| 15.89 | 7.39 | -1.09 | 0.6554625 | 14.3 | 4.09 |
| 138.91 | 62.34 | -1.18 | 0.6276384 | 72.57 | 78.34 |
| 12.79 | 11.29 | -1.29 | 0.48428398 | 9.93 | 2.08 |
| 37.82 | 10.48 | 1.25 | 0.16031955 | 46.93 | 9.63 |
| 2.99 | 1.01 | -1.04 | 0.8469106 | 4.11 | 2.64 |
| 9.56 | 8.36 | 2.18 | 0.04047426 | 17.79 | 3.19 |
| 72.35 | 33.46 | -1.27 | 0.33288923 | 41.32 | 27.95 |
| 44.17 | 11.86 | -1.49 | 0.03316695 | 32.52 | 7.26 |
| 8.67 | 8.94 | -1.02 | 0.96824241 | 5.97 | 2.85 |
| 94.05 | 33.45 | -1.12 | 0.54219574 | 75.14 | 28.12 |
| 93.48 | 26.42 | -1.33 | 0.09211937 | 68.35 | 33.96 |

| | | | | | |
|---|---|---|---|---|---|
| 136.19 | 71.16 | -1.04 | 0.89532089 | 97.98 | 57.94 |
| 15.76 | 30.54 | -1.05 | 0.93840736 | 22.4 | 27.73 |
| 14.14 | 6.24 | 1.01 | 0.9577204 | 14 | 2.72 |
| 5.34 | 5.17 | -1.51 | 0.21236007 | 4.85 | 2.19 |
| 3.53 | 4.16 | -1.23 | 0.4792617 | 6.98 | 3.07 |
| 2.99 | 1.01 | -1.04 | 0.8469106 | 4.11 | 2.64 |
| 13.73 | 9.09 | 1.02 | 0.95032489 | 15.51 | 5.84 |
| 4.81 | 4.64 | 1.25 | 0.53574622 | 5.82 | 4.86 |
| 6.07 | 4.38 | -1.16 | 0.69649184 | 8.1 | 5.33 |
| 14.69 | 12.77 | -1.41 | 0.42546374 | 9.27 | 8.16 |
| 93.8 | 24.34 | -1.09 | 0.40673578 | 89.68 | 24.28 |
| 112.97 | 50.72 | -1.27 | 0.3282733 | 69.27 | 42.57 |
| 16.89 | 12.62 | -1.18 | 0.63999653 | 15.48 | 8.15 |
| 13.68 | 9.31 | -1.08 | 0.82927793 | 7.94 | 4.88 |
| 19.43 | 5.4 | -1.83 | 0.03677039 | 11.22 | 2.24 |
| 942.34 | 271.64 | -1.19 | 0.40009665 | 672.54 | 186.09 |
| 361.59 | 72.76 | -1.05 | 0.71959645 | 328.09 | 63.87 |
| 11.79 | 6.51 | 122.2 | 0.00000001 | 497.44 | 782.78 |
| 256.84 | 72.5 | -1.08 | 0.70705521 | 175.51 | 130.74 |
| 53.36 | 26.8 | -1.58 | 0.22312815 | 29.61 | 17.35 |
| 86.31 | 19.3 | 1.01 | 0.95208406 | 71.26 | 25.73 |
| 254.75 | 22.76 | -1.13 | 0.20775348 | 239.35 | 19.37 |
| 23.43 | 6.52 | -1.05 | 0.8131395 | 15.68 | 11.37 |
| 10.11 | 5.79 | 2.82 | 0.00619248 | 20.23 | 16.39 |
| 78.33 | 23.88 | -1.24 | 0.08535837 | 53.02 | 12.42 |
| 67.33 | 12.43 | 1.06 | 0.68496424 | 56.82 | 15.35 |
| 15.51 | 4.65 | -1.03 | 0.85402638 | 11.39 | 3.99 |
| 3.43 | 2 | 1.23 | 0.47529489 | 5.36 | 2.85 |
| 5.25 | 6.05 | 1.77 | 0.17538621 | 6.65 | 1.78 |
| 15.05 | 3.18 | -1.39 | 0.0304932 | 10.82 | 6.22 |
| 26.32 | 9.92 | -2.47 | 0.00139658 | 11.88 | 5.97 |
| 5.15 | 4.83 | 1.42 | 0.41379532 | 6.41 | 2.27 |
| 11.22 | 8.85 | -1.12 | 0.78334004 | 7.38 | 5.1 |
| 3.24 | 1.28 | 1.35 | 0.29904804 | 4.52 | 2.33 |
| 5.68 | 5.2 | 2 | 0.05430784 | 9.45 | 6.5 |
| 5.75 | 3.95 | -1.94 | 0.04128769 | 6.47 | 1.51 |
| 132.11 | 46.41 | 2.24 | 0.00147867 | 200.06 | 143.24 |
| 3.78 | 3.09 | -1.19 | 0.53084773 | 4.11 | 2.66 |
| 8.62 | 10.42 | 1.28 | 0.61711049 | 7.08 | 1.8 |
| 12.08 | 10.88 | 1.26 | 0.49507174 | 11.69 | 3.61 |
| 28.16 | 19.52 | 1.14 | 0.6923185 | 26.68 | 7.66 |
| 5.14 | 4.24 | 3.16 | 0.00297883 | 9.87 | 6 |
| 174.51 | 31.02 | -1.22 | 0.11850322 | 143.42 | 8.44 |
| 662.44 | 259.43 | -1.25 | 0.41526008 | 416.52 | 169.42 |
| 94.57 | 31.63 | -1.37 | 0.19331065 | 51.96 | 24.96 |
| 206.88 | 45.13 | -1.08 | 0.56257093 | 154.97 | 45.69 |
| 224.25 | 38.81 | -1.15 | 0.15757778 | 218.8 | 12.59 |

| | | | | | |
|---|---|---|---|---|---|
| 387.47 | 89.26 | -1.23 | 0.12618637 | 307.71 | 74.2 |
| 21.81 | 10.33 | -1.25 | 0.32742122 | 18.94 | 13.94 |
| 3.2 | 1.21 | 1.46 | 0.23773271 | 5.39 | 1.72 |
| 493.79 | 95.18 | -1.23 | 0.12702258 | 379.63 | 81.38 |
| 3.79 | 1.79 | -1.15 | 0.62561905 | 4.62 | 2.39 |
| 44.85 | 18.77 | -1.44 | 0.10385274 | 38.41 | 21.24 |
| 249.02 | 42.41 | -1.24 | 0.06683946 | 200.29 | 34 |
| 3.02 | 1.01 | 1.55 | 0.05604677 | 5.99 | 1.46 |
| 29 | 7.73 | -1.09 | 0.66131312 | 22.68 | 6.78 |
| 3.66 | 4.96 | 1.04 | 0.91140097 | 5.71 | 1.92 |
| 165.55 | 17.99 | 1.01 | 0.8493129 | 161.96 | 20.63 |
| 117.32 | 34.67 | -1.21 | 0.34601349 | 80.86 | 24.56 |
| 7.3 | 5.11 | 1.19 | 0.54087561 | 7.46 | 3.33 |
| 14.44 | 13.71 | -1.05 | 0.92867059 | 10.6 | 9.55 |
| 7.75 | 3.83 | -1.33 | 0.30488345 | 10.53 | 2.04 |
| 11.06 | 5.54 | 1.17 | 0.45470637 | 10.3 | 5.84 |
| 130.94 | 49.75 | -1.09 | 0.72999305 | 88.93 | 44.75 |
| 34.28 | 16.64 | -1.31 | 0.26000726 | 23.9 | 3.43 |
| 36.25 | 13.97 | 1.27 | 0.28595591 | 30.41 | 12.03 |
| 36.03 | 13.24 | -1.46 | 0.1381166 | 20 | 19.46 |
| 11.52 | 9.85 | -1.15 | 0.7342633 | 8.74 | 11.34 |
| 2.99 | 1.01 | 2.58 | 0.00888187 | 6.33 | 3.69 |
| 13.3 | 5.3 | -1.37 | 0.26686379 | 9.54 | 3.74 |
| 1794.94 | 274.95 | -1.18 | 0.20045832 | 1550.5 | 175.85 |
| 115.79 | 22.06 | -1.21 | 0.07613774 | 116.05 | 38.78 |
| 32.14 | 11.67 | 1.01 | 0.96777976 | 29.91 | 4.47 |
| 4.68 | 6.76 | 2.14 | 0.08029278 | 8.64 | 6.8 |
| 21.03 | 15.2 | -1.44 | 0.27800333 | 11.92 | 14.67 |
| 29.73 | 7.69 | -1.09 | 0.52922326 | 28.09 | 7.04 |
| 3.48 | 3.02 | 1.28 | 0.50380552 | 4.85 | 2.19 |
| 3.02 | 1.01 | -1.05 | 0.80892092 | 5.18 | 2.42 |
| 5.51 | 5.38 | 3.3 | 0.00451055 | 11.46 | 6.98 |
| 80.54 | 20.67 | -1.14 | 0.28121594 | 63.92 | 9.67 |
| 19.34 | 17.02 | 1.62 | 0.19709305 | 20.68 | 13.16 |
| 6.44 | 7.22 | 3.34 | 0.0127701 | 11.19 | 7.12 |
| 36.52 | 14.43 | 1.19 | 0.44328406 | 36.55 | 15.21 |
| 8.92 | 5.24 | 2.18 | 0.00926497 | 17.42 | 14.45 |
| 40.74 | 47.4 | -1.52 | 0.57899421 | 14.42 | 40.64 |
| 8.99 | 5.85 | 2.28 | 0.00796965 | 16.39 | 5.8 |
| 186.17 | 36.47 | 1.05 | 0.72381657 | 183.32 | 38.02 |
| 7.73 | 6.06 | 1.02 | 0.9687807 | 13.47 | 5.83 |
| 5.29 | 4.64 | 1.63 | 0.26026577 | 7.33 | 7.39 |
| 18.41 | 15.64 | 1.53 | 0.33081755 | 16.26 | 14.31 |
| 101.52 | 26.23 | -1.16 | 0.23257664 | 88.7 | 8.45 |
| 171.5 | 63.09 | -1.17 | 0.47893575 | 109.13 | 66.33 |
| 2.99 | 1.01 | 1.1 | 0.63673508 | 4.11 | 2.64 |
| 437.65 | 133.15 | -1.14 | 0.50135815 | 352.41 | 51.74 |

| | | | | | |
|---|---|---|---|---|---|
| 17.42 | 13.93 | -1.1 | 0.79939264 | 12.85 | 12.46 |
| 59.31 | 21.21 | -1.03 | 0.87779695 | 42.73 | 20.19 |
| 9.64 | 8.75 | 1.99 | 0.18620783 | 15.81 | 15.01 |
| 14.72 | 10.78 | 2.67 | 0.00627545 | 29.25 | 12.26 |
| 5.15 | 3.67 | 1.64 | 0.1760952 | 5.73 | 2.75 |
| 9.44 | 11.7 | 1.42 | 0.4783558 | 8.23 | 5.48 |
| 3.34 | 1.86 | 1.19 | 0.53327096 | 6.31 | 4.02 |
| 4 | 5.57 | 1.14 | 0.71746141 | 5.45 | 2.37 |
| 5.28 | 4.18 | -1.08 | 0.86038625 | 4.11 | 2.64 |
| 123.67 | 25.97 | -1.17 | 0.21124832 | 117.23 | 19.09 |
| 281.6 | 79.25 | -1.14 | 0.32336041 | 210.85 | 75.6 |
| 4.62 | 2.78 | 2.05 | 0.00693864 | 10.39 | 5.55 |
| 18.12 | 18.14 | -1.11 | 0.83246487 | 11.29 | 7.86 |
| 12.43 | 7.86 | -1.23 | 0.48620135 | 9.13 | 4.64 |
| 9.01 | 6.7 | 1.07 | 0.85302436 | 11.44 | 4.04 |
| 18.58 | 8.68 | -1.66 | 0.10817674 | 19.58 | 5.95 |
| 118.94 | 27.04 | -1.22 | 0.27584165 | 95.44 | 44.37 |
| 30.3 | 16.33 | -1.42 | 0.19591017 | 21.32 | 6.51 |
| 17.04 | 36.91 | -1.15 | 0.74637908 | 13.29 | 8.1 |
| 3.77 | 2.5 | 1.14 | 0.63079262 | 5.71 | 1.92 |
| 25.85 | 17.38 | -1.1 | 0.71441609 | 19.41 | 13.88 |
| 10.56 | 3.79 | -1.17 | 0.46173611 | 9.01 | 4.69 |
| 105.5 | 49.9 | -1.5 | 0.14791334 | 55.97 | 56.34 |
| 3.5 | 1.75 | 2.68 | 0.01049994 | 8.56 | 3.6 |
| 16.79 | 10.37 | -1.22 | 0.54824048 | 11.41 | 5.48 |
| 36.26 | 14.32 | -1.6 | 0.02287356 | 20.91 | 10.71 |
| 163.75 | 38.36 | -2.74 | 0.00049147 | 78.15 | 24.58 |
| 126.57 | 45.87 | -1.3 | 0.14686128 | 83.84 | 33.42 |
| 5 | 5.2 | 1.05 | 0.89807224 | 5.57 | 3.73 |
| 18.6 | 16.38 | 5.67 | 0.00033613 | 66.23 | 46.92 |
| 13.09 | 7.53 | -1.46 | 0.20209062 | 15.93 | 4.83 |
| 58.25 | 27.44 | -3.62 | 0.0023256 | 12.17 | 11.78 |
| 508 | 106.02 | -1.14 | 0.29615664 | 366.42 | 110.85 |
| 49.4 | 15.67 | -1.72 | 0.00114748 | 39.48 | 12.96 |
| 664.46 | 179.09 | -1.2 | 0.30232772 | 488.72 | 91.81 |
| 73.18 | 19.09 | -1.3 | 0.12748364 | 48.46 | 14.53 |
| 46.95 | 20.74 | -1.3 | 0.25282067 | 28.6 | 19.16 |
| 114.39 | 20.18 | -1.06 | 0.60336852 | 107.23 | 4.5 |
| 41.38 | 10.8 | 1.6 | 0.00131697 | 57.41 | 14.72 |
| 36.64 | 11.84 | -1.28 | 0.29547945 | 22.99 | 4.25 |
| 85.14 | 18.17 | -1.25 | 0.03364366 | 68.84 | 15.84 |
| 56.71 | 22.39 | -1.03 | 0.87904173 | 70.77 | 12.79 |
| 98.32 | 26.22 | -1.2 | 0.33504909 | 76.76 | 18.4 |
| 26.95 | 12.97 | -1.32 | 0.35966611 | 18.71 | 13.03 |
| 3.18 | 1.13 | 1.34 | 0.28409168 | 4.94 | 2.14 |
| 282.58 | 126.48 | -1.62 | 0.15577585 | 145.55 | 98.55 |
| 140.66 | 39.21 | -1.24 | 0.33354342 | 95.99 | 56.06 |

| | | | | | |
|---|---|---|---|---|---|
| 2.99 | 1.01 | 1.4 | 0.22389297 | 4.54 | 2.38 |
| 6.69 | 6.2 | 1.81 | 0.22176296 | 7.24 | 3.11 |
| 3.48 | 3.02 | 1.22 | 0.49465925 | 4.11 | 2.64 |
| 857.33 | 114.32 | -1.16 | 0.0767829 | 718.93 | 61.18 |
| 15.89 | 6.04 | -1.26 | 0.39177123 | 11.73 | 3.53 |
| 9.94 | 11.9 | -1.16 | 0.71757072 | 6.2 | 5.99 |
| 15.19 | 3.94 | -2.16 | 0.05122804 | 10.52 | 2.39 |
| 39.87 | 27.55 | 1.03 | 0.94987291 | 27.63 | 27.17 |
| 14.75 | 13.12 | -1.22 | 0.55085123 | 9.79 | 4.22 |
| 2115.13 | 467.42 | -1.26 | 0.07081376 | 1781.09 | 219.62 |
| 15.24 | 10.11 | -1.33 | 0.54651868 | 18.45 | 5.09 |
| 37.82 | 17.29 | -2.14 | 0.00809088 | 24.26 | 7.93 |
| 2999.02 | 401.05 | -1.17 | 0.09908363 | 2883.24 | 550.69 |
| 114.24 | 24.94 | -1.41 | 0.03453765 | 87.86 | 15.95 |
| 54.26 | 23.75 | -1.75 | 0.04225403 | 27.79 | 11.62 |
| 105.87 | 56.53 | -1.18 | 0.62486881 | 62.36 | 26.95 |
| 51.8 | 14.6 | -1.13 | 0.45697197 | 45.13 | 11.26 |
| 10.12 | 4.38 | -1.29 | 0.52403271 | 10.93 | 2.66 |
| 6.41 | 6.81 | 1.45 | 0.36104643 | 7.11 | 2.16 |
| 42.34 | 9.75 | -1.15 | 0.40595737 | 41.43 | 13.96 |
| 268.96 | 40.26 | -1.08 | 0.47468677 | 263.08 | 55.7 |
| 243.59 | 42.95 | -1.25 | 0.04160903 | 189.94 | 65.71 |
| 288.15 | 98.79 | -1.06 | 0.83235228 | 208.7 | 81.72 |
| 6.37 | 4.9 | 1.62 | 0.14826196 | 7.71 | 4.89 |
| 30.89 | 7.89 | -1.39 | 0.06246158 | 25.77 | 6.22 |
| 2.99 | 1.01 | 1.21 | 0.41063192 | 4.11 | 2.64 |
| 676.26 | 100.99 | -1.13 | 0.19540262 | 533.2 | 65.81 |
| 386.19 | 44.13 | -1.18 | 0.04409121 | 312.13 | 93.15 |
| 952.61 | 177.81 | -1.2 | 0.10620692 | 731.56 | 156.77 |
| 208.55 | 58.09 | -1.27 | 0.25886253 | 140.09 | 59.68 |
| 172.03 | 53.88 | -1.49 | 0.06020997 | 106.2 | 57.74 |
| 94.56 | 23.58 | -1.26 | 0.17639162 | 100.09 | 26.14 |
| 32.28 | 12.57 | -1.47 | 0.14261335 | 17.36 | 2.96 |
| 26.01 | 10.45 | -1.52 | 0.12445977 | 20.52 | 5.63 |
| 25.98 | 11.5 | 1.34 | 0.28268957 | 25.13 | 8.4 |
| 40.6 | 18.32 | -1.07 | 0.85345727 | 23.27 | 13.86 |
| 17.83 | 10.09 | 3.02 | 0.01207072 | 37.23 | 19.94 |
| 133.72 | 31.89 | -1.14 | 0.36295372 | 146.08 | 36.12 |
| 7.42 | 6.82 | -1.45 | 0.35362676 | 7.83 | 8.32 |
| 3.51 | 2.17 | 1.5 | 0.12794714 | 5.27 | 2.68 |
| 3.24 | 1.28 | 11.36 | 0.00000001 | 26.85 | 20.26 |
| 260.24 | 46.74 | -1.16 | 0.09171396 | 221.29 | 56.7 |
| 7.93 | 7.28 | -1.15 | 0.73928362 | 7.74 | 3.09 |
| 4.76 | 3.43 | 1.66 | 0.17661941 | 7.38 | 5.1 |
| 154.98 | 17.3 | -1.09 | 0.41600609 | 144.08 | 25.09 |
| 5.33 | 4.4 | 5.1 | 0.00347193 | 17.63 | 17.61 |
| 2.99 | 1.01 | 1.1 | 0.63673508 | 4.11 | 2.64 |

| | | | | | |
|---|---|---|---|---|---|
| 577.55 | 119.54 | -1.12 | 0.30342153 | 453.72 | 65.74 |
| 757.47 | 175.01 | -1.21 | 0.2270761 | 551.41 | 153.66 |
| 409.02 | 46.75 | -1.15 | 0.12890458 | 372.04 | 18.73 |
| 4.76 | 4.15 | -1.15 | 0.66168928 | 5.22 | 3.2 |
| 34.45 | 11.38 | -1.22 | 0.34363544 | 27.14 | 9.43 |
| 20.78 | 9.05 | -1.01 | 0.97841835 | 17.44 | 10.15 |
| 4.2 | 3.09 | 1.25 | 0.5002079 | 6.96 | 2.7 |
| 7.93 | 4.98 | -1.34 | 0.30027467 | 7.2 | 2.74 |
| 9.85 | 9.92 | -1.5 | 0.40283671 | 8.96 | 13.66 |
| 9.85 | 10.43 | 1.18 | 0.7233696 | 6.75 | 1.64 |
| 17.24 | 9.99 | -1.19 | 0.56466359 | 14.4 | 3.9 |
| 2.99 | 1.01 | 1.1 | 0.63673508 | 5.07 | 2.34 |
| 104.95 | 20.97 | -1.34 | 0.04129653 | 86.02 | 13.03 |
| 46.14 | 8.52 | -1.12 | 0.27042586 | 53.52 | 13.25 |
| 87.95 | 22.64 | 1.02 | 0.8429727 | 88.2 | 15.75 |
| 176.52 | 41.93 | -1.22 | 0.22206657 | 139.15 | 29 |
| 336.69 | 123.67 | -1.17 | 0.51855528 | 209.93 | 78.37 |
| 37.06 | 18.94 | 1.1 | 0.77262354 | 29.74 | 20.92 |
| 12.41 | 7.75 | 1.72 | 0.05869976 | 19.39 | 8.81 |
| 18.92 | 7.43 | -1.3 | 0.12106586 | 19.09 | 10.85 |
| 36.9 | 19.49 | 1.08 | 0.72368711 | 27.13 | 9.99 |
| 206.42 | 25.01 | -1.08 | 0.33369306 | 186.74 | 20 |
| 26.77 | 19.86 | 1.16 | 0.76050156 | 18.83 | 23.77 |
| 6.75 | 5.83 | 1.93 | 0.0476535 | 9.38 | 6.89 |
| 46.86 | 14.12 | -1.58 | 0.01299946 | 37.55 | 6.71 |
| 152.15 | 48.46 | -1.22 | 0.27506214 | 115.37 | 22.83 |
| 627.42 | 136.43 | -1.24 | 0.12830782 | 461.66 | 77.49 |
| 414.68 | 113.34 | -1.24 | 0.14224164 | 316 | 56.86 |
| 95.52 | 40.3 | -1.03 | 0.85515958 | 113.44 | 33.39 |
| 983.71 | 287.23 | -1.19 | 0.35310981 | 687.15 | 252.65 |
| 74.37 | 33.31 | -1.01 | 0.95346266 | 84.64 | 31.07 |
| 680.87 | 163.41 | -1.11 | 0.51635343 | 531.25 | 206.08 |
| 2084.03 | 742.29 | -1.25 | 0.32940596 | 1321.88 | 680.56 |
| 55.36 | 23.87 | -1.29 | 0.36202204 | 34.64 | 21.19 |
| 749.95 | 186.81 | -1.18 | 0.23350485 | 589.82 | 121.77 |
| 11.6 | 7.93 | -1.25 | 0.5753606 | 7.69 | 4.3 |
| 20.2 | 15.05 | 6.27 | 0.00343106 | 116.22 | 84.24 |
| 9.76 | 14.51 | -1.14 | 0.73868555 | 8.99 | 12.53 |
| 120.98 | 40.71 | 1.02 | 0.92525202 | 113.84 | 25.73 |
| 441.47 | 119.5 | -1.15 | 0.37920597 | 415.69 | 81.9 |
| 9.08 | 7.81 | 1.58 | 0.22568491 | 13.48 | 5.08 |
| 4.98 | 4.14 | -1.13 | 0.73853135 | 5.77 | 2 |
| 32.8 | 14.88 | -1.3 | 0.32637572 | 18.29 | 9.72 |
| 69.5 | 34.99 | -1.81 | 0.00754758 | 37.24 | 16.25 |
| 3.18 | 1.13 | 2.21 | 0.00838773 | 6.51 | 1.78 |
| 20.51 | 11.47 | 4.48 | 0.00004529 | 46.23 | 39.52 |
| 3.62 | 2.55 | 1.46 | 0.26586059 | 4.15 | 2.66 |

| | | | | | |
|---|---|---|---|---|---|
| 78.59 | 26.68 | -1.42 | 0.04553426 | 44.37 | 6.68 |
| 17.89 | 8.36 | 1.12 | 0.64030921 | 17.64 | 5.68 |
| 87.11 | 30.72 | -1.02 | 0.94474626 | 71.07 | 42.82 |
| 18.76 | 7.79 | -1.54 | 0.06933649 | 14.53 | 4.99 |
| 35.56 | 15.98 | 1.22 | 0.32754305 | 37.05 | 8.21 |
| 53.27 | 15.18 | -1.11 | 0.4959361 | 40.7 | 17.18 |
| 117.6 | 29.74 | -1.14 | 0.20682417 | 113.41 | 25.23 |
| 64.75 | 23.33 | -1.09 | 0.52496088 | 60.84 | 14.63 |
| 28.6 | 9.9 | -1.41 | 0.04197471 | 18.91 | 4.17 |
| 5.42 | 6.06 | 1.05 | 0.913939 | 5.24 | 2.5 |
| 2.99 | 1.01 | -1.04 | 0.8469106 | 4.11 | 2.64 |
| 25.51 | 7.87 | -1.02 | 0.91682243 | 30.24 | 7.62 |
| 3.65 | 2.12 | 1.43 | 0.17567706 | 6.21 | 1.83 |
| 6.76 | 8.25 | 1.23 | 0.64409339 | 6.57 | 5.51 |
| 77.51 | 26.48 | -1.17 | 0.40144309 | 59.99 | 26.27 |
| 53.93 | 29.99 | -1.21 | 0.5827961 | 36.48 | 25.67 |
| 3.51 | 2.17 | 1.45 | 0.27258584 | 4.52 | 2.33 |
| 4.95 | 4.57 | -1.17 | 0.59551162 | 4.85 | 2.19 |
| 6.57 | 9.98 | 1.49 | 0.35621864 | 8.68 | 4.55 |
| 425.11 | 85.1 | -1.2 | 0.16301256 | 328.71 | 68.54 |
| 104.44 | 84.05 | 1.14 | 0.72551352 | 52.63 | 36.39 |
| 423.81 | 75.63 | -1.2 | 0.09762558 | 355.2 | 94.25 |
| 72.54 | 21.21 | -1.31 | 0.13306923 | 51.92 | 11.31 |
| 8.73 | 5.08 | -1.46 | 0.22818416 | 5.98 | 2.76 |
| 246.62 | 55.53 | -1.1 | 0.41017815 | 231.94 | 34.06 |
| 317.39 | 88.3 | 1.14 | 0.28715867 | 336.91 | 100.1 |
| 12.93 | 3.74 | -1.83 | 0.09300946 | 10.98 | 5.17 |
| 6.32 | 4.37 | 3.44 | 0.01271749 | 17.67 | 12.94 |
| 127.34 | 10.38 | -1.19 | 0.08250352 | 122.95 | 13.09 |
| 274.5 | 29.14 | -1.14 | 0.13874836 | 238.05 | 20.88 |
| 18.25 | 9.87 | 1.03 | 0.89961243 | 14.15 | 6.01 |
| 34.24 | 17.22 | 1.05 | 0.8176955 | 44.93 | 18.99 |
| 57.1 | 12.98 | -1.24 | 0.20893791 | 54.86 | 17.73 |
| 2.99 | 1.01 | 2.63 | 0.00013027 | 8.26 | 4.96 |
| 445.53 | 65.62 | -1.07 | 0.60192996 | 389.22 | 96.78 |
| 36.18 | 6.87 | -1.76 | 0.00003957 | 19.77 | 10.27 |
| 21.88 | 6.24 | -1.12 | 0.71120214 | 12.34 | 12.18 |
| 81.44 | 43.81 | 1.35 | 0.36514753 | 72.12 | 64.04 |
| 2.99 | 1.01 | 1.52 | 0.12979881 | 4.11 | 2.64 |
| 3.29 | 1.47 | -1.08 | 0.75099444 | 4.11 | 2.64 |
| 4.28 | 4.25 | 1.17 | 0.60891396 | 5.71 | 1.42 |
| 311.2 | 147.95 | 1.13 | 0.6416657 | 253.47 | 130.24 |
| 667.11 | 99.21 | -1.13 | 0.31589073 | 530.32 | 118.4 |
| 45.19 | 30.19 | 3.19 | 0.0095824 | 78.13 | 61.08 |
| 153.36 | 18.75 | -1.06 | 0.29540497 | 162.46 | 27.96 |
| 44.94 | 21.64 | -1.36 | 0.32502204 | 29.62 | 7.28 |
| 30.4 | 19.72 | -1.35 | 0.23434593 | 21.51 | 13.39 |

| | | | | | |
|---|---|---|---|---|---|
| 6.64 | 9.91 | 3.78 | 0.00383011 | 32.79 | 11.02 |
| 3.02 | 1.01 | 9.84 | 0.0002083 | 22.34 | 16.87 |
| 1659.57 | 483.52 | -1.39 | 0.06808725 | 1070.73 | 459.01 |
| 24.8 | 10.37 | -1.04 | 0.86983824 | 21.12 | 9.1 |
| 520.11 | 165.49 | -1.25 | 0.28268182 | 350.28 | 196.82 |
| 34.33 | 11.46 | -1.06 | 0.67023587 | 28.03 | 10.51 |
| 20.73 | 11.98 | -1.07 | 0.83413917 | 12.4 | 8.12 |
| 181.55 | 68.63 | -1.15 | 0.59582424 | 122 | 90.07 |
| 10.01 | 5.98 | -1.1 | 0.74568373 | 8.59 | 4.93 |
| 383.73 | 74.97 | -1.17 | 0.21873283 | 292.02 | 115.25 |
| 440.28 | 116.36 | -1.14 | 0.24367735 | 420.14 | 44.41 |
| 29.17 | 17.78 | -1.44 | 0.17015007 | 16.25 | 4.37 |
| 21.73 | 7.22 | -1.05 | 0.79304177 | 20.35 | 3.05 |
| 6.35 | 9.79 | -1.16 | 0.73319983 | 4.84 | 2.21 |
| 6.8 | 8.95 | 1.12 | 0.78478974 | 7.38 | 5.1 |
| 45.3 | 28.54 | -1.05 | 0.90597719 | 30.68 | 22.88 |
| 3.51 | 2.17 | -1.19 | 0.48286533 | 4.11 | 2.64 |
| 54.7 | 13.47 | -1.65 | 0.00731351 | 43 | 12.09 |
| 364.36 | 119.22 | -1.22 | 0.35420954 | 245.86 | 90.67 |
| 28.96 | 10.5 | -1.31 | 0.09257387 | 32.44 | 6.12 |
| 4.23 | 5.48 | 2.03 | 0.03807702 | 8.13 | 2.39 |
| 81.28 | 30.82 | -1.45 | 0.17362785 | 45.69 | 18.28 |
| 2.99 | 1.01 | -1.04 | 0.8469106 | 4.15 | 2.66 |
| 18.3 | 7.87 | 2.73 | 0.00024938 | 48.57 | 26.46 |
| 12.87 | 7.48 | -2.53 | 0.02113602 | 6.75 | 5 |
| 14.18 | 4.22 | 1.74 | 0.07675276 | 17.04 | 5.74 |
| 2.99 | 1.01 | 6.77 | 0.00035484 | 14.68 | 13.7 |
| 40.48 | 24.98 | 1.75 | 0.04718731 | 52.79 | 34.83 |
| 7.4 | 8.24 | 2.6 | 0.05402044 | 14.73 | 15.47 |
| 4.61 | 3.76 | 1.32 | 0.423448 | 6.18 | 1.63 |
| 3.98 | 5.14 | 1.73 | 0.1785055 | 4.93 | 2.44 |
| 55.19 | 21.49 | 1.8 | 0.09433497 | 75.76 | 49.47 |
| 43.09 | 12.61 | -1.51 | 0.012453 | 28.23 | 6.15 |
| 136.35 | 61.89 | -1.41 | 0.23458083 | 75.81 | 34.54 |
| 22.24 | 7.92 | -1.27 | 0.20654379 | 20.97 | 7.89 |
| 59.4 | 30.93 | -1.42 | 0.29604375 | 29.84 | 29.37 |
| 20.6 | 7.61 | -1.41 | 0.26277575 | 22.65 | 6.38 |
| 4.47 | 8.23 | -1.04 | 0.92443573 | 6.98 | 2.29 |
| 21.51 | 14.16 | -1.4 | 0.39862585 | 11.38 | 11.58 |
| 64.53 | 25.78 | -1.28 | 0.31976089 | 36.35 | 22.16 |
| 3.18 | 1.13 | 1.11 | 0.66631901 | 5.07 | 2.34 |
| 185.99 | 91.46 | -1.46 | 0.25697005 | 91.41 | 63.82 |
| 95.94 | 27.14 | -1.32 | 0.2807824 | 69.72 | 9.42 |
| 3.1 | 1.04 | 1.23 | 0.50737065 | 4.12 | 2.69 |
| 7.82 | 4.99 | 1.2 | 0.56670201 | 8.27 | 8.12 |
| 11.78 | 7.86 | -1.22 | 0.6122486 | 5.18 | 2.42 |
| 57.58 | 26.73 | -1.2 | 0.43427876 | 43.51 | 14.94 |

| | | | | | |
|---|---|---|---|---|---|
| 4.62 | 7.51 | 1.06 | 0.88236284 | 6.21 | 3.5 |
| 8.11 | 5.91 | -1.4 | 0.39305973 | 6.85 | 2.26 |
| 12.22 | 7.62 | 3.45 | 0.000648 | 35.14 | 19.06 |
| 5.16 | 6.29 | 1.86 | 0.10840269 | 8.87 | 4.67 |
| 4.3 | 5.06 | 1.31 | 0.52484107 | 5.18 | 2.42 |
| 3.02 | 1.01 | 1.09 | 0.73431605 | 4.11 | 2.64 |
| 8.2 | 9.99 | 1.42 | 0.43671072 | 7.87 | 4.18 |
| 22.56 | 5.74 | -1.61 | 0.14451845 | 11.61 | 1.97 |
| 9.39 | 6.64 | 1.11 | 0.78520191 | 7.71 | 4.19 |
| 4.02 | 3.26 | 1.13 | 0.70264083 | 5.33 | 1.72 |
| 2.99 | 1.01 | 1.6 | 0.03109859 | 4.84 | 2.21 |
| 14.92 | 14.52 | -1.01 | 0.97935432 | 8.79 | 8.45 |
| 28.63 | 16.39 | -1.06 | 0.8565824 | 21.46 | 11.02 |
| 261.52 | 59.78 | -1.12 | 0.46595779 | 216.89 | 85.72 |
| 12.38 | 9.75 | -2.7 | 0.0112379 | 7.66 | 5.07 |
| 13.4 | 9.06 | 1.19 | 0.68962663 | 9.33 | 7.12 |
| 4.91 | 6.4 | 1.44 | 0.39276877 | 6.64 | 4.09 |
| 38.13 | 16.04 | -1.2 | 0.53054661 | 27.28 | 22.69 |
| 71.38 | 24.48 | -1.14 | 0.51328683 | 54.95 | 26.63 |
| 32.34 | 20.14 | -1.02 | 0.96566457 | 20.51 | 11.36 |
| 4.33 | 4.14 | 1.33 | 0.42762864 | 5.07 | 2.34 |
| 5.34 | 5.3 | 1.21 | 0.6637814 | 6.41 | 1.5 |
| 21.33 | 10.18 | -2.85 | 0.00473727 | 9.43 | 2.59 |
| 4.55 | 4.32 | -1.28 | 0.4093717 | 4.11 | 2.64 |
| 7.19 | 6.36 | 1.56 | 0.19177455 | 8.4 | 2.67 |
| 18.78 | 11.4 | -1.18 | 0.61574113 | 13.58 | 5.27 |
| 47.28 | 12.78 | -1.17 | 0.38790074 | 37.41 | 5 |
| 5.5 | 5.45 | 1.28 | 0.53822798 | 5.71 | 1.42 |
| 3.55 | 2.15 | -1.1 | 0.72384751 | 5.33 | 1.72 |
| 13.47 | 5.56 | -1.5 | 0.09463812 | 9.39 | 7.85 |
| 6.09 | 6.68 | -1.32 | 0.46387604 | 6.72 | 2.16 |
| 23.73 | 128.55 | -2.97 | 0.09331571 | 8.36 | 5.3 |
| 803.03 | 146.3 | -1.03 | 0.78464472 | 664.12 | 90.09 |
| 179.04 | 66.67 | -1.01 | 0.96190172 | 214.75 | 69.55 |
| 307.92 | 111.74 | -1.06 | 0.82842779 | 217.25 | 101.21 |
| 16 | 12.57 | 1.32 | 0.48898664 | 33.76 | 6.91 |
| 488.56 | 208.07 | -1.82 | 0.10917356 | 235.97 | 47.98 |
| 55.73 | 15.33 | 1.1 | 0.39792618 | 57.64 | 12.17 |
| 9.72 | 5.5 | 1.17 | 0.55345601 | 9.39 | 2.79 |
| 17.22 | 17.01 | -1.12 | 0.83008271 | 18.6 | 15.02 |
| 236.56 | 61.39 | -1.27 | 0.15616469 | 177.28 | 46.46 |
| 7.23 | 5.37 | 1.65 | 0.13856526 | 10.22 | 2.59 |
| 69.59 | 15.26 | -1.38 | 0.02663056 | 44.02 | 12.11 |
| 49.16 | 16.7 | -1.18 | 0.30824476 | 40.89 | 11.16 |
| 109.54 | 13.36 | -1.21 | 0.06243617 | 110 | 6.79 |
| 57.29 | 20.61 | -1.14 | 0.59524882 | 46.45 | 9.15 |
| 3.97 | 3.63 | -1.02 | 0.9419874 | 4.52 | 2.33 |

| | | | | | |
|---|---|---|---|---|---|
| 26.19 | 14.74 | -1.03 | 0.88666505 | 22.24 | 13.44 |
| 6.09 | 4.39 | 1.36 | 0.41003028 | 7.24 | 3.11 |
| 207.44 | 20.51 | -1.15 | 0.04797677 | 195.64 | 24.05 |
| 21.22 | 2.45 | 1.02 | 0.84616619 | 22.85 | 4.88 |
| 3.47 | 3.15 | 6.22 | 0.00022011 | 21.25 | 16.19 |
| 65.19 | 19.57 | -1.34 | 0.02056275 | 44.68 | 8.35 |
| 66.07 | 25.96 | 1.07 | 0.74107707 | 89.07 | 29.99 |
| 2.99 | 1.01 | -1.04 | 0.8469106 | 4.11 | 2.64 |
| 116.75 | 17.75 | -1.06 | 0.36511207 | 110.75 | 25.09 |
| 54.48 | 10.37 | 1.03 | 0.78072006 | 60.86 | 13.07 |
| 83.55 | 27.73 | -1.2 | 0.36896676 | 51.86 | 11.4 |
| 50.42 | 17.09 | -1 | 0.99552494 | 40.65 | 23.02 |
| 13.39 | 3.79 | -1.13 | 0.56892323 | 11.09 | 2.93 |
| 48.13 | 11.56 | -1.01 | 0.90642828 | 55.59 | 6.91 |
| 21.32 | 8.64 | -1.33 | 0.26199034 | 22.78 | 7.89 |
| 37.94 | 10.64 | -1.27 | 0.07199083 | 28.02 | 10.43 |
| 107.43 | 38.59 | -1.04 | 0.848153 | 84.29 | 48.51 |
| 35.17 | 14.43 | -1.21 | 0.31147528 | 30.43 | 15 |
| 76.13 | 9.42 | 1 | 0.97500074 | 67.92 | 11.45 |

| WT-CTL | StDev of W | gKO-LPS vs. WT-CTL | P value of: gKO-LPS vs. WT-CTL |
|---|---|---|---|
| 26.37 | 8.68 | -2.01 | 0.00750167 |
| 4.39 | 3.79 | -1.07 | 0.84374195 |
| 5.83 | 5.95 | 1.38 | 0.28139243 |
| 40.61 | 11.58 | -1.22 | 0.13351049 |
| 14.27 | 5.96 | -1.5 | 0.0370039 |
| 5.45 | 5.26 | 1.54 | 0.19383405 |
| 122.62 | 32.5 | -1.28 | 0.10077886 |
| 454.41 | 214.9 | -1.82 | 0.08440495 |
| 134.14 | 28.97 | -1.3 | 0.06721751 |
| 43.23 | 21.18 | -1.74 | 0.06773581 |
| 89.93 | 16.33 | -1.2 | 0.26826641 |
| 210.54 | 76.59 | -1.52 | 0.12819353 |
| 57.09 | 13.73 | -1.36 | 0.29804337 |
| 40.58 | 7.9 | -1.06 | 0.62436253 |
| 9.53 | 11.53 | 1.09 | 0.78590101 |
| 510.39 | 137.13 | -1.46 | 0.03153893 |
| 10.82 | 12.43 | 2.07 | 0.07491255 |
| 1008.81 | 156.02 | -1.43 | 0.01121129 |
| 32.48 | 18.3 | -2.33 | 0.01265948 |
| 45.16 | 15.55 | -1.59 | 0.01227885 |
| 95.49 | 35.5 | -1.53 | 0.10528569 |
| 3.5 | 1.75 | 1.17 | 0.5958128 |
| 112.85 | 27.39 | -1.28 | 0.1534756 |
| 225.36 | 35.65 | 1.01 | 0.88280851 |
| 3.02 | 1.01 | 2.35 | 0.05185854 |
| 57.74 | 27.25 | -1.64 | 0.06079888 |
| 273.61 | 67.54 | -1.32 | 0.03741048 |
| 120.37 | 40.04 | -1.25 | 0.16393402 |
| 26.82 | 16.31 | -1.24 | 0.19544227 |
| 236.13 | 84.29 | -1.65 | 0.06121946 |
| 49.27 | 17.13 | -1.41 | 0.19770725 |
| 84.1 | 42.72 | -1.51 | 0.25648957 |
| 442.01 | 76.28 | -1.03 | 0.73917305 |
| 45.15 | 18.22 | -1.61 | 0.10536142 |
| 70.51 | 34.19 | -1.46 | 0.21315074 |
| 4.26 | 3.19 | 1.8 | 0.03108872 |
| 66.78 | 20.17 | -1.35 | 0.10282302 |
| 20.72 | 6.83 | -1.15 | 0.57141513 |
| 126.03 | 37.78 | -1.39 | 0.07120807 |
| 54.64 | 28.33 | -1.43 | 0.21318004 |
| 8.16 | 7.84 | 1.05 | 0.88817084 |
| 22.55 | 8.26 | -1.63 | 0.00773949 |
| 4.46 | 4.34 | 1.42 | 0.19718447 |
| 926.94 | 176.92 | -1.17 | 0.10787214 |
| 27.32 | 9.08 | 1.5 | 0.01921528 |
| 20.67 | 9.9 | -1.38 | 0.27785838 |

| | | | |
|---|---|---|---|
| 14.87 | 8.87 | 2.42 | 0.08221931 |
| 188.44 | 49.92 | 1.06 | 0.75635689 |
| 6.63 | 4.98 | -1.24 | 0.41358668 |
| 436.31 | 198.43 | -1.78 | 0.07737803 |
| 19.06 | 11.02 | -1.56 | 0.05368978 |
| 295.06 | 75.85 | -1.32 | 0.05571619 |
| 30.43 | 20.56 | -1.05 | 0.85615897 |
| 3.1 | 1.04 | 1.72 | 0.00830848 |
| 426.95 | 144.13 | -1.81 | 0.01134775 |
| 83.95 | 29.16 | 1 | 0.9827618 |
| 3.18 | 1.13 | 1.91 | 0.08251287 |
| 3.47 | 3.15 | 1.68 | 0.04336975 |
| 3.74 | 4.15 | 1.1 | 0.7876206 |
| 21.72 | 19.53 | -1.57 | 0.22705777 |
| 6.33 | 4.09 | -1.42 | 0.3113417 |
| 57.49 | 18.98 | -1.53 | 0.05541013 |
| 168.79 | 57.97 | 1.6 | 0.00776686 |
| 325.64 | 73.51 | -1.32 | 0.01236382 |
| 17.93 | 9.67 | -1.5 | 0.26451951 |
| 405.24 | 125.99 | -1.5 | 0.02565927 |
| 189.81 | 60.21 | -1.47 | 0.03211385 |
| 3.02 | 1.01 | 2.27 | 0.00530256 |
| 175.49 | 33.45 | -1.11 | 0.3326124 |
| 65.61 | 18.49 | -1.32 | 0.09350619 |
| 155.23 | 71.29 | -1.48 | 0.20665805 |
| 3.53 | 1.73 | 1.69 | 0.01064456 |
| 95.39 | 34.53 | -1.26 | 0.44515041 |
| 225.39 | 85.5 | -1.26 | 0.35178006 |
| 97.93 | 24.55 | -1.6 | 0.03538283 |
| 9.22 | 6.62 | 1.62 | 0.09666194 |
| 7.37 | 9.72 | 2.34 | 0.06204626 |
| 71.13 | 63.13 | -1.09 | 0.76366234 |
| 10.46 | 7.56 | 1.73 | 0.24036913 |
| 2.99 | 1.01 | 1.37 | 0.28734329 |
| 90.83 | 41.99 | -1.27 | 0.34418482 |
| 2.99 | 1.01 | 1.37 | 0.28734329 |
| 1515.03 | 560.62 | -1.66 | 0.07656691 |
| 58.66 | 10.42 | 1.02 | 0.91700053 |
| 3.74 | 2.45 | 1.64 | 0.04268837 |
| 22.68 | 10.97 | -1.37 | 0.24574819 |
| 22.01 | 10.07 | -1.18 | 0.39130828 |
| 2.99 | 1.01 | 7.59 | 0.00142977 |
| 3.68 | 2.59 | 1.48 | 0.09690243 |
| 35.3 | 24.26 | -1.78 | 0.15591796 |
| 3.72 | 3.15 | 1.5 | 0.25107449 |
| 31.35 | 14.73 | -1.6 | 0.05576848 |
| 3.18 | 1.13 | 1.29 | 0.38933089 |

| | | | |
|---|---|---|---|
| 5.49 | 7.2 | 20.36 | 0.00481105 |
| 6.57 | 7.55 | 1.78 | 0.27489719 |
| 5.13 | 4.55 | 1.42 | 0.27077022 |
| 3.6 | 2.5 | 5.32 | 0.00557791 |
| 3.1 | 1.04 | 4.64 | 0.00372046 |
| 53.09 | 27.9 | -1.89 | 0.03908902 |
| 229.51 | 35.73 | -1.18 | 0.11034594 |
| 5.14 | 4.32 | -1.06 | 0.85456258 |
| 2.99 | 1.01 | 1.37 | 0.28734329 |
| 4.58 | 4.18 | 1.35 | 0.38833424 |
| 3.87 | 3.35 | 5.51 | 0.0024123 |
| 2.99 | 1.01 | 1.62 | 0.07585831 |
| 2.99 | 1.01 | 1.37 | 0.28734329 |
| 4.11 | 4.81 | 1.26 | 0.50900191 |
| 16.28 | 15.27 | -1.64 | 0.32677492 |
| 42.7 | 19.85 | -1.02 | 0.93434584 |
| 3.66 | 4.96 | 1.86 | 0.04546095 |
| 8.45 | 6.29 | -1.26 | 0.45923609 |
| 5.12 | 4.65 | 1.06 | 0.85957003 |
| 4.35 | 4 | 1.04 | 0.90521842 |
| 2.99 | 1.01 | 1.37 | 0.28734329 |
| 6.27 | 9.86 | -1.02 | 0.96763998 |
| 14.21 | 9.34 | -1.05 | 0.89324021 |
| 3.64 | 3.41 | 1.39 | 0.29127076 |
| 363.77 | 123.81 | -1.35 | 0.19400427 |
| 5.92 | 4.53 | 1.18 | 0.56193912 |
| 13.62 | 6.93 | -1.2 | 0.50960159 |
| 4.41 | 4.95 | 1.6 | 0.19403116 |
| 21.9 | 17.72 | -2.05 | 0.21216868 |
| 9.32 | 9.98 | -1.12 | 0.77875543 |
| 22.04 | 19.06 | -1.14 | 0.73078686 |
| 211.06 | 74.41 | -1.31 | 0.1959272 |
| 15.81 | 11.71 | -1.6 | 0.16649239 |
| 4.97 | 4.57 | 2.3 | 0.01314915 |
| 2.99 | 1.01 | 1.37 | 0.28734329 |
| 3.6 | 2.5 | 1.37 | 0.25862369 |
| 27.61 | 7.82 | -1.25 | 0.0525741 |
| 23.46 | 8.26 | -1.22 | 0.32102886 |
| 22.92 | 15.06 | -1.2 | 0.59159744 |
| 48.85 | 31.12 | 3.14 | 0.08594662 |
| 13.95 | 11.59 | -1.93 | 0.02444627 |
| 2.99 | 1.01 | 1.74 | 0.05554172 |
| 15.44 | 9.2 | 2.17 | 0.05063996 |
| 2.99 | 1.01 | 3.16 | 0.00038037 |
| 13.55 | 9.56 | -1.47 | 0.15248907 |
| 16.34 | 9.49 | -1.71 | 0.13887148 |
| 91.76 | 27.94 | -1.59 | 0.04094965 |

| | | | |
|---|---|---|---|
| 36.61 | 8.6 | -1.55 | 0.00871335 |
| 86.12 | 14.88 | -1.09 | 0.26283973 |
| 3.72 | 3.25 | 1.53 | 0.25213623 |
| 29.54 | 10.3 | -1.79 | 0.06410123 |
| 10.37 | 10.92 | -1.06 | 0.87814873 |
| 7.87 | 7.79 | -1.15 | 0.70074803 |
| 108.9 | 51.26 | -2.49 | 0.00033941 |
| 2.99 | 1.01 | 2.33 | 0.00146559 |
| 229.46 | 92.23 | 1.07 | 0.76105827 |
| 22.09 | 6.16 | -1.29 | 0.16355889 |
| 309.92 | 67.32 | -1.32 | 0.08238699 |
| 19.31 | 15.5 | -1.58 | 0.16455819 |
| 1241.44 | 168.07 | -1.3 | 0.00272397 |
| 102.72 | 15.83 | -1.19 | 0.04770104 |
| 153.14 | 60.86 | -1.76 | 0.07348598 |
| 18.72 | 11.7 | -1.59 | 0.20865229 |
| 130.44 | 44.1 | -1.52 | 0.01152228 |
| 1565.79 | 306.94 | -1.29 | 0.0425602 |
| 33.29 | 10.58 | 3.73 | 0.00416429 |
| 2.99 | 1.01 | 1.37 | 0.28734329 |
| 45.41 | 14.78 | -1.57 | 0.01974993 |
| 148.98 | 47.8 | -1.4 | 0.13751829 |
| 20.41 | 7 | 1.06 | 0.72276312 |
| 35.82 | 16.62 | -1.71 | 0.03342335 |
| 4.44 | 3.78 | 1.18 | 0.58937919 |
| 94.37 | 43.81 | 1.37 | 0.22421061 |
| 75.86 | 28.83 | -1.68 | 0.04466571 |
| 5.99 | 5.36 | -1.05 | 0.87249404 |
| 10.42 | 10.39 | -1.05 | 0.8903234 |
| 45.76 | 11.81 | -1.3 | 0.12255397 |
| 3.5 | 1.75 | 1.29 | 0.34316432 |
| 4.28 | 3.94 | 1.57 | 0.12018437 |
| 91.98 | 21.74 | 1.1 | 0.35724887 |
| 112.61 | 32.91 | -1.12 | 0.54648155 |
| 3.02 | 1.01 | 1.77 | 0.00579485 |
| 69.09 | 33.08 | -2.08 | 0.0254882 |
| 71.4 | 30.94 | -1.75 | 0.02767722 |
| 2.99 | 1.01 | 27.19 | 0.00325066 |
| 3.81 | 4.28 | 2.48 | 0.00718903 |
| 5.77 | 4.7 | 1.06 | 0.86265743 |
| 54.42 | 25.57 | -1.79 | 0.01466803 |
| 18.25 | 2.84 | 1.38 | 0.03239597 |
| 4.69 | 6.19 | 1.9 | 0.14158145 |
| 11.19 | 9.66 | -1 | 0.99560076 |
| 55 | 15.8 | -1.18 | 0.4530279 |
| 21.77 | 12.42 | -1.58 | 0.19163893 |
| 40.55 | 32.36 | 1.59 | 0.29919648 |

| | | | |
|---|---|---|---|
| 62.03 | 18.46 | -1.15 | 0.44438219 |
| 142.94 | 37.84 | -1.37 | 0.03450433 |
| 362.4 | 127.32 | -1.71 | 0.03289973 |
| 5.48 | 6.25 | 1.04 | 0.89616776 |
| 2.99 | 1.01 | 1.62 | 0.06929228 |
| 7.56 | 4.88 | -1.13 | 0.5970633 |
| 21.62 | 8.87 | -1.43 | 0.17414567 |
| 57.52 | 24.91 | -1.69 | 0.1017197 |
| 3.6 | 2.5 | 1.14 | 0.67962903 |
| 94.16 | 27.49 | -1.35 | 0.0430436 |
| 4.49 | 4.94 | 1.02 | 0.94454187 |
| 13.62 | 7.72 | 1.04 | 0.86562037 |
| 290.65 | 66.95 | -1.37 | 0.04127354 |
| 3.29 | 1.47 | 1.25 | 0.46738774 |
| 60.59 | 11.94 | -1.14 | 0.19676155 |
| 18.32 | 13.37 | -1.77 | 0.13286884 |
| 94.23 | 39.22 | -1.38 | 0.24889888 |
| 42.29 | 20.98 | -1.27 | 0.39533055 |
| 14.43 | 9.64 | -1.55 | 0.16616198 |
| 41.76 | 14.56 | 1.39 | 0.26658654 |
| 157.55 | 23.72 | -1.23 | 0.17573796 |
| 591.65 | 103.81 | -1.17 | 0.15043563 |
| 20.66 | 8.64 | -1.52 | 0.04594143 |
| 11.14 | 8.17 | 1.26 | 0.53661782 |
| 33.67 | 15.17 | -2.23 | 0.00070845 |
| 9.89 | 5.57 | -1.01 | 0.97104889 |
| 2.99 | 1.01 | 1.74 | 0.05554172 |
| 133.5 | 58.37 | -1.53 | 0.15539314 |
| 2.99 | 1.01 | 1.37 | 0.28734329 |
| 27.11 | 8.5 | 1.19 | 0.54476994 |
| 4.66 | 3.76 | 1.04 | 0.89962184 |
| 40.01 | 18.73 | -3.29 | 0.00375692 |
| 89.04 | 39.56 | -1.88 | 0.03005389 |
| 45.9 | 22.67 | -1.02 | 0.94091129 |
| 60.56 | 12.96 | -1.03 | 0.87352151 |
| 3.69 | 2.98 | 1.31 | 0.35586885 |
| 47.68 | 19.34 | 1.01 | 0.96189332 |
| 40.37 | 17.96 | -1.06 | 0.75126129 |
| 188.6 | 53.06 | -1.14 | 0.44112405 |
| 73.42 | 25.11 | -1.99 | 0.00045878 |
| 490.49 | 192.09 | -1.69 | 0.02435867 |
| 6.22 | 4.29 | -1.07 | 0.8172273 |
| 13 | 7.53 | -1.34 | 0.30220956 |
| 3.51 | 3.45 | 1.59 | 0.1949579 |
| 25.44 | 12.93 | -1.89 | 0.04047694 |
| 9.18 | 7.45 | 1.16 | 0.63229752 |
| 3.53 | 4.16 | 1.28 | 0.44035849 |

| | | | |
|---|---|---|---|
| 11.36 | 9.22 | 1.04 | 0.8920595 |
| 6.12 | 8.53 | 2.25 | 0.04418526 |
| 14.06 | 9.51 | 1.55 | 0.22505775 |
| 17.82 | 11.75 | 3.18 | 0.01206678 |
| 4.09 | 4.62 | 2.36 | 0.0206138 |
| 3.57 | 3.98 | 1.86 | 0.04294425 |
| 4.11 | 2.35 | 1.6 | 0.06569796 |
| 22.75 | 7.98 | -1.55 | 0.03462807 |
| 5.11 | 4.71 | 1.1 | 0.78649336 |
| 25.49 | 10.49 | -1.2 | 0.31635585 |
| 3.63 | 1.7 | 1.33 | 0.2712388 |
| 243.64 | 92.58 | -1.52 | 0.21855067 |
| 8.84 | 10.22 | 1.42 | 0.38304767 |
| 29.17 | 15.52 | 1.76 | 0.2592133 |
| 8.75 | 6.39 | 1.17 | 0.55137646 |
| 13.68 | 10.35 | 2.14 | 0.07228857 |
| 5.42 | 5.6 | -1.06 | 0.88540417 |
| 2.99 | 1.01 | 1.79 | 0.00521556 |
| 138.29 | 22.76 | -1.5 | 0.00637698 |
| 176.94 | 46.24 | -1.29 | 0.06388915 |
| 55.28 | 16.82 | 1.2 | 0.36030382 |
| 18.33 | 11.19 | 1.33 | 0.37450394 |
| 32.6 | 18.14 | -1.71 | 0.02117152 |
| 38.96 | 11.13 | 1.03 | 0.80114496 |
| 13.23 | 4.88 | 7.58 | 0.01310592 |
| 28 | 9.64 | 1.27 | 0.17415074 |
| 49.7 | 19.19 | -1.68 | 0.05964108 |
| 1056.8 | 429.42 | -1.62 | 0.08638515 |
| 89.55 | 16.42 | -1.24 | 0.08018779 |
| 5.93 | 5.01 | 1.26 | 0.47599924 |
| 7.32 | 5.63 | -1.02 | 0.95334584 |
| 23.52 | 4.94 | -2.98 | 0.0059666 |
| 52.16 | 17.72 | -1.43 | 0.01603295 |
| 3.85 | 2.5 | 3.56 | 0.00295087 |
| 15.85 | 8.74 | -2.55 | 0.01157638 |
| 147.66 | 48.04 | -1.07 | 0.68527955 |
| 412.59 | 153.25 | -1.47 | 0.1828884 |
| 123.09 | 58.4 | 1.48 | 0.11769144 |
| 10.74 | 2.64 | -1.28 | 0.53199732 |
| 139.64 | 77.64 | -1.88 | 0.07819135 |
| 7.81 | 10.52 | 1.39 | 0.42697403 |
| 82.23 | 12.34 | -1.12 | 0.32013455 |
| 39.46 | 13.99 | 1.31 | 0.08237634 |
| 166.89 | 49.34 | -1.4 | 0.01636427 |
| 21.41 | 6.67 | -1.48 | 0.20820875 |
| 3.6 | 2.5 | 1.44 | 0.21802616 |
| 3.02 | 1.01 | 1.5 | 0.13214917 |

| | | | |
|---|---|---|---|
| 4.74 | 4.02 | 1.62 | 0.12131612 |
| 115.05 | 37.03 | -1.46 | 0.08511765 |
| 117.89 | 38.25 | -1.42 | 0.0737566 |
| 3.62 | 2.55 | 2.06 | 0.02100035 |
| 31.6 | 4.77 | -1.19 | 0.08157232 |
| 172.91 | 39.4 | -1.38 | 0.04738636 |
| 22.41 | 6.61 | -1.1 | 0.54631209 |
| 37.23 | 11.76 | 1.01 | 0.96561307 |
| 2.99 | 1.01 | 1.7 | 0.06360078 |
| 19.06 | 11.9 | -2.15 | 0.00545628 |
| 219.95 | 29.52 | 1.25 | 0.00384811 |
| 3.93 | 2.53 | 1.42 | 0.23972067 |
| 47.45 | 14.3 | -1.14 | 0.34536463 |
| 6.07 | 5.94 | -1.34 | 0.40405223 |
| 217.74 | 67.66 | -1.47 | 0.01600087 |
| 4.85 | 5.57 | 2.46 | 0.02478828 |
| 76.46 | 20.69 | -1.14 | 0.1664452 |
| 13.52 | 9.21 | -1.62 | 0.0949203 |
| 114.93 | 34.63 | -1.31 | 0.11512998 |
| 4.58 | 4.16 | 1.99 | 0.02511007 |
| 18.48 | 11.14 | -2.38 | 0.00671863 |
| 162.11 | 75.04 | -1.38 | 0.12096477 |
| 35.75 | 25.25 | 1.44 | 0.37152985 |
| 29.41 | 7.6 | -1.06 | 0.59395874 |
| 31.33 | 23.78 | -2.21 | 0.03520769 |
| 6.74 | 6.4 | 2.09 | 0.04828129 |
| 8.31 | 6.05 | 3.27 | 0.00355144 |
| 37.92 | 18.18 | 1.88 | 0.15329801 |
| 39.88 | 20.01 | 4.8 | 0.00840233 |
| 88.23 | 40.1 | -1.59 | 0.11524753 |
| 46.11 | 23.44 | -1.22 | 0.51055044 |
| 11.28 | 7.57 | -1.44 | 0.25666788 |
| 27.9 | 12.61 | -1.48 | 0.0357406 |
| 40.66 | 19.3 | -1.79 | 0.07430424 |
| 17.25 | 11.41 | -1.34 | 0.26543027 |
| 4.35 | 4.74 | 2.77 | 0.01819726 |
| 58.51 | 19.69 | -1.31 | 0.19900134 |
| 3.02 | 1.01 | 1.94 | 0.03455006 |
| 41.12 | 21.23 | -1.79 | 0.08209962 |
| 16.23 | 8.91 | -1.21 | 0.34829739 |
| 7.33 | 3.7 | -1.26 | 0.48098823 |
| 6.91 | 6.12 | 1.71 | 0.212754 |
| 3.85 | 4.26 | 2.39 | 0.01083465 |
| 4.53 | 3.92 | 5.2 | 0.0002599 |
| 3.39 | 1.91 | 1.58 | 0.04483341 |
| 60.57 | 14.33 | -1.04 | 0.73520482 |
| 6.24 | 6.54 | 1.17 | 0.64160222 |

| | | | |
|---|---|---|---|
| 3.18 | 1.13 | 5.05 | 0.00624328 |
| 3.5 | 1.75 | 1.53 | 0.04719256 |
| 7 | 7.44 | 2.12 | 0.06326801 |
| 6.66 | 6.26 | 1.01 | 0.98042315 |
| 3.29 | 1.47 | 1.57 | 0.1135198 |
| 4.76 | 3.81 | 3.23 | 0.00103921 |
| 5.08 | 4.75 | 1.65 | 0.20210437 |
| 549.68 | 185.89 | -1.48 | 0.08070388 |
| 26.55 | 8.61 | -1.2 | 0.29495132 |
| 28.97 | 10.58 | 1.18 | 0.51187122 |
| 3.24 | 1.28 | 3.47 | 0.00035516 |
| 27.59 | 24.47 | -1.61 | 0.28794423 |
| 8.76 | 6.69 | 5.45 | 0.00130126 |
| 31.71 | 13.26 | 1.29 | 0.16889046 |
| 13.92 | 12.97 | 1.41 | 0.38868815 |
| 3.73 | 2.13 | 1.54 | 0.19149062 |
| 3.5 | 1.75 | 1.41 | 0.20381901 |
| 7.92 | 9.75 | 4.46 | 0.02096403 |
| 5.83 | 7.24 | 1.27 | 0.48197106 |
| 4.95 | 4.86 | 1.45 | 0.22445084 |
| 70.11 | 30.12 | -1.58 | 0.05349176 |
| 17.04 | 8.37 | -1.36 | 0.27822831 |
| 14.38 | 9.54 | -1.5 | 0.17810921 |
| 49.11 | 9.63 | 1.03 | 0.83632988 |
| 2.99 | 1.01 | 1.37 | 0.28734329 |
| 143.59 | 59.51 | -1.97 | 0.00444897 |
| 14.34 | 6.18 | -1.51 | 0.03318491 |
| 146.42 | 43 | -1.36 | 0.03580935 |
| 57.5 | 18.64 | -1.7 | 0.01580232 |
| 68.49 | 20.02 | -1.11 | 0.60915047 |
| 64.14 | 20.99 | -1.45 | 0.09570836 |
| 103.64 | 35.93 | -1.35 | 0.10280432 |
| 11.52 | 1.11 | -1.33 | 0.00225751 |
| 502.91 | 129.98 | -1.42 | 0.01171311 |
| 9.89 | 5.25 | -1.32 | 0.26069513 |
| 141.22 | 46.09 | -1.45 | 0.13676436 |
| 15.89 | 7.39 | -1.11 | 0.60548568 |
| 138.91 | 62.34 | -1.91 | 0.07878516 |
| 12.79 | 11.29 | -1.29 | 0.42518538 |
| 37.82 | 10.48 | 1.24 | 0.10426039 |
| 2.99 | 1.01 | 1.37 | 0.28734329 |
| 9.56 | 8.36 | 1.86 | 0.08179407 |
| 72.35 | 33.46 | -1.75 | 0.06961989 |
| 44.17 | 11.86 | -1.36 | 0.0273276 |
| 8.67 | 8.94 | -1.45 | 0.35864177 |
| 94.05 | 33.45 | -1.25 | 0.23366736 |
| 93.48 | 26.42 | -1.37 | 0.13585623 |

| | | | |
|---|---|---|---|
| 136.19 | 71.16 | -1.39 | 0.22684416 |
| 15.76 | 30.54 | 1.42 | 0.61355197 |
| 14.14 | 6.24 | -1.01 | 0.96661317 |
| 5.34 | 5.17 | -1.1 | 0.77421194 |
| 3.53 | 4.16 | 1.98 | 0.02888225 |
| 2.99 | 1.01 | 1.37 | 0.28734329 |
| 13.73 | 9.09 | 1.13 | 0.65586376 |
| 4.81 | 4.64 | 1.21 | 0.62375551 |
| 6.07 | 4.38 | 1.33 | 0.39334804 |
| 14.69 | 12.77 | -1.58 | 0.2664088 |
| 93.8 | 24.34 | -1.05 | 0.73252261 |
| 112.97 | 50.72 | -1.63 | 0.07781339 |
| 16.89 | 12.62 | -1.09 | 0.79135615 |
| 13.68 | 9.31 | -1.72 | 0.07656512 |
| 19.43 | 5.4 | -1.73 | 0.0012655 |
| 942.34 | 271.64 | -1.4 | 0.05103927 |
| 361.59 | 72.76 | -1.1 | 0.36762103 |
| 11.79 | 6.51 | 42.2 | 0.00782955 |
| 256.84 | 72.5 | -1.46 | 0.16625224 |
| 53.36 | 26.8 | -1.8 | 0.05654676 |
| 86.31 | 19.3 | -1.21 | 0.25833508 |
| 254.75 | 22.76 | -1.06 | 0.20055558 |
| 23.43 | 6.52 | -1.49 | 0.1355304 |
| 10.11 | 5.79 | 2 | 0.12585595 |
| 78.33 | 23.88 | -1.48 | 0.01651471 |
| 67.33 | 12.43 | -1.19 | 0.17930759 |
| 15.51 | 4.65 | -1.36 | 0.09458036 |
| 3.43 | 2 | 1.56 | 0.15772906 |
| 5.25 | 6.05 | 1.27 | 0.41945532 |
| 15.05 | 3.18 | -1.39 | 0.14974096 |
| 26.32 | 9.92 | -2.22 | 0.01003553 |
| 5.15 | 4.83 | 1.24 | 0.46002525 |
| 11.22 | 8.85 | -1.52 | 0.24336484 |
| 3.24 | 1.28 | 1.4 | 0.21590203 |
| 5.68 | 5.2 | 1.66 | 0.28061229 |
| 5.75 | 3.95 | 1.12 | 0.64266956 |
| 132.11 | 46.41 | 1.51 | 0.19475862 |
| 3.78 | 3.09 | 1.09 | 0.79702848 |
| 8.62 | 10.42 | -1.22 | 0.58711982 |
| 12.08 | 10.88 | -1.03 | 0.91828227 |
| 28.16 | 19.52 | -1.06 | 0.83548641 |
| 5.14 | 4.24 | 1.92 | 0.07473692 |
| 174.51 | 31.02 | -1.22 | 0.01798845 |
| 662.44 | 259.43 | -1.59 | 0.05339642 |
| 94.57 | 31.63 | -1.82 | 0.0214416 |
| 206.88 | 45.13 | -1.33 | 0.06192416 |
| 224.25 | 38.81 | -1.02 | 0.72827053 |

| | | | |
|---|---|---|---|
| 387.47 | 89.26 | -1.26 | 0.08138707 |
| 21.81 | 10.33 | -1.15 | 0.62629586 |
| 3.2 | 1.21 | 1.68 | 0.01301075 |
| 493.79 | 95.18 | -1.3 | 0.03083068 |
| 3.79 | 1.79 | 1.22 | 0.4698984 |
| 44.85 | 18.77 | -1.17 | 0.49889851 |
| 249.02 | 42.41 | -1.24 | 0.03121255 |
| 3.02 | 1.01 | 1.98 | 0.00059689 |
| 29 | 7.73 | -1.28 | 0.12699749 |
| 3.66 | 4.96 | 1.56 | 0.13777906 |
| 165.55 | 17.99 | -1.02 | 0.74934012 |
| 117.32 | 34.67 | -1.45 | 0.03347369 |
| 7.3 | 5.11 | 1.02 | 0.93774474 |
| 14.44 | 13.71 | -1.36 | 0.47099584 |
| 7.75 | 3.83 | 1.36 | 0.16075975 |
| 11.06 | 5.54 | -1.07 | 0.85976607 |
| 130.94 | 49.75 | -1.47 | 0.11366665 |
| 34.28 | 16.64 | -1.43 | 0.0869645 |
| 36.25 | 13.97 | -1.19 | 0.4201301 |
| 36.03 | 13.24 | -1.8 | 0.06961004 |
| 11.52 | 9.85 | -1.32 | 0.49973452 |
| 2.99 | 1.01 | 2.12 | 0.00964117 |
| 13.3 | 5.3 | -1.39 | 0.15053302 |
| 1794.94 | 274.95 | -1.16 | 0.06997365 |
| 115.79 | 22.06 | 1 | 0.98883492 |
| 32.14 | 11.67 | -1.07 | 0.61004883 |
| 4.68 | 6.76 | 1.84 | 0.14579912 |
| 21.03 | 15.2 | -1.76 | 0.15584862 |
| 29.73 | 7.69 | -1.06 | 0.67098194 |
| 3.48 | 3.02 | 1.39 | 0.2620481 |
| 3.02 | 1.01 | 1.72 | 0.0592312 |
| 5.51 | 5.38 | 2.08 | 0.06996965 |
| 80.54 | 20.67 | -1.26 | 0.05356257 |
| 19.34 | 17.02 | 1.07 | 0.85787904 |
| 6.44 | 7.22 | 1.74 | 0.19960417 |
| 36.52 | 14.43 | 1 | 0.9973371 |
| 8.92 | 5.24 | 1.95 | 0.1002411 |
| 40.74 | 47.4 | -2.83 | 0.17627575 |
| 8.99 | 5.85 | 1.82 | 0.05719628 |
| 186.17 | 36.47 | -1.02 | 0.88480932 |
| 7.73 | 6.06 | 1.74 | 0.13422652 |
| 5.29 | 4.64 | 1.38 | 0.37557808 |
| 18.41 | 15.64 | -1.13 | 0.77573037 |
| 101.52 | 26.23 | -1.14 | 0.20106035 |
| 171.5 | 63.09 | -1.57 | 0.08975883 |
| 2.99 | 1.01 | 1.37 | 0.28734329 |
| 437.65 | 133.15 | -1.24 | 0.10987692 |

| | | | |
|---|---|---|---|
| 17.42 | 13.93 | -1.36 | 0.42414579 |
| 59.31 | 21.21 | -1.39 | 0.1500522 |
| 9.64 | 8.75 | 1.64 | 0.27775812 |
| 14.72 | 10.78 | 1.99 | 0.06393314 |
| 5.15 | 3.67 | 1.11 | 0.74384624 |
| 9.44 | 11.7 | -1.15 | 0.75659341 |
| 3.34 | 1.86 | 1.89 | 0.07333359 |
| 4 | 5.57 | 1.36 | 0.37218994 |
| 5.28 | 4.18 | -1.29 | 0.45857957 |
| 123.67 | 25.97 | -1.05 | 0.62651354 |
| 281.6 | 79.25 | -1.34 | 0.13507609 |
| 4.62 | 2.78 | 2.25 | 0.01445919 |
| 18.12 | 18.14 | -1.61 | 0.26444149 |
| 12.43 | 7.86 | -1.36 | 0.31690526 |
| 9.01 | 6.7 | 1.27 | 0.47909018 |
| 18.58 | 8.68 | 1.05 | 0.78907746 |
| 118.94 | 27.04 | -1.25 | 0.22306387 |
| 30.3 | 16.33 | -1.42 | 0.10660472 |
| 17.04 | 36.91 | -1.28 | 0.54680115 |
| 3.77 | 2.5 | 1.52 | 0.08184908 |
| 25.85 | 17.38 | -1.33 | 0.30936584 |
| 10.56 | 3.79 | -1.17 | 0.49032435 |
| 105.5 | 49.9 | -1.89 | 0.08607908 |
| 3.5 | 1.75 | 2.45 | 0.00286319 |
| 16.79 | 10.37 | -1.47 | 0.18548329 |
| 36.26 | 14.32 | -1.73 | 0.0575106 |
| 163.75 | 38.36 | -2.1 | 0.00057151 |
| 126.57 | 45.87 | -1.51 | 0.05601991 |
| 5 | 5.2 | 1.11 | 0.78010613 |
| 18.6 | 16.38 | 3.56 | 0.03423291 |
| 13.09 | 7.53 | 1.22 | 0.45750624 |
| 58.25 | 27.44 | -4.79 | 0.00168162 |
| 508 | 106.02 | -1.39 | 0.04608776 |
| 49.4 | 15.67 | -1.25 | 0.21962595 |
| 664.46 | 179.09 | -1.36 | 0.03349033 |
| 73.18 | 19.09 | -1.51 | 0.01866801 |
| 46.95 | 20.74 | -1.64 | 0.10688028 |
| 114.39 | 20.18 | -1.07 | 0.34282318 |
| 41.38 | 10.8 | 1.39 | 0.03770161 |
| 36.64 | 11.84 | -1.59 | 0.00825098 |
| 85.14 | 18.17 | -1.24 | 0.08476967 |
| 56.71 | 22.39 | 1.25 | 0.18009725 |
| 98.32 | 26.22 | -1.28 | 0.10064647 |
| 26.95 | 12.97 | -1.44 | 0.2037591 |
| 3.18 | 1.13 | 1.55 | 0.09392657 |
| 282.58 | 126.48 | -1.94 | 0.03452758 |
| 140.66 | 39.21 | -1.47 | 0.11285275 |

| | | | |
|---|---|---|---|
| 2.99 | 1.01 | 1.52 | 0.12365852 |
| 6.69 | 6.2 | 1.08 | 0.81151122 |
| 3.48 | 3.02 | 1.18 | 0.6139099 |
| 857.33 | 114.32 | -1.19 | 0.01091398 |
| 15.89 | 6.04 | -1.35 | 0.11325884 |
| 9.94 | 11.9 | -1.6 | 0.33191574 |
| 15.19 | 3.94 | -1.44 | 0.02099073 |
| 39.87 | 27.55 | -1.44 | 0.36634436 |
| 14.75 | 13.12 | -1.51 | 0.21756226 |
| 2115.13 | 467.42 | -1.19 | 0.09197887 |
| 15.24 | 10.11 | 1.21 | 0.51716399 |
| 37.82 | 17.29 | -1.56 | 0.04712934 |
| 2999.02 | 401.05 | -1.04 | 0.68876499 |
| 114.24 | 24.94 | -1.3 | 0.02164877 |
| 54.26 | 23.75 | -1.95 | 0.01087761 |
| 105.87 | 56.53 | -1.7 | 0.06086886 |
| 51.8 | 14.6 | -1.15 | 0.31894743 |
| 10.12 | 4.38 | 1.08 | 0.70251167 |
| 6.41 | 6.81 | 1.11 | 0.74744761 |
| 42.34 | 9.75 | -1.02 | 0.8892374 |
| 268.96 | 40.26 | -1.02 | 0.83541614 |
| 243.59 | 42.95 | -1.28 | 0.09511073 |
| 288.15 | 98.79 | -1.38 | 0.12854366 |
| 6.37 | 4.9 | 1.21 | 0.54857153 |
| 30.89 | 7.89 | -1.2 | 0.23817609 |
| 2.99 | 1.01 | 1.37 | 0.28734329 |
| 676.26 | 100.99 | -1.27 | 0.00718493 |
| 386.19 | 44.13 | -1.24 | 0.11567003 |
| 952.61 | 177.81 | -1.3 | 0.029492 |
| 208.55 | 58.09 | -1.49 | 0.05255706 |
| 172.03 | 53.88 | -1.62 | 0.0530236 |
| 94.56 | 23.58 | 1.06 | 0.69888073 |
| 32.28 | 12.57 | -1.86 | 0.00306385 |
| 26.01 | 10.45 | -1.27 | 0.16934144 |
| 25.98 | 11.5 | -1.03 | 0.87985229 |
| 40.6 | 18.32 | -1.74 | 0.06357543 |
| 17.83 | 10.09 | 2.09 | 0.05125511 |
| 133.72 | 31.89 | 1.09 | 0.51455873 |
| 7.42 | 6.82 | 1.05 | 0.8916977 |
| 3.51 | 2.17 | 1.5 | 0.18578523 |
| 3.24 | 1.28 | 8.29 | 0.00201626 |
| 260.24 | 46.74 | -1.18 | 0.2175083 |
| 7.93 | 7.28 | -1.03 | 0.93969798 |
| 4.76 | 3.43 | 1.55 | 0.15378739 |
| 154.98 | 17.3 | -1.08 | 0.40397623 |
| 5.33 | 4.4 | 3.31 | 0.01916143 |
| 2.99 | 1.01 | 1.37 | 0.28734329 |

| | | | |
|---|---|---|---|
| 577.55 | 119.54 | -1.27 | 0.02592904 |
| 757.47 | 175.01 | -1.37 | 0.03413778 |
| 409.02 | 46.75 | -1.1 | 0.07361653 |
| 4.76 | 4.15 | 1.1 | 0.79664797 |
| 34.45 | 11.38 | -1.27 | 0.17915829 |
| 20.78 | 9.05 | -1.19 | 0.45263177 |
| 4.2 | 3.09 | 1.66 | 0.07937127 |
| 7.93 | 4.98 | -1.1 | 0.71076089 |
| 9.85 | 9.92 | -1.1 | 0.8317467 |
| 9.85 | 10.43 | -1.46 | 0.32608598 |
| 17.24 | 9.99 | -1.2 | 0.40716767 |
| 2.99 | 1.01 | 1.7 | 0.06360078 |
| 104.95 | 20.97 | -1.22 | 0.04308254 |
| 46.14 | 8.52 | 1.16 | 0.23501559 |
| 87.95 | 22.64 | 1 | 0.98063427 |
| 176.52 | 41.93 | -1.27 | 0.07389738 |
| 336.69 | 123.67 | -1.6 | 0.04073847 |
| 37.06 | 18.94 | -1.25 | 0.47480094 |
| 12.41 | 7.75 | 1.56 | 0.168029 |
| 18.92 | 7.43 | 1.01 | 0.97060341 |
| 36.9 | 19.49 | -1.36 | 0.18955377 |
| 206.42 | 25.01 | -1.11 | 0.1346031 |
| 26.77 | 19.86 | -1.42 | 0.43311796 |
| 6.75 | 5.83 | 1.39 | 0.4473221 |
| 46.86 | 14.12 | -1.25 | 0.08288417 |
| 152.15 | 48.46 | -1.32 | 0.06687719 |
| 627.42 | 136.43 | -1.36 | 0.01149863 |
| 414.68 | 113.34 | -1.31 | 0.03779472 |
| 95.52 | 40.3 | 1.19 | 0.35236016 |
| 983.71 | 287.23 | -1.43 | 0.05470353 |
| 74.37 | 33.31 | 1.14 | 0.5339821 |
| 680.87 | 163.41 | -1.28 | 0.15144302 |
| 2084.03 | 742.29 | -1.58 | 0.05978039 |
| 55.36 | 23.87 | -1.6 | 0.10106902 |
| 749.95 | 186.81 | -1.27 | 0.06328908 |
| 11.6 | 7.93 | -1.51 | 0.23417155 |
| 20.2 | 15.05 | 5.75 | 0.00660679 |
| 9.76 | 14.51 | -1.08 | 0.86243296 |
| 120.98 | 40.71 | -1.06 | 0.70345366 |
| 441.47 | 119.5 | -1.06 | 0.63193542 |
| 9.08 | 7.81 | 1.49 | 0.26146159 |
| 4.98 | 4.14 | 1.16 | 0.61300844 |
| 32.8 | 14.88 | -1.79 | 0.02712292 |
| 69.5 | 34.99 | -1.87 | 0.02270013 |
| 3.18 | 1.13 | 2.05 | 0.00113698 |
| 20.51 | 11.47 | 2.25 | 0.06381987 |
| 3.62 | 2.55 | 1.14 | 0.67261201 |

| | | | |
|---|---|---|---|
| 78.59 | 26.68 | -1.77 | 0.00242574 |
| 17.89 | 8.36 | -1.01 | 0.94910687 |
| 87.11 | 30.72 | -1.23 | 0.42175418 |
| 18.76 | 7.79 | -1.29 | 0.23291664 |
| 35.56 | 15.98 | 1.04 | 0.82524937 |
| 53.27 | 15.18 | -1.31 | 0.17361793 |
| 117.6 | 29.74 | -1.04 | 0.76661134 |
| 64.75 | 23.33 | -1.06 | 0.69793427 |
| 28.6 | 9.9 | -1.51 | 0.0153298 |
| 5.42 | 6.06 | -1.03 | 0.92756087 |
| 2.99 | 1.01 | 1.37 | 0.28734329 |
| 25.51 | 7.87 | 1.19 | 0.28456908 |
| 3.65 | 2.12 | 1.7 | 0.02108178 |
| 6.76 | 8.25 | -1.03 | 0.94721645 |
| 77.51 | 26.48 | -1.29 | 0.20372862 |
| 53.93 | 29.99 | -1.48 | 0.20767789 |
| 3.51 | 2.17 | 1.29 | 0.37043098 |
| 4.95 | 4.57 | -1.02 | 0.9418242 |
| 6.57 | 9.98 | 1.32 | 0.49661431 |
| 425.11 | 85.1 | -1.29 | 0.02929863 |
| 104.44 | 84.05 | -1.98 | 0.14200711 |
| 423.81 | 75.63 | -1.19 | 0.19895919 |
| 72.54 | 21.21 | -1.4 | 0.03320432 |
| 8.73 | 5.08 | -1.46 | 0.27575234 |
| 246.62 | 55.53 | -1.06 | 0.52649635 |
| 317.39 | 88.3 | 1.06 | 0.69403613 |
| 12.93 | 3.74 | -1.18 | 0.44059655 |
| 6.32 | 4.37 | 2.8 | 0.0254626 |
| 127.34 | 10.38 | -1.04 | 0.51453835 |
| 274.5 | 29.14 | -1.15 | 0.02125214 |
| 18.25 | 9.87 | -1.29 | 0.23962142 |
| 34.24 | 17.22 | 1.31 | 0.35020283 |
| 57.1 | 12.98 | -1.04 | 0.80210733 |
| 2.99 | 1.01 | 2.76 | 0.00321676 |
| 445.53 | 65.62 | -1.14 | 0.25286564 |
| 36.18 | 6.87 | -1.83 | 0.01465393 |
| 21.88 | 6.24 | -1.77 | 0.06615817 |
| 81.44 | 43.81 | -1.13 | 0.72066349 |
| 2.99 | 1.01 | 1.37 | 0.28734329 |
| 3.29 | 1.47 | 1.25 | 0.46738774 |
| 4.28 | 4.25 | 1.33 | 0.29568207 |
| 311.2 | 147.95 | -1.23 | 0.47607774 |
| 667.11 | 99.21 | -1.26 | 0.04313948 |
| 45.19 | 30.19 | 1.73 | 0.22535676 |
| 153.36 | 18.75 | 1.06 | 0.49392459 |
| 44.94 | 21.64 | -1.52 | 0.07832357 |
| 30.4 | 19.72 | -1.41 | 0.25472414 |

| | | | |
|---|---|---|---|
| 6.64 | 9.91 | 4.94 | 0.00121278 |
| 3.02 | 1.01 | 7.4 | 0.00192927 |
| 1659.57 | 483.52 | -1.55 | 0.03684263 |
| 24.8 | 10.37 | -1.17 | 0.43500999 |
| 520.11 | 165.49 | -1.48 | 0.09798847 |
| 34.33 | 11.46 | -1.22 | 0.2334757 |
| 20.73 | 11.98 | -1.67 | 0.12916619 |
| 181.55 | 68.63 | -1.49 | 0.17094842 |
| 10.01 | 5.98 | -1.17 | 0.64470828 |
| 383.73 | 74.97 | -1.31 | 0.11540733 |
| 440.28 | 116.36 | -1.05 | 0.63619721 |
| 29.17 | 17.78 | -1.8 | 0.02789412 |
| 21.73 | 7.22 | -1.07 | 0.6008473 |
| 6.35 | 9.79 | -1.31 | 0.50392115 |
| 6.8 | 8.95 | 1.08 | 0.8429563 |
| 45.3 | 28.54 | -1.48 | 0.27113965 |
| 3.51 | 2.17 | 1.17 | 0.61764014 |
| 54.7 | 13.47 | -1.27 | 0.08371731 |
| 364.36 | 119.22 | -1.48 | 0.04907729 |
| 28.96 | 10.5 | 1.12 | 0.45526654 |
| 4.23 | 5.48 | 1.92 | 0.0532273 |
| 81.28 | 30.82 | -1.78 | 0.01249634 |
| 2.99 | 1.01 | 1.39 | 0.27767301 |
| 18.3 | 7.87 | 2.65 | 0.00622522 |
| 12.87 | 7.48 | -1.91 | 0.13175264 |
| 14.18 | 4.22 | 1.2 | 0.37694222 |
| 2.99 | 1.01 | 4.91 | 0.00509477 |
| 40.48 | 24.98 | 1.3 | 0.36844957 |
| 7.4 | 8.24 | 1.99 | 0.17536539 |
| 4.61 | 3.76 | 1.34 | 0.26373097 |
| 3.98 | 5.14 | 1.24 | 0.52433026 |
| 55.19 | 21.49 | 1.37 | 0.31174043 |
| 43.09 | 12.61 | -1.53 | 0.00689755 |
| 136.35 | 61.89 | -1.8 | 0.03534091 |
| 22.24 | 7.92 | -1.06 | 0.78539306 |
| 59.4 | 30.93 | -1.99 | 0.08313469 |
| 20.6 | 7.61 | 1.1 | 0.58510858 |
| 4.47 | 8.23 | 1.56 | 0.20528643 |
| 21.51 | 14.16 | -1.89 | 0.09435233 |
| 64.53 | 25.78 | -1.78 | 0.04363366 |
| 3.18 | 1.13 | 1.6 | 0.09653381 |
| 185.99 | 91.46 | -2.03 | 0.03444638 |
| 95.94 | 27.14 | -1.38 | 0.02146216 |
| 3.1 | 1.04 | 1.33 | 0.34384 |
| 7.82 | 4.99 | 1.06 | 0.87088722 |
| 11.78 | 7.86 | -2.27 | 0.03174499 |
| 57.58 | 26.73 | -1.32 | 0.23049615 |

| | | | |
|---|---|---|---|
| 4.62 | 7.51 | 1.35 | 0.41884488 |
| 8.11 | 5.91 | -1.18 | 0.57496417 |
| 12.22 | 7.62 | 2.88 | 0.01297409 |
| 5.16 | 6.29 | 1.72 | 0.1306861 |
| 4.3 | 5.06 | 1.21 | 0.58375376 |
| 3.02 | 1.01 | 1.36 | 0.30205497 |
| 8.2 | 9.99 | -1.04 | 0.91313839 |
| 22.56 | 5.74 | -1.94 | 0.00010377 |
| 9.39 | 6.64 | -1.22 | 0.52204758 |
| 4.02 | 3.26 | 1.33 | 0.2655313 |
| 2.99 | 1.01 | 1.62 | 0.07585831 |
| 14.92 | 14.52 | -1.7 | 0.23601787 |
| 28.63 | 16.39 | -1.33 | 0.26933676 |
| 261.52 | 59.78 | -1.21 | 0.26825801 |
| 12.38 | 9.75 | -1.62 | 0.19396649 |
| 13.4 | 9.06 | -1.44 | 0.34593132 |
| 4.91 | 6.4 | 1.35 | 0.41853428 |
| 38.13 | 16.04 | -1.4 | 0.27083632 |
| 71.38 | 24.48 | -1.3 | 0.25191998 |
| 32.34 | 20.14 | -1.58 | 0.13765307 |
| 4.33 | 4.14 | 1.17 | 0.63274795 |
| 5.34 | 5.3 | 1.2 | 0.51845872 |
| 21.33 | 10.18 | -2.26 | 0.00120341 |
| 4.55 | 4.32 | -1.11 | 0.76953912 |
| 7.19 | 6.36 | 1.17 | 0.62498897 |
| 18.78 | 11.4 | -1.38 | 0.19990863 |
| 47.28 | 12.78 | -1.26 | 0.09296513 |
| 5.5 | 5.45 | 1.04 | 0.89633864 |
| 3.55 | 2.15 | 1.5 | 0.07122439 |
| 13.47 | 5.56 | -1.43 | 0.23075569 |
| 6.09 | 6.68 | 1.1 | 0.76042104 |
| 23.73 | 128.55 | -2.84 | 0.11579856 |
| 803.03 | 146.3 | -1.21 | 0.0435745 |
| 179.04 | 66.67 | 1.2 | 0.3289119 |
| 307.92 | 111.74 | -1.42 | 0.14106506 |
| 16 | 12.57 | 2.11 | 0.05659803 |
| 488.56 | 208.07 | -2.07 | 0.00317624 |
| 55.73 | 15.33 | 1.03 | 0.80491161 |
| 9.72 | 5.5 | -1.04 | 0.8782078 |
| 17.22 | 17.01 | 1.08 | 0.87468708 |
| 236.56 | 61.39 | -1.33 | 0.05939501 |
| 7.23 | 5.37 | 1.41 | 0.22892256 |
| 69.59 | 15.26 | -1.58 | 0.00494886 |
| 49.16 | 16.7 | -1.2 | 0.27590352 |
| 109.54 | 13.36 | 1 | 0.93863243 |
| 57.29 | 20.61 | -1.23 | 0.17406356 |
| 3.97 | 3.63 | 1.14 | 0.68541414 |

| | | | |
|---|---|---|---|
| 26.19 | 14.74 | -1.18 | 0.52579474 |
| 6.09 | 4.39 | 1.19 | 0.56506342 |
| 207.44 | 20.51 | -1.06 | 0.36096793 |
| 21.22 | 2.45 | 1.08 | 0.45644274 |
| 3.47 | 3.15 | 6.12 | 0.00212711 |
| 65.19 | 19.57 | -1.46 | 0.01320375 |
| 66.07 | 25.96 | 1.35 | 0.19254541 |
| 2.99 | 1.01 | 1.37 | 0.28734329 |
| 116.75 | 17.75 | -1.05 | 0.59786248 |
| 54.48 | 10.37 | 1.12 | 0.3888402 |
| 83.55 | 27.73 | -1.61 | 0.00822526 |
| 50.42 | 17.09 | -1.24 | 0.36620522 |
| 13.39 | 3.79 | -1.21 | 0.22428872 |
| 48.13 | 11.56 | 1.15 | 0.17257784 |
| 21.32 | 8.64 | 1.07 | 0.77899158 |
| 37.94 | 10.64 | -1.35 | 0.08453921 |
| 107.43 | 38.59 | -1.27 | 0.2988112 |
| 35.17 | 14.43 | -1.16 | 0.53545028 |
| 76.13 | 9.42 | -1.12 | 0.1911495 |
